# Oxidized MIF is an Alzheimer’s Disease drug target relaying external risk factors to tau pathology

**DOI:** 10.1101/2021.09.11.459903

**Authors:** Andreas Müller-Schiffmann, Felix Torres, Anatolly Kitaygorodskyy, Anand Ramani, Argyro Alatza, Sarah K. Tschirner, Ingrid Prikulis, Shaofeng Yu, Debendranath Dey, Suguna Mallesh, Dharma Prasad, Dennis Solas, Verian Bader, Annemieke Rozemuller, Selina Wray, Jay Gopalakrishnan, Roland Riek, Vishwanath R. Lingappa, Carsten Korth

## Abstract

The viral life cycle usurps host cellular factors, redirecting them from physiological functions to viral needs thereby revealing their “moonlighting” functions, disturbing cellular proteostasis, and increasing risk of specific, virus-associated protein misfolding diseases (PMD). Identifying such virus-repurposed host proteins therefore allow study of fundamental cellular events leading to associated “sporadic” PMD. Here, we identified a small molecule with unprecedented activity against neurotropic herpes simplex virus 1 (HSV-1) modulating an allosteric site of Macrophage Migration Inhibitory Factor (MIF). The compound efficiently reduced HSV-1-mediated tau phosphorylation or aggregation *in vitro* and *in vivo*, even without HSV-1 infection. The lead compound specifically interacted with an oxidized conformer of MIF (oxMIF) from either recombinant MIF or *post-mortem* brain homogenates of patients with Alzheimer’s disease (AD). OxMIF thus participates in a host-viral interface connecting HSV-1 infection, and possibly other external stressors, with tau cellular pathology characteristic for PMD, including Alzheime’s disease.

## Introduction

Despite an abundance of epidemiological evidence highlighting external risk factors critically contributing to the emergence of sporadic neurodegenerative diseases, the exact molecular links of how these factors are relayed to the well-characterized neuropathological or key cellular and molecular features of neurodegenerative diseases have remained unknown. Among those risk factors are such diverse conditions as obesity, diabetes, trauma, chronic infections, inflammation or immune activations (Arnold et al., 2018; Itzhaki et al., 2020). Although known not to cause neurodegenerative diseases directly, the subchronic exposure to neurotropic viruses has been associated to the occurrence to, for example, Parkinson’s disease (PD) for influenza virus (Marreiros et al., 2020), Alzheimer’s disease (AD) for herpes viruses (Itzhaki et al., 1997), or others (Levine et al., 2023). Either direct effects of the virus on host factors also involved in critical molecular mechanisms of neurodegeneration or indirect effects of the virus through immune activation or its elicited cellular responses, are conceivable.

Virus propagation involves recruitment of host cell proteins for replication and assembly of its components. This occurs through a series of discrete steps and leads to a disruption of proteostasis, that eventually can result in protein misfolding and, depending on the cellular context, protein aggregates (Müller-Schiffmann et al., 2021). The identification of particular host proteins diverted by a virus and also involved in balancing protein aggregation, is thus a strategy to identify pathophysiological mechanisms of protein aggregation. These host-viral molecular interfaces may have a more general significance beyond virus-specific effects, e.g., as general converging hubs or relays for external stressors.

The discovery of novel antiviral drugs has so far primarily focused on virus-encoded proteins, which makes intuitive sense, since the virus is the causative agent of virus-induced disease. Viral replication *in vivo*, however, is dependent on cellular host factors, which therefore provide an alternative target, albeit with the challenge of avoiding host toxicity. Virus dependence on host factors is not limited to nucleic acid replication but includes capsid assembly, which is catalyzed in cells by virus-recruited host proteins, perhaps exploiting their moonlighting functions (Jeffery, 2020), and can be reconstituted in cell-free protein synthesis and assembly (CFPSA) systems (Lingappa et al., 2013a; Lingappa et al., 2013b; Reed et al., 2021). The establishment of CFPSA assays enables novel drug screens (Lingappa et al., 2013b). Drug-targeting of these cellular host factors has several advantages: First, drug resistance development is greatly diminished since the selection of host factors is on a much longer time scale than viral replication cycles. Second, the effects caused by host factor recruitment can give clues as to the basis for virus-induced cellular pathology, which ultimately causes clinical disease (Müller-Schiffmann et al., 2021). Third, the ability to target a small subpopulation of specific host proteins recruited for their moonlighting functions could, in principle, not affect their canonical functions in cellular homeostasis. Bundling these advantages are challenging to obtain by rational design. However, they can be gleaned from the viruses who achieved those innovations by natural selection over deep evolutionary time (Müller-Schiffmann et al., 2021).

Herpes simplex virus (HSV-1) is a human pathogenic virus, which in rare cases causes overt encephalitis but endemically stays latent in sensory neurons causing reactivation upon a variety of conditions. Reactivation of latent herpes virus infection has been associated to Alzheimer’s disease (AD) even though the molecular mechanisms of this connection remains unknown (Eimer et al., 2018; Itzhaki et al., 1997; Lovheim et al., 2015; Wozniak et al., 2009). Recently, a positive association of HSV-1 antibodies to increased levels of phosphorylated tau in CSF of AD patients was demonstrated (Goldhardt et al., 2022).

We here present the discovery of a drug (PAV-174) highly active against HSV-1 in human brain organoids and human neuronal cell lines with a clear structure-activity relationship (SAR). By NMR spectrometry we demonstrate that the lead compound directly targets and partially reverses a distinct oxidized conformation of macrophage migration inhibitory factor (MIF). We validate MIF as a critical host factor by CRISPR/Cas9 knockout experiments and demonstrate its role in tau molecular pathology relevant for neurodegenerative diseases such as AD or the frontotemporal dementias (FTD). PAV-174 reduced HSV-1 induced tau phosphorylation but was also active in the absence of infection *in vitro* and *in vivo*. A surrogate of the oxidized conformation of MIF (oxMIF) is capable of directly inducing tau phosphorylation and we detected increased amounts of oxMIF in brain tissue of AD patients and after infection of differentiated neurons with HSV-1. Thus, oxMIF appears to be a missing molecular link connecting HSV-1 infection, and possibly other risk factors, with cellular pathology characteristic for neurodegenerative diseases involving aberrant tau phosphorylation or aggregation.

## Results

### PAV-174 prevents HSV-1 infection in differentiated LUHMES cells and human brain organoids

From a cell-free capsid assembly assay screen of a chemical library (Lingappa et al., 2010; Lingappa et al., 2013b) similar to what we have successfully described for rabies virus (Lingappa et al., 2013a) and human immunodeficiency virus (HIV) (Reed et al., 2021), an early lead compound (PAV-645) was identified with activity in inhibiting herpes virus capsid assembly. This compound was subsequently optimized for activity over seven generations of analog synthesis and screening, to an improved compound PAV-174 that inhibited HSV-1 replication in a dose-dependent manner in HSV-1-infected Vero cells, both in a standard plaque assay (**Figure 1A**) and in an in-cell ELISA (**Figure 1B**) with an EC_50_ of 34 nM or 21 nM, respectively, and an CC_50_ of 1.2 µM (Table 1, **Supplementary Figure 1B**). The compound was also active in differentiated primary neuron-like human dopaminergic LUnd Human MESencephalic (LUHMES) cells (**Figure 1C, D**), as well as in differentiated 60 d human brain organoids (**Figure 1E, F**) where it also lowered HSV-1-induced neurotoxicity (**Figure 1G**). This indicated clear antiherpetic activity in several independent systems, including human primary-like cells. The compound also displayed a clear SAR as measured by an in-cell ELISA in Vero cells (**Table 1, Supplementary Figure 1**).

**Figure 1.**
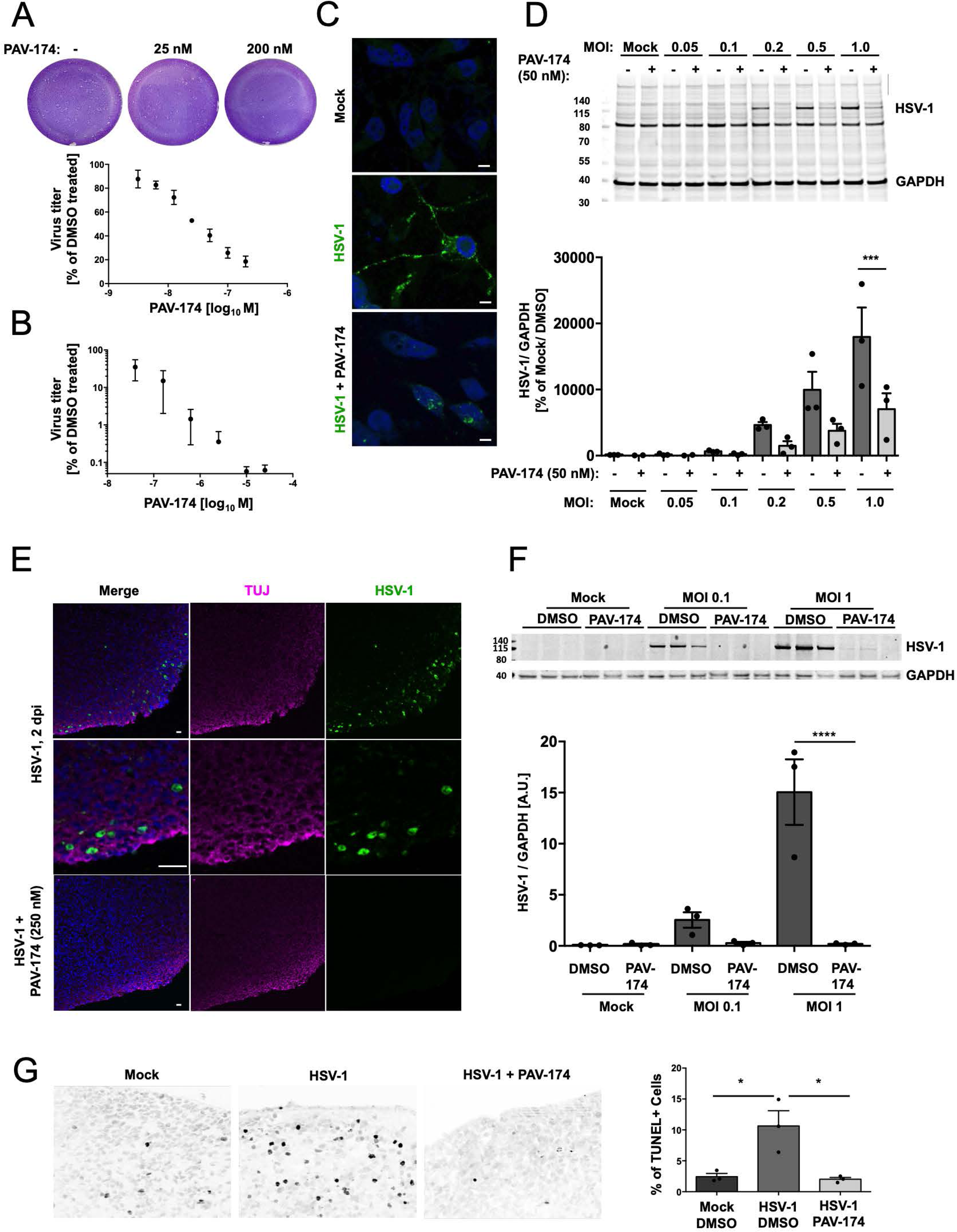
PAV-174 prevents HSV-1 infection in differentiated LUHMES and human brain organoids. (A) The antiviral activity of PAV-174 was determined by standard plaque assay displaying infective viral units from the lysates of infected Vero cells (MOI = 1) that were treated with PAV-174 between 6 nM and 200 nM. Each data point was normalized to the negative control and displays the mean +/− SEM of three experiments (n=3). The IC_50_ of PAV-174 was calculated to be 34 nM using the logarithm of inhibitor concentration on GraphPad Prism 6.0. (B) The results from (A) were validated by an in-Cell ELISA, measuring HSV-1-gC in Vero cells treated with increasing concentrations of PAV-174 and infected with HSV-1 (MOI = 1). Each data point was normalized to the DMSO negative control and displays the mean +/− SEM of three independent experiments (n=3). Here, the IC_50_ of PAV-174 was calculated to be 21 nM using the logarithm of inhibitor concentration on GraphPad Prism 6.0. (C) ICC of differentiated LUHMES cells infected with HSV-1 (MOI = 1) for 24h either treated with DMSO or 50 nM of PAV-174. The cells were stained with an antibody against HSV-1-gC (green). Cell nuclei were stained with DAPI. Viral antigens were detected within the nucleus and cytoplasm as well as in the axons of non-treated cells. PAV-174 markedly reduced the presence of viral antigens. Only within the nucleus signals could be detected. (bar = 10 µm) (D) Western Blot of cell lysates derived from differentiated LUHMES cells that were infected with increasing MOI of HSV-1 (0.05 to 1.0) and treated with DMSO or 50 nM of PAV-174. The upper signal represents an HSV-1 antigen detected with ab9533. GAPDH was used as internal control. The diagram below shows the HSV-1 signals normalized to GAPDH from three (n=3) independent experiments. Data were analyzed by Two-way ANOVA (Sidak’s post-hoc). (E) Staining of 60d human brain organoids that were infected with HSV-1 (MOI = 1) for 48h and treated with DMSO or 250 nM of PAV-174. HSV-1-gC (green) infected TUJ-1-positive neurons (magenta) in the outer layer of the organoids. Infection with HSV-1 was completely abolished when PAV-174 was present (lower panel). For the reason of contrast the signals for HSV-1-gC detected with the Alexa-fluor-594 antibody is displayed in green. (bars = 30 µm) (F) Western Blot with lysates from 60d human brain organoids infected with HSV-1 (MOI of 0.1 or 1.0). HSV-1 signals were normalized with GAPDH. The diagram below represents the results from three infected organoids per group. Data were analyzed by Two-way ANOVA (Sidak’s post-hoc). (G) PAV-174 (250 nM) lowered cell toxicity mediated by HSV-1 infection. Compared to the mock condition, HSV-1 infected organoids displayed increased TUNEL-positive cells, which was reverted by addition of PAV-174. For each condition three different organoids (n=3) and at least 10 images of every organoid were analyzed. Data were analyzed by One-way ANOVA (Tukeýs post hoc). See also Figure S1.

**Table 1.**
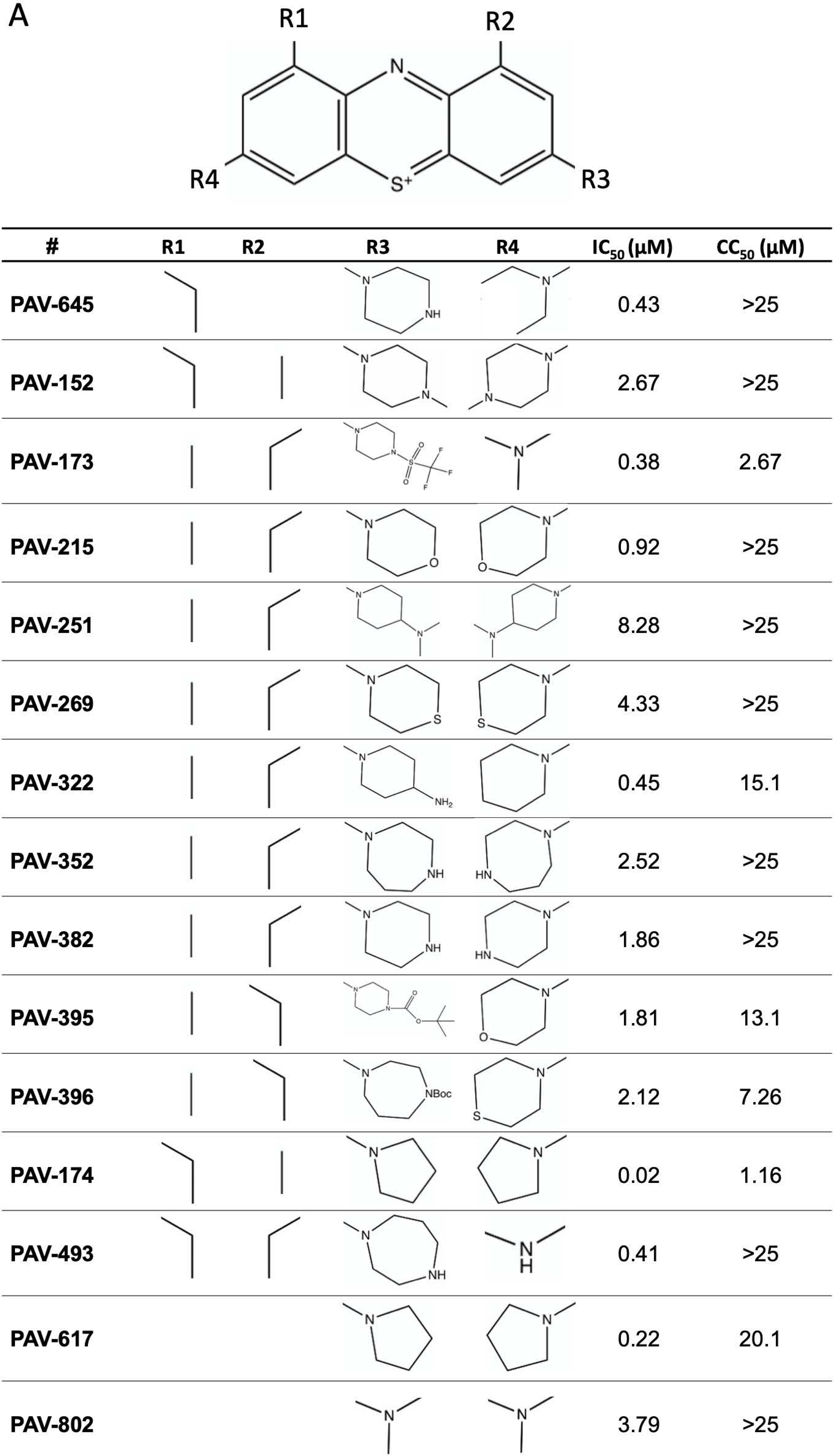
Structures of tested compounds including IC_50_ and CC_50_ values derived by in-Cell ELISA and MTT assays, respectively. See also Figure S1.

### PAV-174 binds MIF at its multimerization interface

In order to determine the cellular target of PAV-174, we performed drug resin-affinity chromatography (DRAC) with immobilized parent compound PAV-645. We used pig brain lysates (by availability) as material and, after washing, the final binding proteins were eluted with an excess of free PAV-645. The eluate was analyzed by tandem mass spectrometry. Several proteins were identified, some of them having been described in the context of HSV-1- or AD-related research (**Supplementary Figure 2A**): macrophage migration inhibitory factor (MIF; (Petralia et al., 2020)), copper binding protein (cutA; (Hou et al., 2015), glutathione-S-transferase Pi1 (GSTP1; (Wang, 2015)), peroxiredoxin 2 (PRDX2; (Szeliga, 2020)) and calmodulin-like 3 (CALML3; (Chen et al., 2019)). Even though these results suggested that multiple proteins, or a multiprotein complex, could be ligands of active compound, we focused on MIF because of its known role in infection and neurodegenerative disorders (Park et al., 2022; Petralia et al., 2020; Ruan et al., 2021; Shvil et al., 2018). We further validated these findings by DRAC of MIF in SH-SY5Y neuroblastoma cells (**Supplementary Figure 2B**) as well as in human brains (see below). In order to confirm binding of PAV-174 to MIF we measured ^15^N-Heteronuclear Single Quantum Coherence (HSQC) spectra of PAV-174 with wildtype MIF recombinantly expressed and purified from *E. coli* (**Supplementary Figure 2C**). Spectra revealed discrete binding sites (**Supplementary Figure 2D**) suggesting specific binding. An NMR structure of recombinant MIF together with compound PAV-174 demonstrated a binding site at the interface of MIF monomers forming a trimer (**Figure 2A, B**). The MIF-PAV-174 complex structure revealed a ligand pose in the aromatic hinge formed by the residues Tyr36, Ile64, Val106, Trp108, and Phe113 at the interface between the different monomers of MIF. PAV-174 is in particular inserting in the hinge by forming two π-π interactions, one with Trp108 from a monomeric subunit, and the second with Tyr36 from a different monomeric subunit. The long tricyclic aromatic core moiety of PAV-174 engages in interactions with a more important number of residues than it is typically observed from other molecules known to bind at the same binding pocket, such as ISO-1 or AV1013 (Bloom et al., 2016), Indeed, ISO-1 inserts deeper in the binding pocket, at the location of the embedded pyrrolidine in beta of the ethyl aliphatic chain (**Figure 2C**). The ethyl aliphatic locates in the same region as the isopropyl chain of the AV1013, suggesting an importance of a C2/C3 aliphatic substitution at this location to establish hydrophobic interactions. Finally, the part of PAV-174 that points out of the pocket is engaging into π-π interaction with Trp108, this interaction seems to be missing for ISO-1 and AV1013. ISO-1 misses also the aliphatic C2/C3 chain that is observed to collocate for AV1013 and PAV-174.

**Figure 2.**
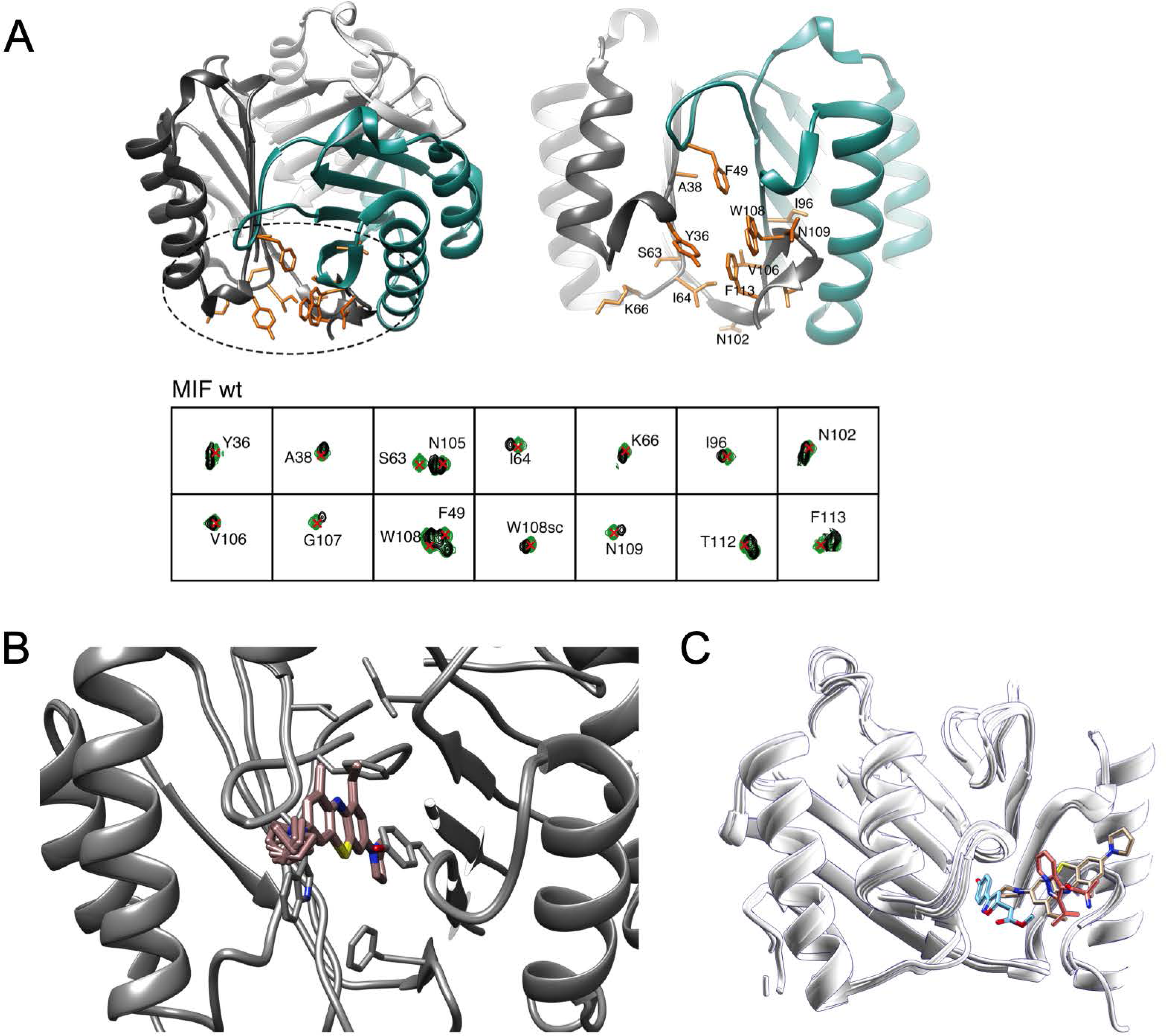
PAV-174 binds MIF at the multimerization interface. (A) Structure of the MIF trimer interface with the residues showing compound PAV-174-induced chemical shift perturbations in the ^15^N-HSQC spectra (Supplementary Figure 2D) represented in orange sticks. Each monomer is represented in a different color, i.e., cyan and grey. (B) Structure of the compound PAV-174 in complex with MIF calculated with the *N*MR^2^ protocol using ligand to protein’s aromatics and ligand to protein’s methyls distance restraints. The 10 best calculated poses have been retained, i.e., the poses with the lowest target function. (C) Overlay of binding poses of PAV-174 (beige, PDB ID: tbd), ISO-1 (blue, PDB ID: 1LJT), and AV1013 (orange, PDB ID: 3IJG) on MIF (Bloom et al., 2016). The structures were overlaid using the Chimera software for visualization. The structure overlay reveals a longer insertion into the binding pocket of PAV-174 in comparison to ISO-1 which inserts deeper in the binding pocket, and to AV1013 which inserts at the surface of the binding pocket. See also Figure S2.

### PAV-174 reduces tau phosphorylation also in the absence of infection in a MIF- and dose-dependent manner

In order to analyze effects of PAV-174 in a cellular model for neurodegenerative diseases we generated a neuroblastoma-derived cell line stably overexpressing human tau (2N4R) including the familial mutation P301S (SH-SY5Y-tau-P301S). These cells could efficiently be infected with HSV-1 **(Supplementary Figure 3A)**. HSV-1 infection increased tau phosphorylation at residues Ser202, Thr205 (epitopes of monoclonal antibody AT8) in SH-SY5Y-tau-P301S cells (**Figure 3A**). Compound PAV-174 reduced tau phosphorylation at these sites (**Figure 3A**), but also in absence of virus in a dose-dependent manner at low nanomolar concentrations (**Figure 3B**). A dose dependent reduction of phosphorylation was also found at T231 (antibody AT180) but only at higher concentrations on T181 (antibody AT270) indicating a specific effect (**Supplementary Figure 3B**). The tau phosphorylation-inhibiting effects displayed a SAR (**Figure 3C; Supplementary Figure 3C**) that positively correlated with the anti-HSV-1 activity (**Figure 3D**). Next, we intended to confirm the functional relevance of MIF for PAV-174. The effect of PAV-174 on inhibiting tau phosphorylation was MIF dependent. This was demonstrated in SH-SY5Y-tau-P301S that were knocked down for MIF by using the CRISPR/Cas9 technique (**Supplementary Figure 3D**). In those cells PAV-174 effects on tau phosphorylation were absent (**Figure 3E**). Of note, PAV-174 did not act like other well-studied MIF inhibitors such as ISO-1 (Bacher et al., 2010; Li et al., 2015) which does not modulate tau phosphorylation even at high µM concentrations (**Figure 3F).** MIF modulates a range of different interconnected signaling cascades, including ERK1/2 MAP, PI3K/Akt and JNK, via binding to the chemokine receptors CXCR2, CXCR4 and CD74 (Leng et al., 2003; Lue H et al., 2011; Schwartz et al., 2009). PAV-174 dose-dependently increased phosphorylation of Akt at Ser473 as well as GSK3β at Ser9 in a dose-dependent and MIF dependent manner, since the effects were significantly less pronounced in SH-tauP301S-MIF-ko cells (**Supplementary Figure 4A/B**). We conclude that the effects of PAV-174 on MIF dependent tau phosphorylation are executed via Akt and GSK3β.

**Figure 3.**
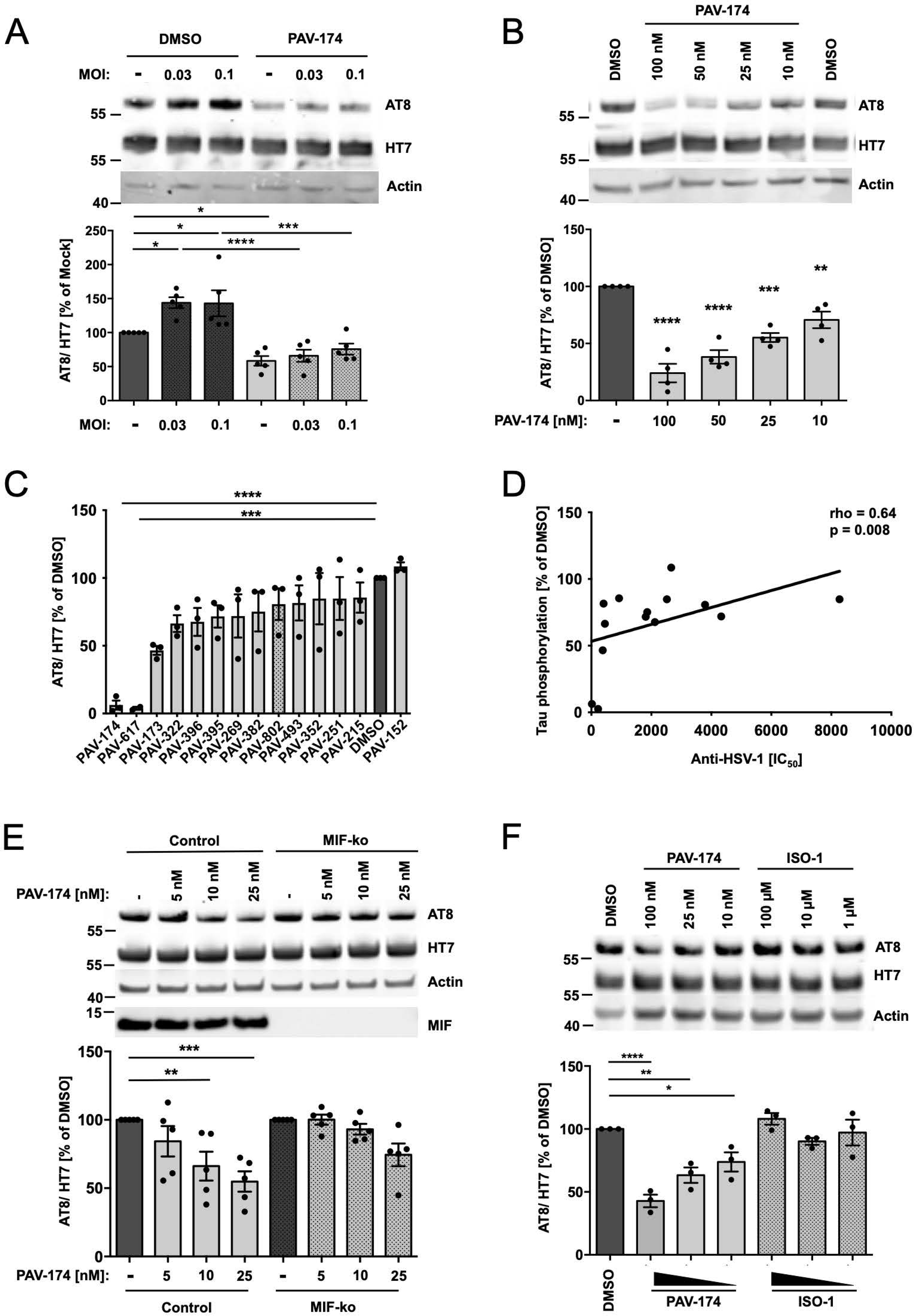
PAV-174 reduces tau phosphorylation also in the absence of infection in a MIF and dose dependent manner. (A) Infection of SH-SY5Y-tau-P301S cells with low MOI of HSV-1 (0.03 and 0.1) increased tau phosphorylation (AT8) 6h p.i. This increase was absent in the presence of PAV-174 (20 nM). The diagram shows the AT8 signals normalized to total tau (HT7) from five (n=5) independent experiments. Staining of actin served as loading control. Data were analyzed by Two-way ANOVA (Sidak’s post-hoc). (B) PAV-174 dose dependently reduced tau phosphorylation at the AT8 site at low nanomolar concentrations also in the absence of viral infection. SH-SY5Y-tau-P301S cells were treated with PAV-174 for 48h. Already 10 nM of PAV-174 were sufficient to significantly reduce tau phosphorylation. Actin served as loading control. The values from four independent experiments (n=4) are presented and were analyzed by One-way ANOVA (Dunnett’s post-hoc). (C) SAR of PAV-174 analogs on tau phosphorylation. SH-SY5Y-tau-P301S cells were treated with 500 nM of compounds for 48h. The diagram shows the AT8 signals normalized to total tau from three independent experiments (n=3). Data were analyzed by One-way ANOVA (Dunnett’s post-hoc). (D) Correlation analysis of the PAV-174 analogs between reduction of tau phosphorylation (% of DMSO) and inhibition of HSV-1 activity (IC_50_). The effects of the compounds on tau phosphorylation positively correlated to their capacity to inhibit HSV-1 replication. Spearmańs rho coefficient and p value are indicated in the graph. (E) Reduction of tau phosphorylation (AT8) by PAV-174 was impaired in SH-SY5Y-tau-P301S-MIF-knockout cells. SH-SY5Y-tau-P301S-control and -MIF-ko cells treated with indicated concentrations of PAV-174. The diagram shows the average of HT7 normalized AT8 values as % of DMSO treated cells derived from five independent experiments (n=5). Actin served as loading control and the absence of MIF was validated with a polyclonal antibody against MIF. Data were analyzed by Two-way ANOVA (Tuckeýs post-hoc). (F) The MIF inhibitor ISO-1 does not modulate tau phosphorylation. SH-SY5Y-tau-P301S cells were treated with indicated amounts of PAV-174 or ISO-1. Even at high concentrations of 100 µM ISO-1 did not reduce tau phosphorylation as detected by AT8 normalized to total tau (HT7). The diagram shows the average values of three independent experiments (n=3). Data were analyzed by One-way ANOVA (Dunnett’s post-hoc). See also Figures S3, S4.

Of note, HSV-1 capsid antigen accumulated faster in the MIF knockdown cell line with increasing multiplicity of infection (MOI). This effect was most pronounced at an MOI of 0.2 (**Supplementary Figure 4C**), whereas infectivity as measured in the plaque assay was not changed, thereby increasing the HSV-1 antigen/titer ratio and corroborating our hypothesis that HSV-1 capsid assembly inhibition leads to an accumulation of non-assembled viral proteins. Full activity of lead compound PAV-174 was also dependent on the presence of MIF. In MIF knockout SH-SY5Y-tau-P301S cells, lead compound PAV-174 did not exert comparable titer-lowering effects (**Supplementary Figure 4D**) indicating that MIF likely, at least in part, mediates the antiherpetic effects of PAV-174.

### PAV-174 decreases tau phosphorylation and aggregation in vivo

To corroborate the effects of lead compound PAV-174 on tau phosphorylation in primary-like human cells, we treated cortical neurons differentiated from human iPSCs of a patient carrying the MAPT IVS10+16 mutation (V97) (Sposito et al., 2015) with PAV-174 for 48h. We observed reduced phosphorylation at position Ser396 and Ser404 of tau that are also targeted by GSK3β (Augustinack et al., 2002), detected by specific antibody PHF-1, and at the AT8 site (**Figure 4A**). In order to validate the effects of PAV-174 *in vivo*, we treated two months old tgTau58/2 mice (van Eersel et al., 2015), (overexpressing human tau with the P301S mutation) with 5 mg/kg i. p. of the structural analog PAV-617 (see Table 1) three times weekly for 4 weeks. We chose PAV-617 due to its improved pharmacokinetic properties. PAV-617 was less toxic in mice and showed a substantially higher bioavailability likely due lower protein binding of 91.2% compared to 99,9% for PAV-174 (**Supplementary Figure 5A**). Both compounds had comparable activity regarding inhibition of tau phosphorylation (**Figure 3C**). PAV-617 showed a high bioavailability following intra peritoneal administration and good tissue penetration including the brain (**Supplementary Figure 5B/C**). We found significantly reduced phosphorylation of tau (PHF-1 and AT8) in brain homogenates of PAV-617-treated mice (**Figure 4B; Supplementary Figure 5D/E**). Furthermore, sarkosyl insoluble tau species after ultracentrifugation were significantly less abundant in brain tissue of treated mice (**Figure 4C; Supplementary Figure 5E**) indicating less tau aggregation in addition to less tau phosphorylation after PAV-174 treatment. Together these observations confirm the potency of these compounds in reducing tau phosphorylation and aggregation in disease-relevant models *in vitro* and *in vivo*.

**Figure 4.**
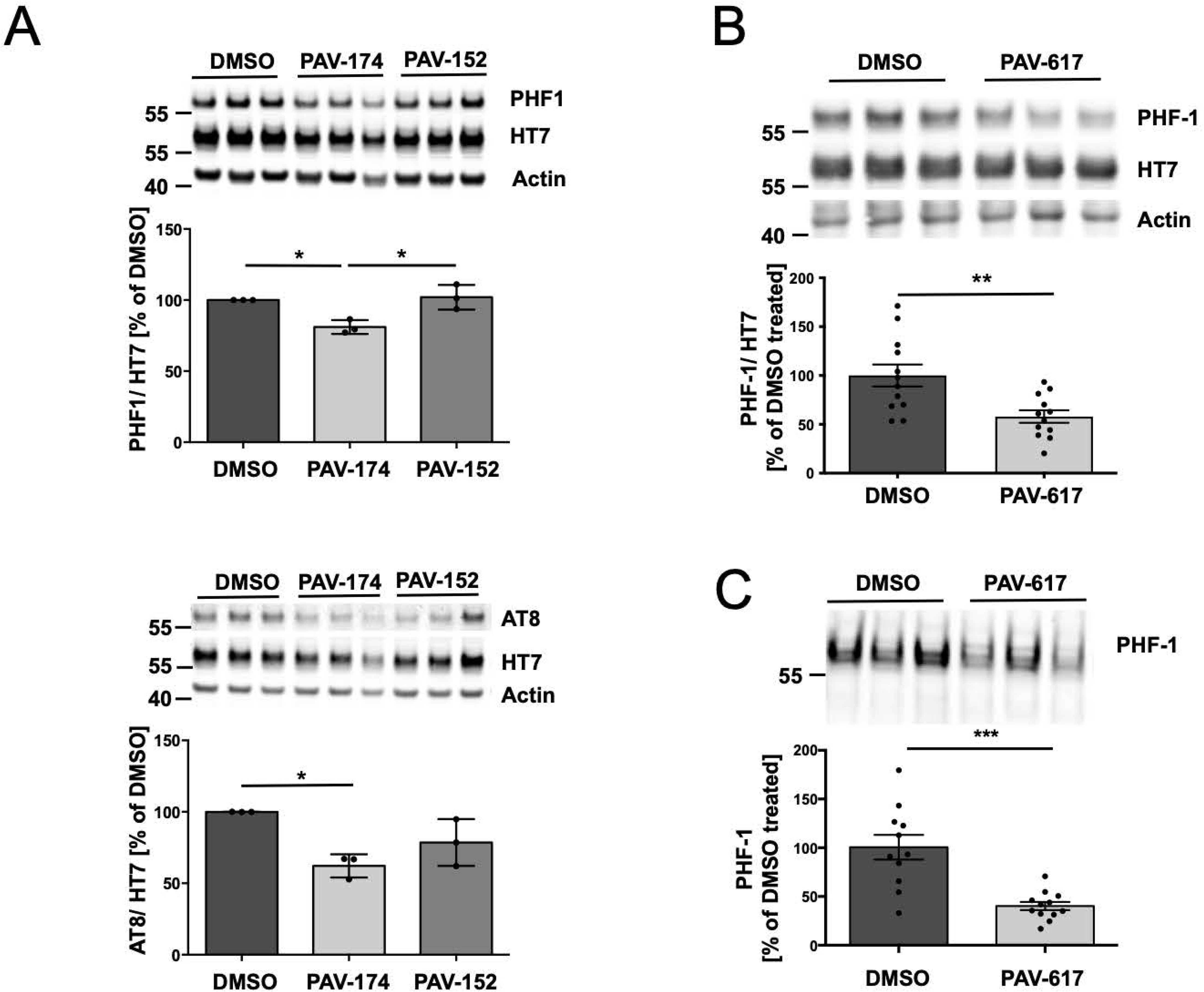
PAV-174 decreases tau phosphorylation and aggregation in vivo. (A) PAV-174 reduced tau phosphorylation (top: PHF-1; below AT8) in differentiated neurons derived from human iPSCs of a patient carrying the MAPT IVS10+16 mutation (V97). The neurons were incubated with PAV-174 (50 nM) or the non-functional analog PAV-152 for 48h. The diagram shows the average values of three independent experiments each with three technical replicates (n=3). Data were analyzed by one Way ANOVA (Tukey’s post-hoc). (B) Reduction of tau phosphorylation was observed *in vivo* after treating tau58/2 mice with 5 mg/kg of PAV-617. HT7 normalized phosphorylated tau (PHF-1) was reduced in the homogenates of the compound-treated mice. The diagram shows the average signals of 12 mice per treatment group derived from three independent Western Blots. Data were analyzed by unpaired two-tailed t-test. (C) Significant less phosphorylated tau (PHF-1) was also detected in the sarkosyl-insoluble fraction of PAV-617 treated mice. The diagram shows the average signals of 12 mice per treatment group derived from two independent Western Blots. Data were analyzed by unpaired two-tailed t-test. See also Figure S5.

### The oxMIF conformer is elevated in post mortem brains of patients with Alzheimer’s disease

To investigate a role of MIF in the brains of AD patients, we obtained brain tissue from the middle frontal gyrus derived from AD patients or healthy controls (**Supplementary Figure 6A**). Braak and Thal staging of *post-mortem* brains had been carefully conducted and was significantly higher in the AD group, whereas age, and post mortem delay (pmd) was equal (**Supplementary Figure 6B**). When we performed DRAC of original compound PAV-645 and eluted with more efficient compound PAV-174, we observed a higher concentration of MIF derived from *post-mortem* brain homogenates of patients with AD compared to the healthy controls (**Figure 5A; Supplementary Figure 6C**).

**Figure 5.**
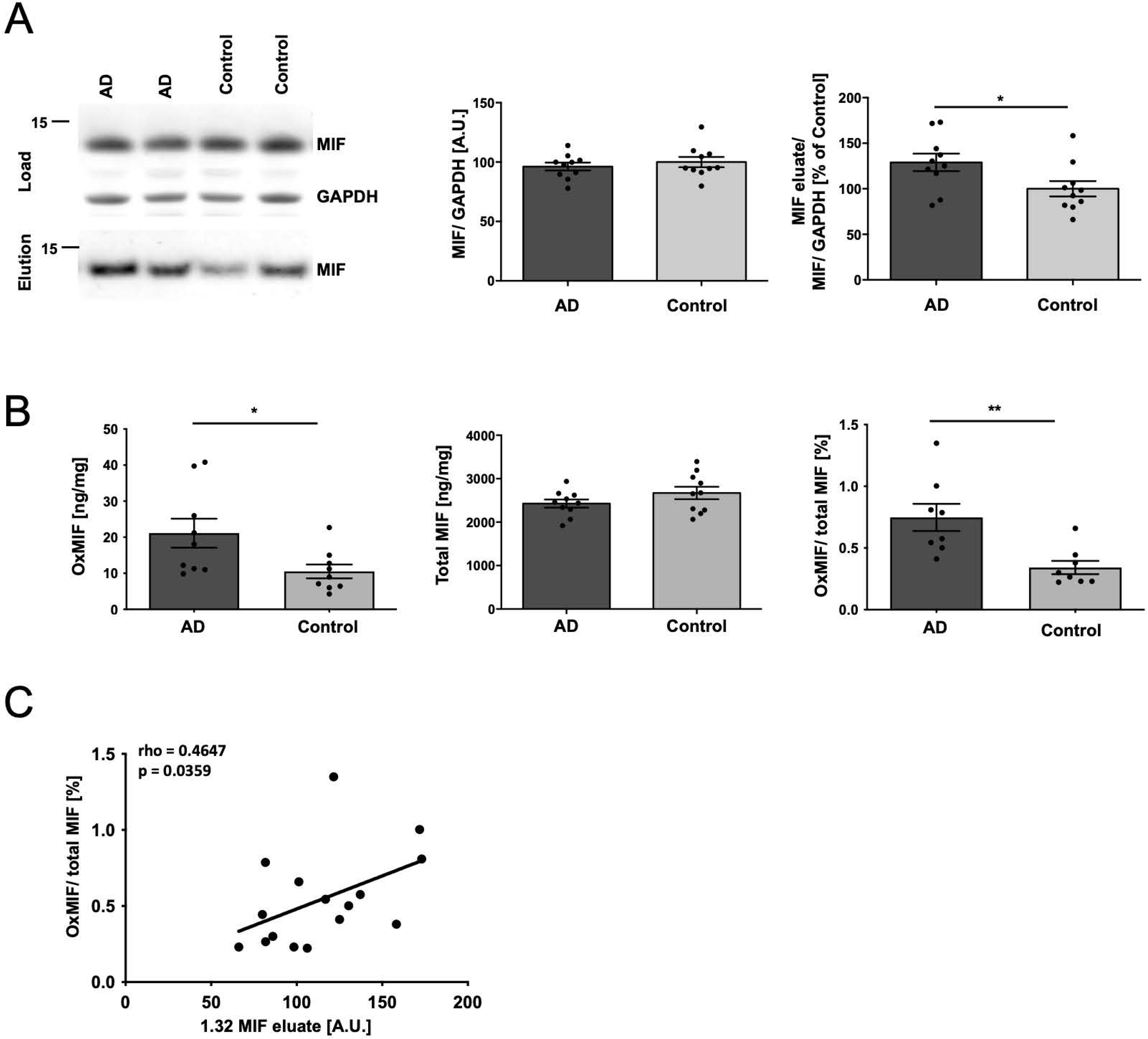
The oxMIF conformer is elevated in post mortem brains of patients with Alzheimer’s disease. (A) DRAC analysis of 20 brain samples from AD patients and age-matched healthy controls. Left: WB (representative section); upper panel shows the loading control (MIF and GAPDH) from brain homogenates applied to PAV-645 resin and the lower panel the pulled-down and PAV-174-eluted MIF. The GAPDH normalized MIF protein levels within the brain samples did not significantly differ between AD samples and healthy controls (left diagram). The eluted MIF was normalized to the input MIF/GAPDH signal (right diagram). Significantly more MIF was precipitated from AD brain samples by DRAC (displayed as % of the average in control samples). Each data point represents the average of two independent experiments. Data were analyzed by unpaired two tailed t-test. (B) Increased amounts of oxMIF in AD brain samples were detected by Sandwich ELISA. A significant higher oxMIF concentration (ng/mg protein) was detected in AD-brain tissue (left) whereas the total MIF concentration as measured by quantitative Western Blot did not differ (middle diagram; Supplementary Figure 6D). The ratio of oxMIF/ total MIF is shown in the right diagram. Each data point represents the average of two independent brain sample preparations. Data were analyzed by unpaired two tailed t-test. (C) Correlation analysis of the 20 brain samples between oxMIF (% of total MIF) levels detected by sandwich ELISA and the amount of MIF species eluted from PAV-645 resin. The oxMIF levels positively correlated with the amounts of MIF species eluted from PAV-645 resin. Spearmańs rho coefficient and p value (one-tailed) are indicated in the graph. See also Figure S6.

MIF has been reported in at least two conformers, an oxidized and a reduced form with the oxidized conformer being associated with disease (Kassaar et al., 2017; Thiele et al., 2015). A conformation-specific antibody for oxMIF, termed Imalumab, has been generated (Kerschbaumer et al., 2012) which we used to establish a sandwich ELISA assay allowing the detection of oxMIF in biological samples. As positive controls for establishing the ELISA, we used wild type MIF treated with Proclin 300 (Thiele et al., 2015), and MIF-C80W that mimics the oxMIF conformation (Schinagl et al., 2018). Total MIF levels were determined by Western blot since in ELISA, seemingly commercial all MIF antibodies had a bias towards oxMIF (data not shown). In these quantitations the oxMIF/total MIF ratio was higher in *post-mortem* brains from AD patients (**Figure 5B; Supplementary Figure 6D**) and the levels of oxMIF positively correlated with the MIF species eluted by DRAC (**Figure 5C**). Of note, we also detected an increase of oxMIF species in the supernatants of differentiated LUHMES cells upon infection with HSV-1 but found no changes in total MIF after infection with HSV-1 (**Supplementary Figure 6E**), corroborating the role of oxMIF in disease-associated states.

### OxMIF is a direct driver of tau phosphorylation modulated by PAV-174

To analyze the relevance of oxMIF on tau phosphorylation we exogenously applied recombinant wild-type MIF and the oxMIF surrogate MIF-C80W (Schinagl et al., 2018) to SH-SY5Y-tau-P301S cells. We observed a weak but significant induction of tau phosphorylation by MIF-C80W, but not with wild-type MIF after 6h (**Figure 6A; Supplementary Figure 7A**). This effect was not observed in MIF-ko cells (**Supplementary Figure 7B**). The induction of tau phosphorylation by MIF-C80W was inhibited when the oxMIF conformer was incubated with equimolar concentrations of PAV-174 1h before application on the cells (**Figure 6B; Supplementary Figure 7C**). By using DRAC assay we observed stronger binding of wild-type MIF to PAV-645 after MIF was oxidized with H_2_O_2_ following a protocol of Skeens et al. (Skeens et al., 2022) This prompted us to analyze the binding of PAV-174 to oxidized MIF by NMR. The ^15^N-HSQC spectra of wildtype MIF (redMIF) and H_2_O_2_ treated wild type-MIF (oxMIF), in the presence or absence of PAV-174, revealed chemical shift perturbations (CSP) upon PAV-174 addition, confirming that PAV-174 bound to both MIF species. However, the low alignment between the chemical shifts of oxMIF and redMIF did not allow to transfer the assignments of the wild-type redMIF ^15^N-HSQC spectrum onto the oxMIF ^15^N-HSQC spectrum. Therefore, the binding site of PAV-174 in oxMIF could not be definitely determined to be precisely the same as for wild-type redMIF. It is worth noticing though that the resonance and shifts corresponding to the Trp108 side chain are easily aligned between MIF conformers suggesting that the PAV-174 binds at the same site or in close proximity in both conformers.

**Figure 6.**
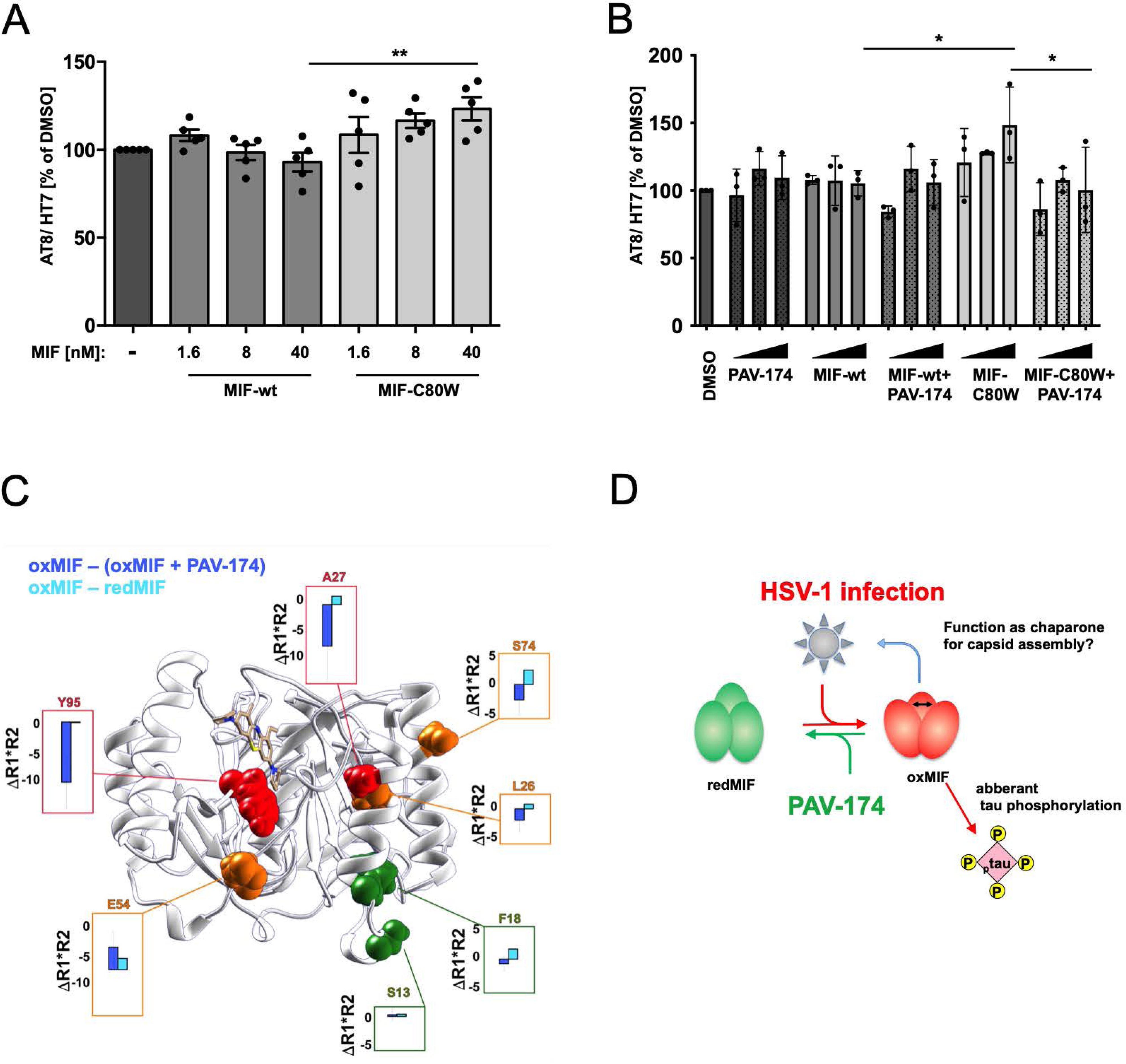
OxMIF is a direct driver of tau phosphorylation modulated by PAV-174. (A) Exogenous applied oxMIF induce tau phosphorylation. SH-SY5Y-tau-P301S-control and–MIF-ko cells were treated with recombinant wt-MIF or MIF-C80W for 6h. MIF-C80W dose dependently induced tau phosphorylation in control cells but not in MIF-ko cells (see Supplementary Fig. 7A/B). The diagram shows the average values of five independent experiments (n=5). Data were analyzed by One-way ANOVA (Sidak’s post-hoc). (B) Preincubation of recombinant MIF-C80W with equimolar concentrations of PAV-174 prevented phosphorylation of tau. 1.6 nM, 8 nM or 40 nM of MIF-C80W were incubated with equimolar concentration of PAV-174 or the equal volume of DMSO for 1h before the mixture was added to SH-SY5Y-tau-P301S cells. As a control PAV-174 was also provided without MIF. The significant increase of tau phosphorylation by MIF-C80W after 6h incubation was abolished by PAV-174. The diagram shows the average values of three independent experiments (n=3). Data were analyzed by One-way ANOVA (Sidak’s post-hoc). (C) Structural dynamic cancelation of compound PAV-174 on oxMIF visualized on the Structure of MIF bound to PAV-174. For the residues highlighted as spheres, the ^15^N NMR relaxation product difference (⊗R1*R2) between oxMIF and oxMIF + PAV-174 (deep blue), and between oxMIF and redMIF (Cyan) is displayed in the boxes. The residues colored in red show a strong cancellation of the motion on the microsecond timescale upon addition of PAV-174, moderate in orange, and little or no effect in green (D) Hypothetic model of MIF isoforms implicated in HSV-1 life cycle and tau phosphorylation. See also Figure S7.

To investigate a potential effect of PAV-174 on differential multimerization of wildtype MIF and MIF-C80W we performed high-pressure liquid chromatography - multi-angle laser light scattering (HPLC-MALS), using a size exclusion column. The HPLC runs were performed for wildtype and the C80W mutant in the presence or absence of PAV-174 (**Supplementary Figure 7E**). The addition of PAV-174 did not lead to different retention times in both MIF species. Furthermore, the molecular weight of the observed species was calculated for each sample: wt-MIF: 34,140 kDa +/− 2%, wt-MIF + PAV-174: 35,270 kDa +/− 2%, MIF-C80W: 37,890 kDa +/− 7%, and MIF-C80W + PAV-174: 37,180 kDa +/− 5%, using the Astra software, correcting the Rayleigh ratio by the UV absorbance at 280 nm. These results indicated that MIF is present as a trimer regardless of the interaction with PAV-174 and that PAV-174 does not perturb the monomer-trimer equilibrium or the oligomerization of both wt-MIF and MIF-C80W (**Supplementary Figure 7E**). The different retention times of wildtype MIF and MIF-C80W are likely due to different conformations of both MIF species that have been described (Schinagl et al., 2018). The resolution of the HPLC-MALS is not in the atomic range and therefore only reports conformation changes at the quaternary structure scale (i.e., the formation of multimers) and to some extent tertiary structure scale (i.e., unfolding). For this reason, it cannot be excluded that PAV-174 affects the secondary/tertiary structure or dynamics for the two MIF conformers.

## Discussion

The molecular interfaces linking external risk factors such as subchronic inflammation to the development of neurodegeneration are poorly understood. Here we demonstrate that oxidized MIF is a cellular response to an external stressor, HSV-1, that mediates tau phosphorylation, a critical event in the genesis of neurodegenerative diseases such as Alzheimer’s disease (AD) or subtypes of frontotemporal dementias (FTD). We also identified a novel compound that, upon binding MIF, stabilizes a non-oxMIF conformation and thereby reverses tau phosphorylation and aggregation *in vitro* and *in vivo*, respectively, demonstrating that this class of pharmaceuticals could become relevant for treating sporadic forms of neurodegenerative diseases such as sporadic AD.

Our findings are remarkable in several ways:

1. Independent of the link to tau phosphorylation, blocking oxMIF obviously is an excellent strategy to prevent HSV-1 infection. In all three cellular assay systems used (see **Figure 1**), Vero cells, primary-like LUHMES cells and human brain organoids, lead compound PAV-174 efficiently interfered with HSV-1 replication at an IC_50_ in a low nanomolar range - acyclovir, the standard competitive inhibitor of herpes virus DNA polymerase, is efficient at only 8.5 µM IC_50_ in Vero cells (Brand et al.). Remarkably, PAV-174 showed no induction of apoptosis in human brain organoids indicating that it prevents HSV-1 infection without being neurotoxic under the conditions used. Since generalized herpes virus encephalitis remains a serious neurological diagnosis with a high mortality (20% even under acyclovir therapy, and severe lasting post-infection symptoms (Gurgel Assis et al., 2021), the pharmacological target oxMIF and the class of compounds presented here may prove to significantly increase antiherpetic pharmacotherapy for the benefit of subacutely HSV-1-infected patients. Antiviral therapy targeting a host-viral interface is thus highly efficient, and in continuity with previous research on antiviral compounds against rabies virus (Lingappa et al 2013), or HIV (Reed et al., 2021).

Intriguingly, despite the fact that inhibition of MIF by PAV-174 is highly effective, the presence of MIF appears not to be essential for HSV-1 life cycle (**Supplementary Figure 4B**). The significant increase in level of capsid protein without enhancement of viral titer argues that, as predicted, misassembly of HSV-1 capsid was achieved by either treatment with PAV-174 or knockout of MIF. That is, the capsid signal/ virus titer ratio increased by 58% or 43% in MIF-ko cells that were infected with HSV-1 at an MOI of 0.2 or 0.4, respectively (**Supplementary Figure 4B**). The lack of a higher effect on reducing the viral titer could be explained by the presence of *D*-Dopachrome Tautomerase (D-DT) a structural cellular homolog of MIF (Sugimoto et al., 1999) that has been reported to share essential functions with MIF (summarized in (Merk et al., 2012)). We did not consider to engineer an additional knock out of D-DT since the combined loss of MIF and D-DT has been shown to increase programmed cell death by inhibition of p53 (Brock et al., 2014).

2. Protein misfolding is ultimately the result of disturbed proteostasis and, accordingly, protein misfolding diseases (PMD) characterized by the deposition of misfolded, aggregated proteins are fundamentally caused by disturbed proteostasis. For the genetic cases of PMD, it is easy to understand that a mutant protein is eventually more prone to misfolding or aggregation. It is less obvious how a similar misfolding is triggered by external factors in the majority of sporadic cases of PMD, i.e., in the absence of mutations in the aggregating protein. We have previously suggested that if a specific viral infection results in a cellular pathology similar to that characteristic for a given PMD, the dissection of host proteins that this virus recruits, e.g. for its (capsid) assembly and that hence are causing the disruption in cellular proteostasis, may reveal important clues about molecular interfaces involved in triggering the given PMD even in the *absence* of virus (Müller-Schiffmann et al., 2021).

We here demonstrate the usefulness of this concept: HSV-1 infection increases tau phosphorylation, a hallmark of aggregated tau, at PMD-relevant residues (Augustinack et al., 2002) *in vitro*, and this can be prevented by lead compound PAV-174 in a MIF-dependent manner since a complete cellular knockdown of MIF cancels PAV-174’s effects (**Figure 4E**). Our data are in accordance with previous findings that demonstrated that the absence of MIF decreased tau phosphorylation in a mouse model of AD (Li et al., 2015). By binding to its canonical receptors CD74 (Jankauskas et al., 2019), and CXCR4 (Bonham et al., 2018; Martinez-Martin et al., 2015; Rajasekaran et al., 2016), MIF interferes with Akt/ GSK3β signaling that are important signaling cascades involved in tau phosphorylation(Schwartz et al., 2009) (**Supplementary Figure 4A/B**) consistent with our observations that PAV-174 specifically reduces phosphorylation at sites (AT8, AT180, PHF-1) modulated by GSK3β. Accordingly, we did not observe a comparable effect of PAV-174 on preventing phosphorylation of residues with not affected by GSK3β (**Supplementary Figure 3B**).

Closely related compound PAV-617 which had a similar effect *in vitro* on tau phosphorylation but a more favorable pharmacokinetic profile, not only decreased tau phosphorylation after four weeks of treatment in mice, but also tau aggregation (**Figure 5**) indicating that this class of compounds targeting oxMIF is highly efficient in blocking tau phosphorylation and aggregation *in vivo*.

MIF is a key factor in mediating inflammatory responses and in addition to its role in the HSV-1 life cycle described here, MIF is also involved in relaying other external triggers of sAD. For example, serum levels of MIF are markedly increased in patients suffering from insulin resistance and type 2 diabetes (summarized in (Morrison and Kleemann, 2015)). The role of different MIF conformers – existing in at least in one oxidized and one reduced conformation - is not yet widely discussed for a role in regulating physiological circuitry. Studies have reported increased oxMIF levels in chronic inflammatory diseases (Thiele et al., 2015) of which some have been reported to present a higher risk to develop dementia, such as asthma, sepsis, psoriasis, ulcerative colitis, Crohn’s disease and systemic lupus erythematosus (Gendelman et al., 2018; Kim et al., 2020; Liu et al., 2022; Nair et al., 2022; Thiele et al., 2015; Zhao et al., 2018). Therefore, we speculate that oxMIF has the potential to be a molecular interface in relaying external events at a broader scale to tau phosphorylation and aggregation, thereby accelerating the risk to develop dementia. Along these lines, the increased oxMIF levels in our AD brain samples (**Figure 5B**) were likely induced by various external stressors, and not restricted to HSV-1.

3. Since antibodies against β-sheeted components of MIF exhibited the largest anti-inflammatory potential (Kerschbaumer et al., 2012) selective antibodies against oxidized MIF were generated and characterized (Schinagl et al., 2018; Thiele et al., 2015). A phase 1 study of one of these antibodies, imalumab, in patients with solid tumors have been published (Mahalingam et al., 2020). However, in the absence of follow up publications, earlier clinical studies on imalumab in antitumor therapy seem to have been abandoned (Douillard et al., 2015) and thus the clinical usefulness of oxMIF-specific antibody therapy remains, as opposed to, for example the class of compounds presented here, to be demonstrated.

Recently, a structure of oxidized MIF was published (Skeens et al., 2022). Remarkably, our structural investigations reveal that binding of PAV-174 to oxMIF restores, to some degree, the redMIF conformation and thus may revert its proinflammatory effects, explaining its modulation on Akt signaling and tau cellular pathology. The effect of PAV-174 seems to be allosteric since the ^15^N-HSQC reveals remote CSPs from the binding site of PAV-174. Among the strongest CSPs, we observe Lys66, Ser63, Gln102, Ala38, Ile96, which are all distant from the binding site of PAV-174 (**Figure 2A**). The differences in the spin relaxation rates ⊗R_1_R_2_ were obtained by NMR (Lakomek et al., 2012) reveal an important reversal of the motion in the microsecond scale for the residues Ala27, Leu26, and Ser74, forming a network connecting the top-end of the two alpha helixes. Furthermore, we can observe a regiospecificity of the effect of PAV-174 binding, as the residues Ser13 and Phe18, located at the bottom-end of the first alpha helix and do not see their dynamics affected upon PAV-174 binding as observed in the spin relaxation rate differences ΔR_1_R_2_ (**Figure 6C**). Notably, the binding site is distinct from the one of ISO-1 and therefore does not seem to involve the tautomerase activity ((Al-Abed et al., 2005; Bloom et al., 2016)) which, accordingly, also did not display any tau phosphorylation-decreasing effects (Figure 3F).

Methylene blue (MB; in this study termed 1.16) is a phenothiazine core derivative, as are both PAV-174 and PAV-617. MB was previously reported to be an inhibitor of tau aggregation, including *in vivo* (Hochgräfe et al., 2015) and is currently studied in a clinical trial for the treatment of frontotemporal dementia and AD (https://www.clinicaltrials.gov/ct2/show/NCT03446001). For two independent reasons we believe the activity observed in our assays is different from those observed for MB by others. First, in our *in vitro* assay PAV-174 was around 2-logs more potent than MB in inhibiting HSV-1 replication or tau phosphorylation. PAV-617 retained that enhanced activity but with superior PK enabled demonstration of *in vivo* effects on tau phosphorylation and aggregation. Second, our NMR data suggest that the mode of action is likely different from that of MB since one of the unique pyrrolidine rings of PAV-174 (not present in MB) is deeply embedded in the hydrophobic pocket formed by Tyr36, Ile64, Val106, Trp108, and Phe113. We hypothesize that the strong hydrophobic interaction with residues located close to the core of the protein engages the allosteric network. The second pyrrolidine ring is pointing outwards from the binding site of MIF. Despite, its solvent exposure, its proximity with the indole of Trp108 suggests potential π-interaction with this residue, which is facing on the other side Cys80, again suggesting its key importance in allosteric effects of PAV-174. Although from our data we cannot exclude that part of PAV-174 effects may overlap with those of MB, our functional data in MIF-ko cells suggest that the main mechanism of PAV-174 occurs via allosteric targeting of MIF. PAV-174 and PAV-617 have previously been observed to have antiviral effects also on pox viruses (Priyamvada et al., 2021). The relationship between these effects and the mechanisms discussed here are currently unknown but are worth to be investigated.

4. The long-known epidemiological association between latent herpes virus reactivation and AD (Allnutt et al., 2020; Itzhaki et al., 2020; Lovheim et al., 2015; Readhead et al., 2018; Tzeng et al., 2018) (Allnutt et al., 2020; Itzhaki et al., 2020; Levine et al., 2023; Lovheim et al., 2015; Readhead et al., 2018; Tzeng et al., 2018) are opposed by much fewer studies unable to find this association in different cohorts (Murphy et al., 2021; Warren-Gash et al., 2022). The molecular mechanism of potentially triggering effects of herpes virus activation have remained unexplained, even though several molecular scenarios have been proposed such as a role for Aβ as a cellular defense (Eimer et al., 2018) or an interference of herpes virus with autophagy (Cirone, 2018; Orvedahl et al., 2007). Our results suggest that a small subfraction of MIF, present in the oxidized MIF conformation, presents the missing molecular link between HSV-1 infection and AD-like cellular pathology, since it has a role both in HSV-1 life cycle and tau phosphorylation. Whether other proteins or even a multiprotein complex that includes oxMIF are involved remains currently open.

We put forward a model (**Figure 7D**) where we speculate that upon HSV-1 infection the induced oxMIF could function as a chaperone, or in some other role, in the process of HSV-1 capsid assembly. This general function of MIF has been described for SOD1, although the specific participation of the oxMIF conformer in the MIF-SOD1 interaction was not studied (Israelson et al., 2015). In addition to the likely many specific functions of oxMIF, the presence of oxMIF increases tau phosphorylation. Both effects can be reverted by PAV-174 by re-establishing the redMIF conformation thereby reducing both efficient capsid assembly as well as aberrant tau phosphorylation. Our discovery of one particular conformer of MIF, oxidized MIF, that has been described as proinflammatory and disease-associated in various contexts (Thiele et al., 2015) is upregulated in *post-mortem* brains from patients with AD (**Figure 4B**) now defines the known role of MIF in AD (Bacher et al., 2010; Petralia et al., 2020) more precisely and corroborates a proinflammatory state in AD.

In conclusion, we have presented a novel, oxMIF-targeted anti-HSV-1 compound with several notable features related to neurodegenerative disease therapeutics. This compound prevents aberrant tau phosphorylation and aggregation and hence is likely relevant to potential drugs against tau-dependent neurodegenerative disorders like AD, independent of HSV-1 infection. MIF, and possibly other proteins in conjunction with MIF in a multiprotein complex, are at the intersection between HSV-1 replication and AD-like cellular pathology providing a molecular basis for further analyzing the long-known epidemiological connection between these disorders. The reported functional pleiotropy of MIF suggests that also other AD risk factors may relay via the MIF molecular interface.

## Acknowledgements

We thank Jesus Requena for critical discussion. This research was funded by BMBF REMOVAGE (#01GQ1422A), the Research Commission of the Medical Faculty of the Heinrich Heine University Düsseldorf (#9772726), the DFG (KO1679/10-1, 15-1), the Deutsche Alzheimer Gesellschaft, and a grant from Prosetta Biosciences.

## Author contributions

AMS, CK and VRL conceived the broad hypotheses and concepts. AMS, SKT, IP and VB performed experiments. AK synthesized PAV-174 and key analogs described. AR and JG prepared the human brain organoids. AA and SW conducted the iPSC experiments. SY and DD performed early DRAC studies and sample preparation for MS-MS. SM and DP designed the mouse PK and toxicity studies. VL established the CFPSA system, contributed to DRAC design and writing the manuscript. FT performed and analyzed the NMR experiments and the resulting 3D structures. RR designed, analyzed, and supervised the NMR experiments and the resulting 3D structures. AR provided and staged the post mortem brains. AMS, VRL, FT and RR and CK wrote the manuscript and all authors provided feedback and comments on the initial version.

## Inclusion and diversity statement

Senior investigators in this study (CK, VL, RR) in this study affirm their support for diversity and inclusion as demonstrated but not limited to the diversity of coauthors in regards to gender, age, and creed.

## METHOD DETAILS

## KEY RESOURCE TABLE

**See separate file**

## RESOURCE AVAILABILITY

### Lead Contact

Further information and requests for resources and reagents should be directed to and will be fulfilled by the lead contact, Carsten Korth.

## EXPERIMENTAL MODEL AND SUBJECT DETAILS

### Animals and human samples and ethics

TgTau58/2 mice (van Eersel et al., 2015) were a kind gift from Novartis (Boston, USA). They were maintained in the Animal Facility at the University of Düsseldorf, Germany, fed ad libitum with standard laboratory chow and water in ventilated cages under a 12h light/dark cycle. All experiments were conducted in conformity with the Animal Protection Law approved by local authorities (LANUV NRW, Recklinghausen, Germany).

Human brain samples were obtained from the The Netherlands Brain Bank, Netherlands Institute for Neuroscience, Amsterdam (www.brainbank.nl). Use of these samples for this study was approved by the Ethics Commission of the Medical Faculty of the Heinrich Heine University, Düsseldorf.

### Cell Culture

Vero (ATCC), GP2-293 (Clontech) and HEK-293FT cells (Clontech) were cultured in DMEM (Gibco) supplemented with 10% FBS xtra (Capricorn Scientific), 1 mM sodium pyruvate (Gibco), and 1% (v/v) penicillin/streptomycin (Sigma). Human neuroblastoma SH-SY5Y cells were obtained from the DSMZ (Leibniz Institute DSMZ-German Collection of Microorganisms and Cell Cultures, Braunschweig, Germany) and cultured in DMEM/F12 Medium (Gibco) supplemented with 10% FBS xtra (Capricorn Scientific), 1x non-essential amino acids (MEM NEAA, Gibco), 1% (v/v) penicillin/streptomycin (Sigma). LUHMES cells (ATCC) were maintained in a proliferative state in advanced DMEM/F12 (Gibco) supplemented with 1% N2 (Thermo Fisher Scientific), 1% (v/v) penicillin/streptomycin (Sigma), 2 mM L-glutamine (Gibco) and 40 ng/mL bFGF (basic recombinant human fibroblast growth factor; Sigma) in flasks coated first with 50 µg/mL poly-L-ornithine (Sigma) and then with 1 µg/mL fibronectin (Sigma). For differentiation into post-mitotic neurons the cells were cultured in advanced DMEM/F12 supplemented with 1 µg/mL doxycyclin (Sigma), 2 ng/mL GDNF (glial cell line-derived neurotropic factor; Sigma), and 1 mM dibutyryl-cAMP (Santa Cruz Biotechnology) in coated flasks for 2 days according to Scholz et al. (Scholz et al., 2011). The cells were then washed with PBS and trypsinized with Trypsin-EDTA solution (Gibco), counted and seeded on target plates in differentiation medium.

Culture and differentiation of TSM(exon10+16)V97 iPSC were performed as previously described, and all reagents were purchased from Thermo Fisher Scientific unless otherwise stated (Arber et al., 2020). iPSC were cultured on geltrex (Thermo Fisher Scientific) coated plates in Essential-8 media. iPSC were grown to 100% confluency prior to neuronal induction using dual SMAD inhibition as described previously (Shi et al., 2012). Briefly, cells were cultured for 10 days in neural induction media (N2B27 containing 10 µM SB431542 (Tocris) and 1 µM dorsomorphin (Tocris)). N2B27 media consists of a 1:1 mixture of Dulbecco’s modified eagle medium F12 (DMEM-F12) and Neurobasal, supplemented with 1 × N-2, 1 × B-27, 5 µg/mL insulin, 1 mM l-glutamine, 100 µM nonessential amino acids, 100 µM 2-mercaptoethanol, 50 U ml−1 penicillin and 50 mg/mL streptomycin. At days 10 and 18, neuronal rosettes were passaged using dispase and plated in laminin-coated wells in N2B27 media. The final passage was performed at day 35 using accutase, and cells were plated at a final density of 50,000 cells per cm^2^ and maintained in N2B27 media until treatment. Mature neurons were treated at 70-90 DIV and harvested 24-48h post-treatment.

Organoids were generated from a commercially available hiPSC line (IMR90, Wi Cell). The hiPSCs were maintained on mTeSR medium, under non differentiating conditions for initial expansion. To differentiate into the neural lineage for organoid generation, the hiPSCs were allowed to self-aggregate in 96-well plates for five days using Neural induction medium (Stem cell technologies) as described earlier (Gabriel et al., 2017). The aggregated neurospheres were further matured to form brain organoids in spinner flaks, in a medium composed of DMEM-F12: Neural Basal medium (1:1), 1:100 B27 v/o Vitamin A (Thermo Scientific), 1:200 N2 (Thermo Scientific), 1:100 L-Glutamine (Gibco), 100 µg/mL Primocin (Thermo Fisher Scientific), 0.05% Insulin (Sigma Aldrich) and 0.1% Matrigel (Corning). All cells were tested at regular time intervals for mycoplasma contamination and cultured at 37°C with 5% CO_2_.

#### Chemical synthesis

##### 1-Ethyl-9-methylphenothiazin-5-ium tetraiodide hydrate

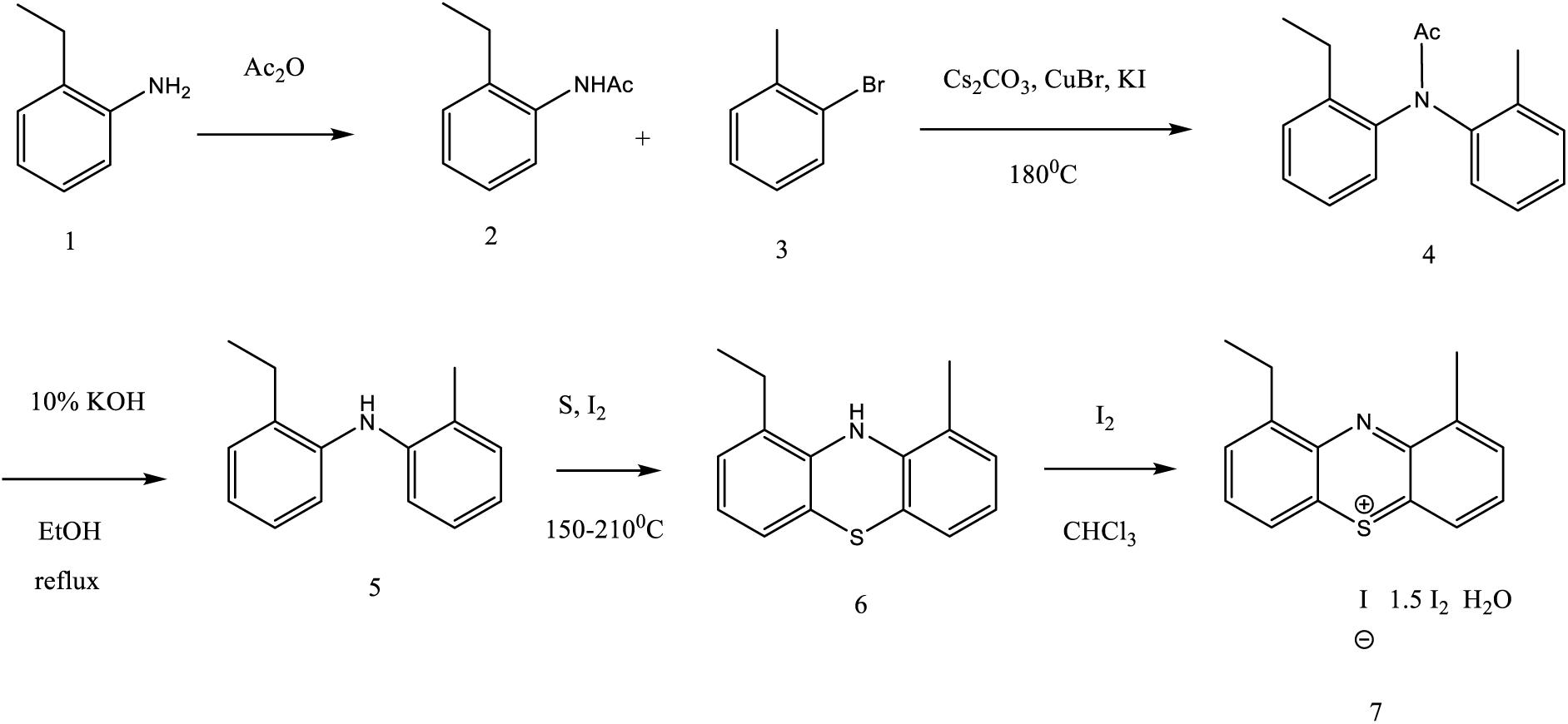

a. N-Acetyl-2-ethylaniline (2): commercial 2-ethylaniline (1) (50mL, 0.40 mol) was dissolved in acetic anhydride (160mL, 1.70 mol) and stirred at room temperature for 2h. Then the reaction mixture was poured into H_2_O, the whole was extracted with ethyl acetate (2×200mL). The combined organic extracts were washed with 5% aqueous NaHCO_3_, brine, dried (K_2_CO_3_), filtered and concentrated to provide the title compound as a white solid (60.0 g, 92%).
b. N-Acetyl-2-ethyl-2’-methyldiphenylamine (4): a mixture of the N-acetyl-2-ethylaniline (35.0 g, 215 mmol), anhydrous Cs_2_CO_3_ (70.0 g, 215 mmol), CuBr (2.86 g, 20 mmol), KI (3.33 g, 20 mmol) and 2-bromotoluene (3) (78mL, 640 mmol) was stirred and heated at 175-180^0^C under an argon atmosphere for 48h. After cooling the reaction mixture was poured into ice-H_2_O and extracted with ethyl acetate (2×200mL), the combined organic extracts were washed with brine, dried over anhydrous K_2_CO_3_, filtered and concentrated to dryness. The obtained crude material was purified by flash chromatography (using ethyl acetate - hexane as an eluent) to afford the N-acetyl-2-ethyl-2’-methyldiphenylamine (35.4 g, 65%).
c. 2-Ethyl-2’-methyldiphenylamine (5): a solution of the N-acetyl-2-ethyl-2’-methyldiphenylamine (32.5 g, 128 mmol) in 10% KOH (72 g, 1.28 mol)/EtOH (120mL) was stirred and refluxed for 6 h, then poured into H_2_O. The mixture was extracted with ethyl acetate (2×100mL). The combined organic layers were washed with brine, dried (Na_2_SO_4_), filtered and concentrated to dryness, gave dark red oil (21.1g, 78%).
d. 1-Ethyl-9-methyl-10H-phenothiazine (6): to a 2-ethyl-2’-methyldiphenylamine (3.0 g, 14.2 mmol), sulfur (909 mg, 28.4 mmol) and iodine (601 mg, 4.7 mmol) were added. Vial was charged with balloon for discharge. The heating block was preheated (150^0^C). The vial was heated on the heating block and after 15 min. temperature was increased to 210^0^C, reaction mixture was stirred and heated for an additional 1 h. The mixture was allowed to cool to 90^0^C. The dark solid material was dissolved in mixture methanol/chloroform and purified by flash chromatography (ethyl acetate - hexane as an eluent) to afford the desired product (790 mg, 23%).
e. 1-Ethyl-9-methylphenothiazin-5-ium tetraiodide hydrate (7) : a solution of 1-ethyl-9-methyl-10H-phenothiazine (4.83g, 20 mmol) in anhydrous chloroform (50mL) was stirred at 5^0^C and the solution of iodine (15.25 g, 60 mmol) in CHCl_3_ (300mL) was added drop wise over 3h. The resulting dark solution was stirred for an additional 3h at 5^0^C, monitored by TLC. After the disappearance of the starting material, the resulting precipitate was filtered, washed with a copious amount of chloroform, dried overnight in vacuum to afford a dark solid (9.18 g, 60%).

##### 3,7-Di-(4-methylpiperazin-1-yl)-1-ethyl-9-methylphenothiazin-5-ium iodide (PAV-152)

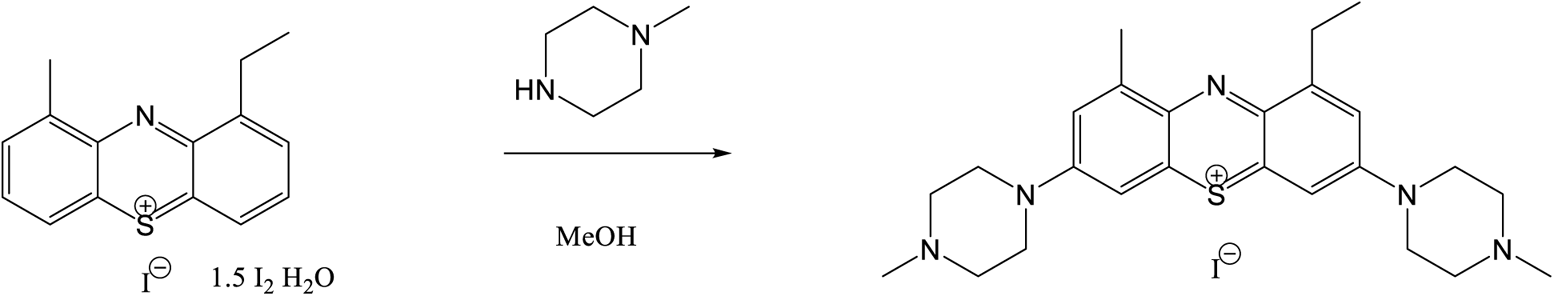

A solution of 1-ethyl-9-methylphenothiazin-5-ium tetraiodide hydrate (50 mg, 0.07 mmol) in methanol (10mL) and 1-methylpiperazine (30 mg, 0.3 mmol) was stirred for 2 h at room temperature. The resulting mixture was concentrated to dryness and purified by flash chromatography using the methanol-chloroform gradient to provide the title compound.

##### 3,7-Di-(morpholin-1-yl)-1-ethyl-9-methylphenothiazin-5-ium iodide (PAV-215)

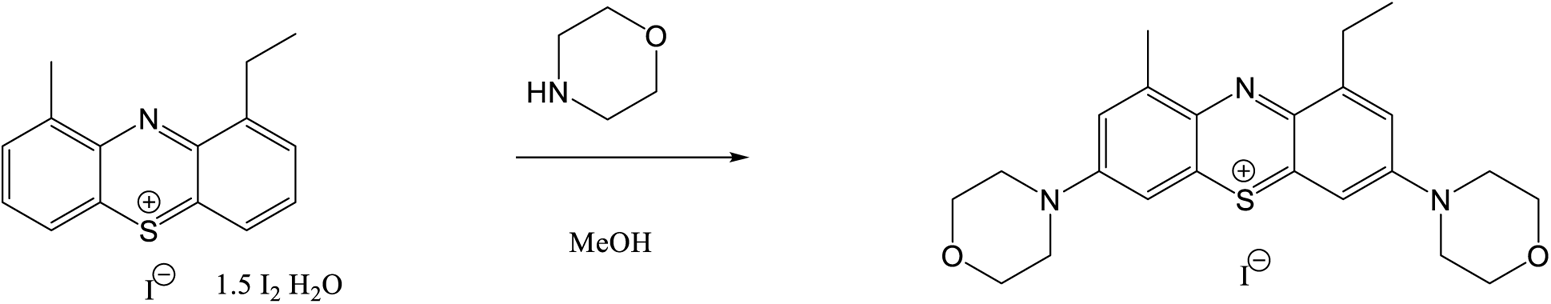

A solution of 1-ethyl-9-methylphenothiazin-5-ium tetraiodide hydrate (50 mg, 0.07 mmol) in methanol (10mL) and morpholine (0.05 mL, 0.5 mmol) was stirred for 1 h at room temperature. The resulting mixture was concentrated to dryness and purified by flash chromatography using the methanol-chloroform gradient to provide the title compound.

##### 3,7-Di-(4-(dimethylamino)piperidin-1-yl)-1-ethyl-9-methylphenothiazin-5-ium iodide (PAV-251)

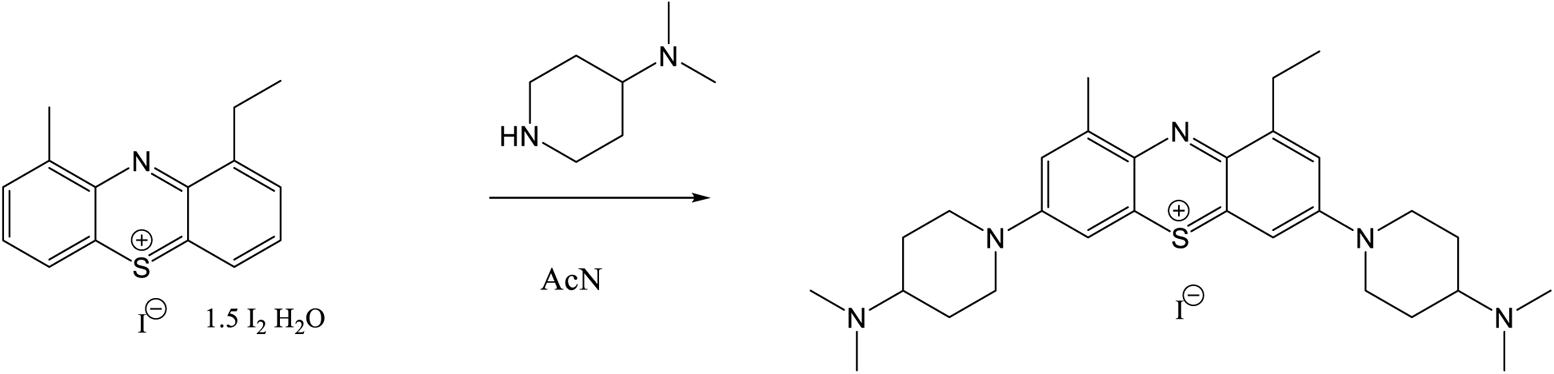

A solution of 1-ethyl-9-methylphenothiazin-5-ium tetraiodide hydrate (382 mg, 0.5 mmol) in acetonitrile (10mL) and 4-(dimethylamino)piperidine (192 mg, 1.5 mmol) was stirred for 1 h at room temperature. The resulting mixture was concentrated to dryness and purified by flash chromatography using the methanol-chloroform gradient to provide the title compound.

##### 3,7-Di-(tiomorpholin-1-yl)-1-ethyl-9-methylphenothiazin-5-ium iodide (PAV-269)

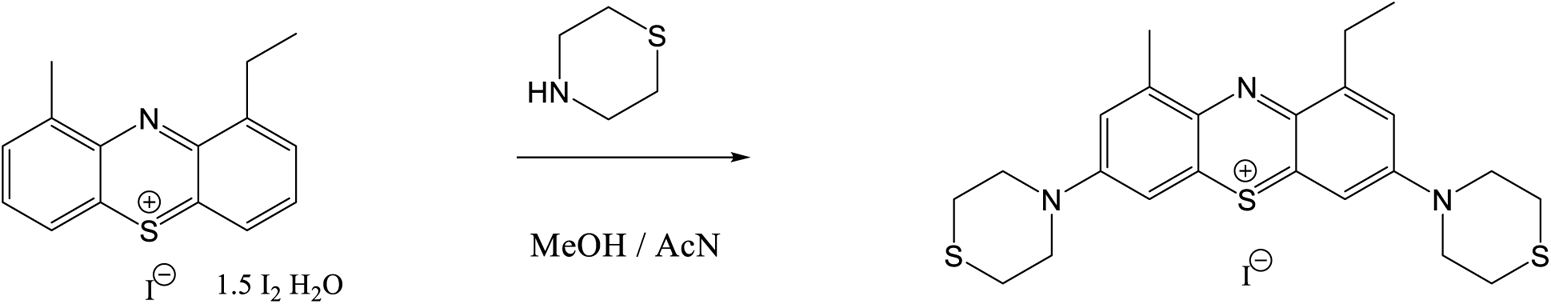

A solution of 1-ethyl-9-methylphenothiazin-5-ium tetraiodide hydrate (191 mg, 0.25 mmol) in mixture methanol and acetonitrile (1:1) (10mL) and tiomorpholine (0.1 mL, 1.0 mmol) was stirred for 1 h at room temperature. The resulting mixture was concentrated to dryness and purified by flash chromatography using the methanol-chloroform gradient to provide the title compound.

##### 3,7-Di(piperazin-1-yl)-1-ethyl-9-methylphenothiazin-5-ium trifluoroacetate (PAV-382)

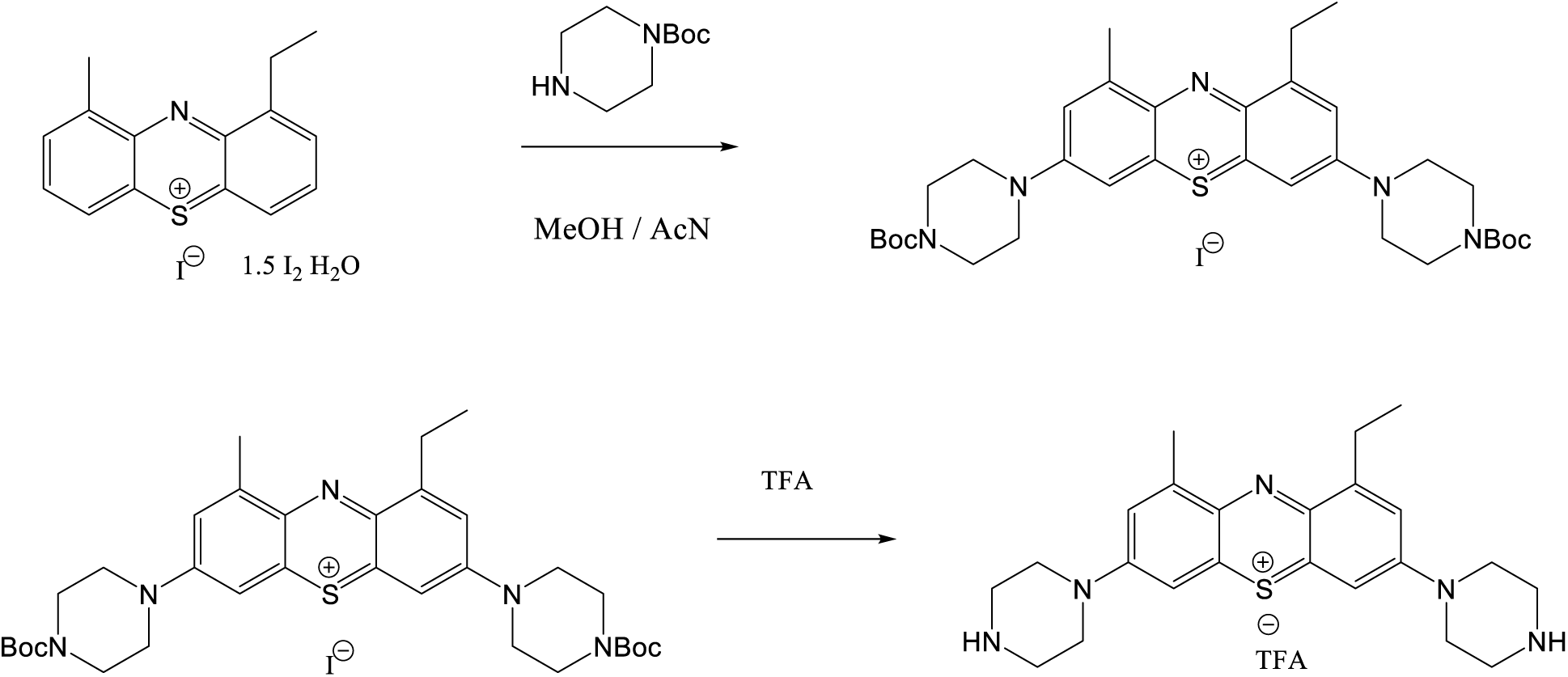

A solution of 1-ethyl-9-methylphenothiazin-5-ium tetraiodide hydrate (153 mg, 0.2 mmol) in mixture methanol and acetonitrile (1:1) (10mL) and N-Bocpiperazine (186 mg, 1.0 mmol) was stirred for 1 h at room temperature. The resulting mixture was concentrated to dryness and dissolved in DCM. TFA (1 mL) was added with stirring. After 30 min. mixture was concentrated and purified by prep-HPLC to provide the title compound.

##### 3,7-Di(homopiperazin-1-yl)-1-ethyl-9-methylphenothiazin-5-ium trifluoroacetate (PAV-352)

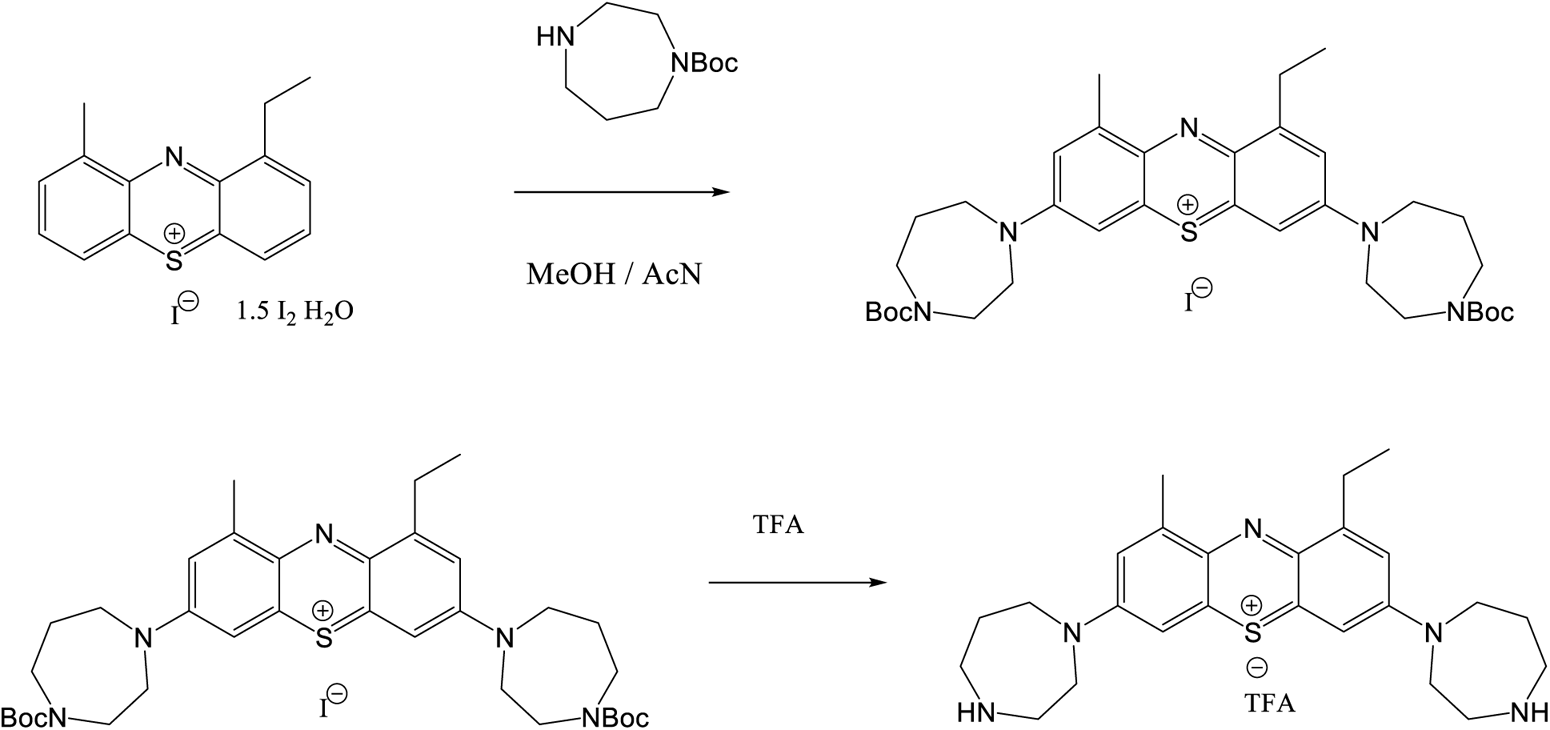

A solution of 1-ethyl-9-methylphenothiazin-5-ium tetraiodide hydrate (153 mg, 0.2 mmol) in mixture methanol and acetonitrile (1:1) (10mL) and N-Bochomopiperazine (200 mg, 1.0 mmol) was stirred for 1 h at room temperature. The resulting mixture was concentrated to dryness and dissolved in DCM. TFA (1 mL) was added with stirring. After 30 min. mixture was concentrated and purified by prep-HPLC to provide the title compound.

##### 3,7-Di(pyrrolidine-1-yl) 1-ethyl-9-methyl-phenothiazin-5-ium iodide (PAV-174)

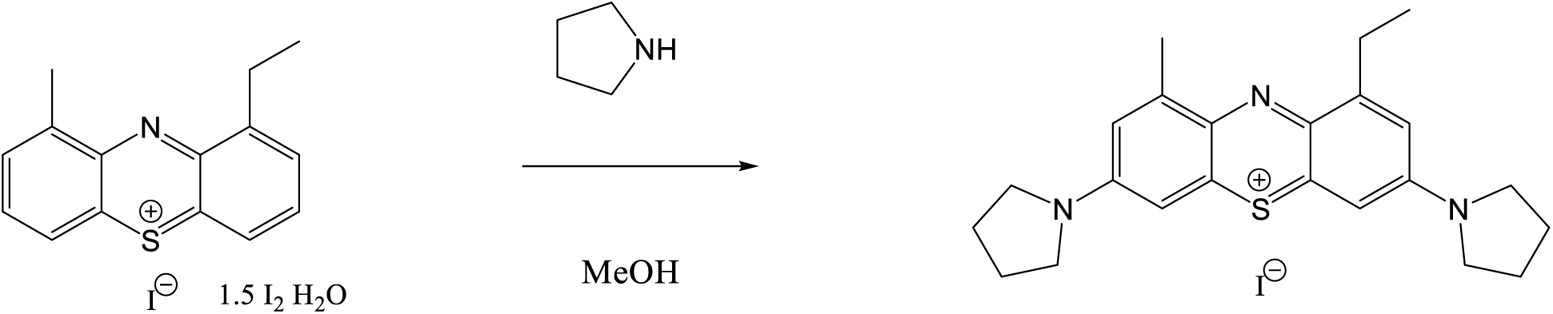

To the stirred mixture of 1-ethyl-9-methylphenothiazin-5-ium tetraiodide hydrate (383 mg, 0.5 mmol) in methanol (20mL) pyrrolidine (142 mg, 2.0 mmol) was added drop wise. The resulting mixture was stirred at room temperature 1 h, concentrated to dryness. Compound was purified with prep-HPLC.

##### 3-(Dimethylamino)-1-ethyl-9-methyl-7-(4-(trifluoromethylsulfonyl)piperazin-1-yl)phenothiazin-5-ium iodide (PAV-173)

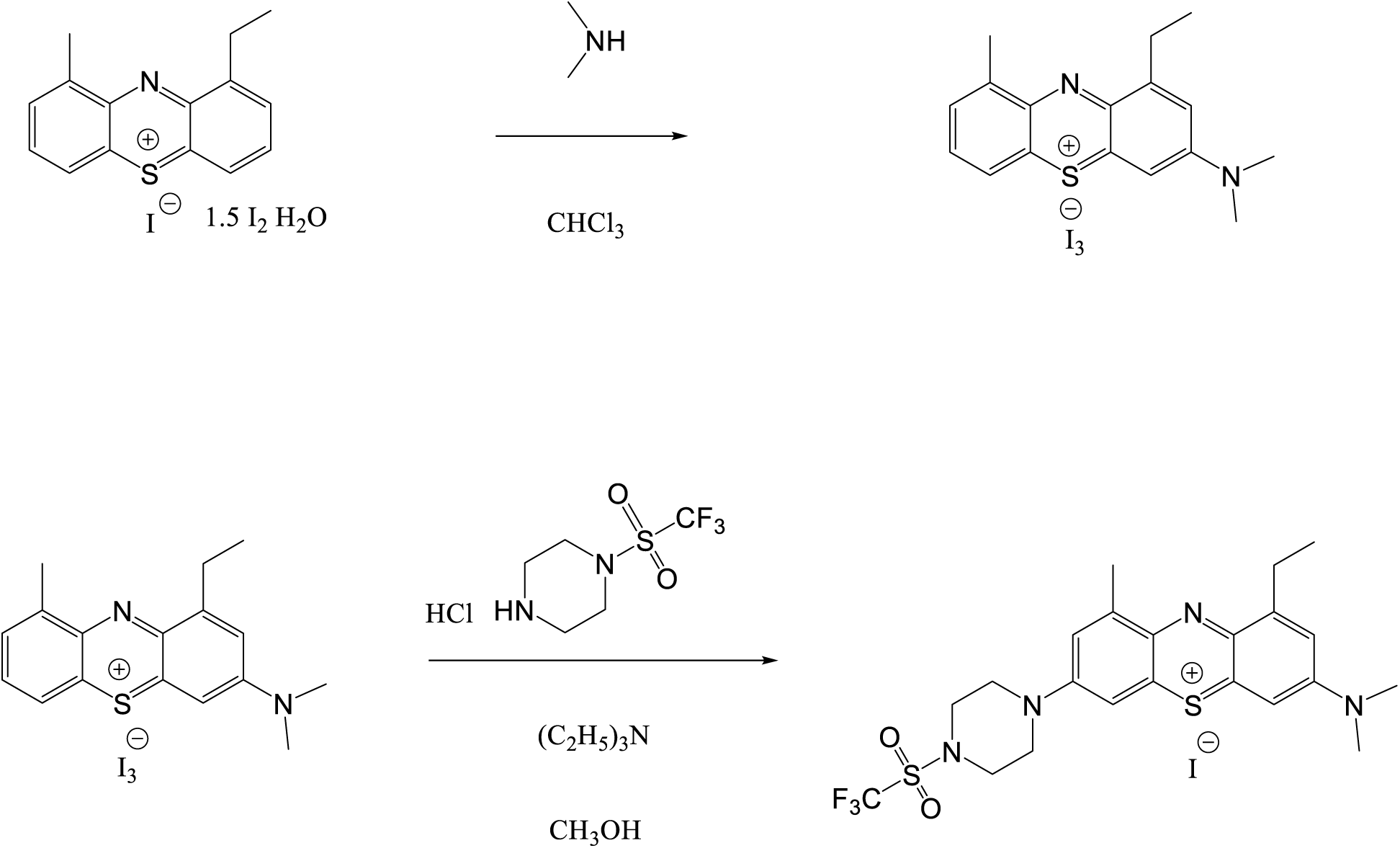

To the stirred mixture of 1-ethyl-9-methylphenothiazin-5-ium tetraiodide hydrate (383 mg, 0.5 mmol) (52) in anhydrous CHCl_3_ (20mL) dimethylamine (0.5mL, 1.0 mmol, 2M solution in THF) was added drop wise over 0.5 h. The resulting mixture was stirred at room temperature 1 h and concentrated to dryness.

A solution of 3-(dimethylamino)-1-ethyl-9-methylphenothiazin-5-ium triiodide (100 mg, 0.15 mmol) in methanol (10mL), (piperazin-1-yl)trifluoromethyl sulfone hydrochloride (115 mg, 0.45 mmol) and triethylamine (0.5mL) was stirred for 2 h at room temperature. The resulting mixture was concentrated to dryness and purified by flash chromatography using the methanol-chloroform gradient to provide the title compound.

##### 1-Ethyl-9-methyl-7-(piperidin-1-yl)-3-(4-(Bocamino)piperidin-1-yl)phenothiazin-5-ium trifluoroacetate (PAV-322)

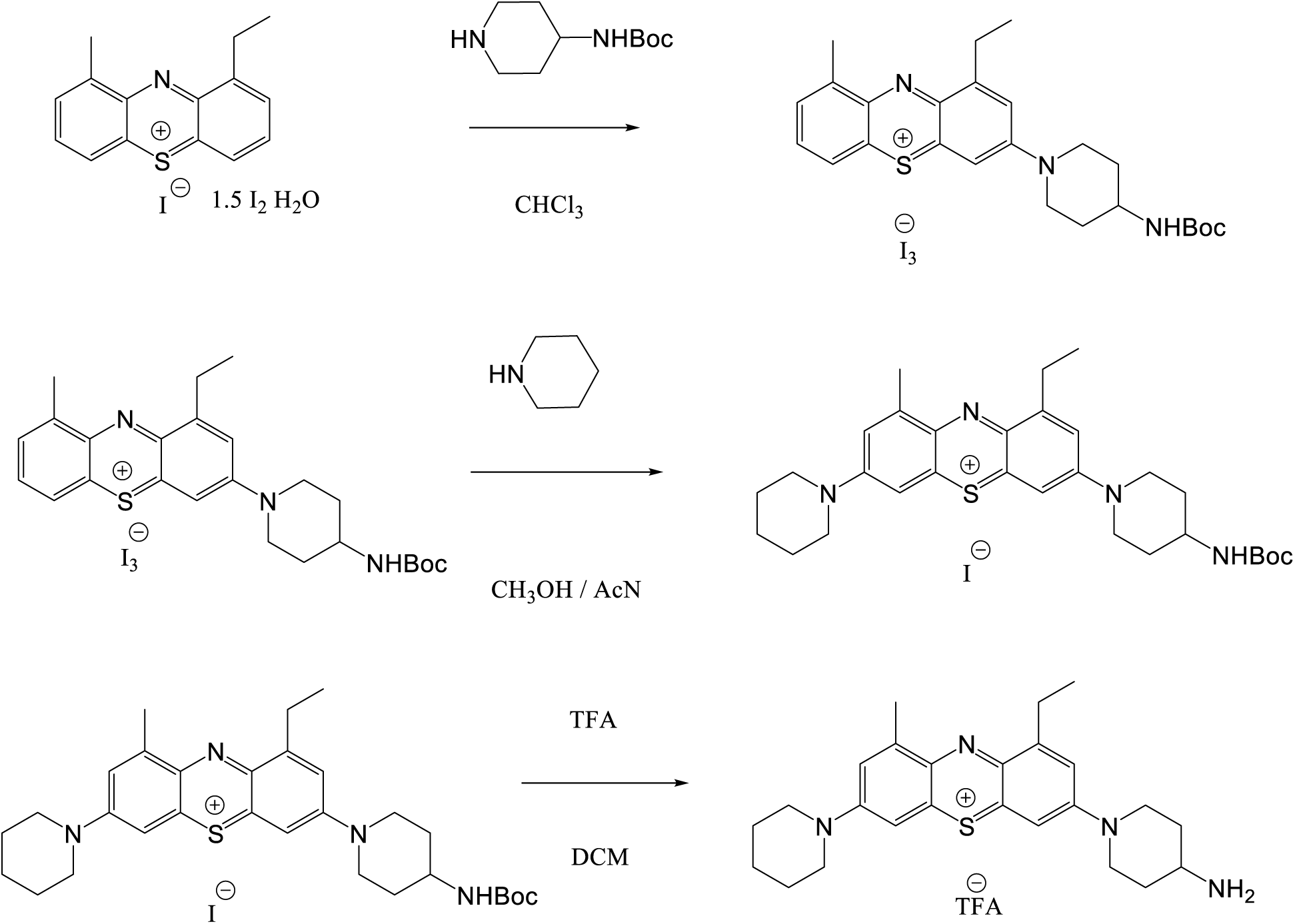

To the stirred mixture of 1-ethyl-9-methylphenothiazin-5-ium tetraiodide hydrate (153 mg, 0.2 mmol) in anhydrous CHCl3 (10mL) 4-Bocaminopiperidine (60 mg, 0.3 mmol) was added with stirring. The resulting mixture was stirred at room temperature overnight, concentrated to dryness. A solution of 1-ethyl-9-methyl-3-(4-Bocamino)piperidin-1yl)phenothiazin-5-ium triiodide (100 mg, 0.12 mmol) in mixture acetonitrile-methanol (1:1) (10mL) and piperidine (0.03 mL, 0.3 mmol) was stirred for 1 h at room temperature. The resulting mixture was concentrated to dryness and purified by flash chromatography using the methanol-chloroform gradient. Product was concentrated to dryness and dissolved in DCM. TFA (1 mL) was added with stirring. After 30 min. mixture was concentrated and purified by prep-HPLC to provide the title compound.

##### 1-Ethyl-9-methyl-7-morpholino-3-(4-Bocaminopiperidin-1-yl)phenothiazin-5-ium iodide (PAV-395)

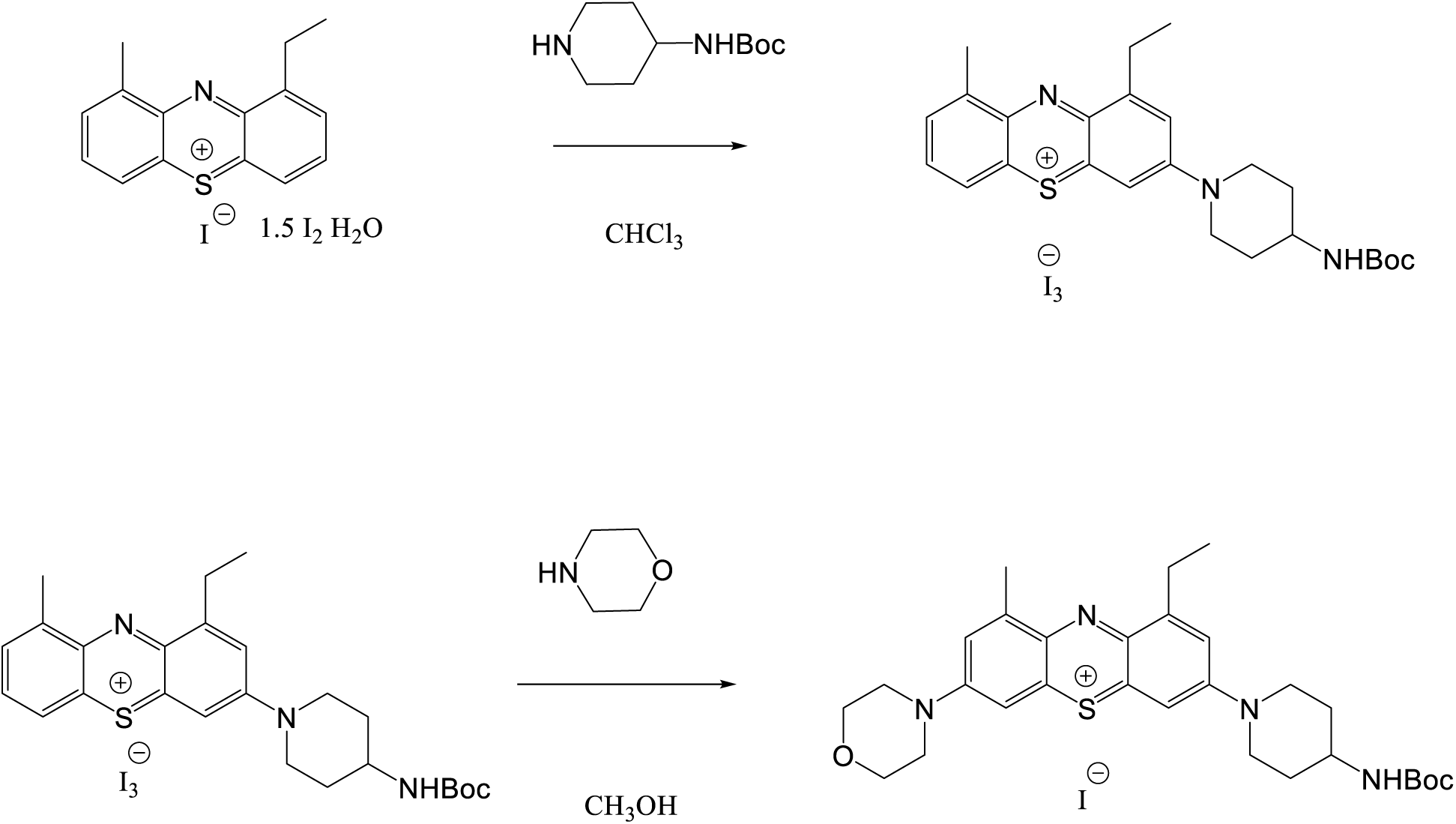

To the stirred mixture of 1-ethyl-9-methylphenothiazin-5-ium tetraiodide hydrate (153 mg, 0.2 mmol) in anhydrous CHCl_3_ (10mL) 4-Bocaminopiperidine (60 mg, 0.3 mmol) was added with stirring. The resulting mixture was stirred at room temperature overnight, concentrated to dryness. A solution of 1-ethyl-9-methyl-3-(4-Bocaminopiperidin-1yl)phenothiazin-5-ium triiodide (165 mg, 0.2 mmol) in methanol (10mL) and morpholine (17.4 mg, 0.2 mmol) was stirred for 4 h at room temperature. The resulting mixture was concentrated to dryness and purified by flash chromatography using the methanol-chloroform gradient.

##### 1-Ethyl-9-methyl-7-tiomorpholino-3-(4-Boc-1,4-diazepane-1-yl)phenothiazin-5-ium iodide (PAV-396)

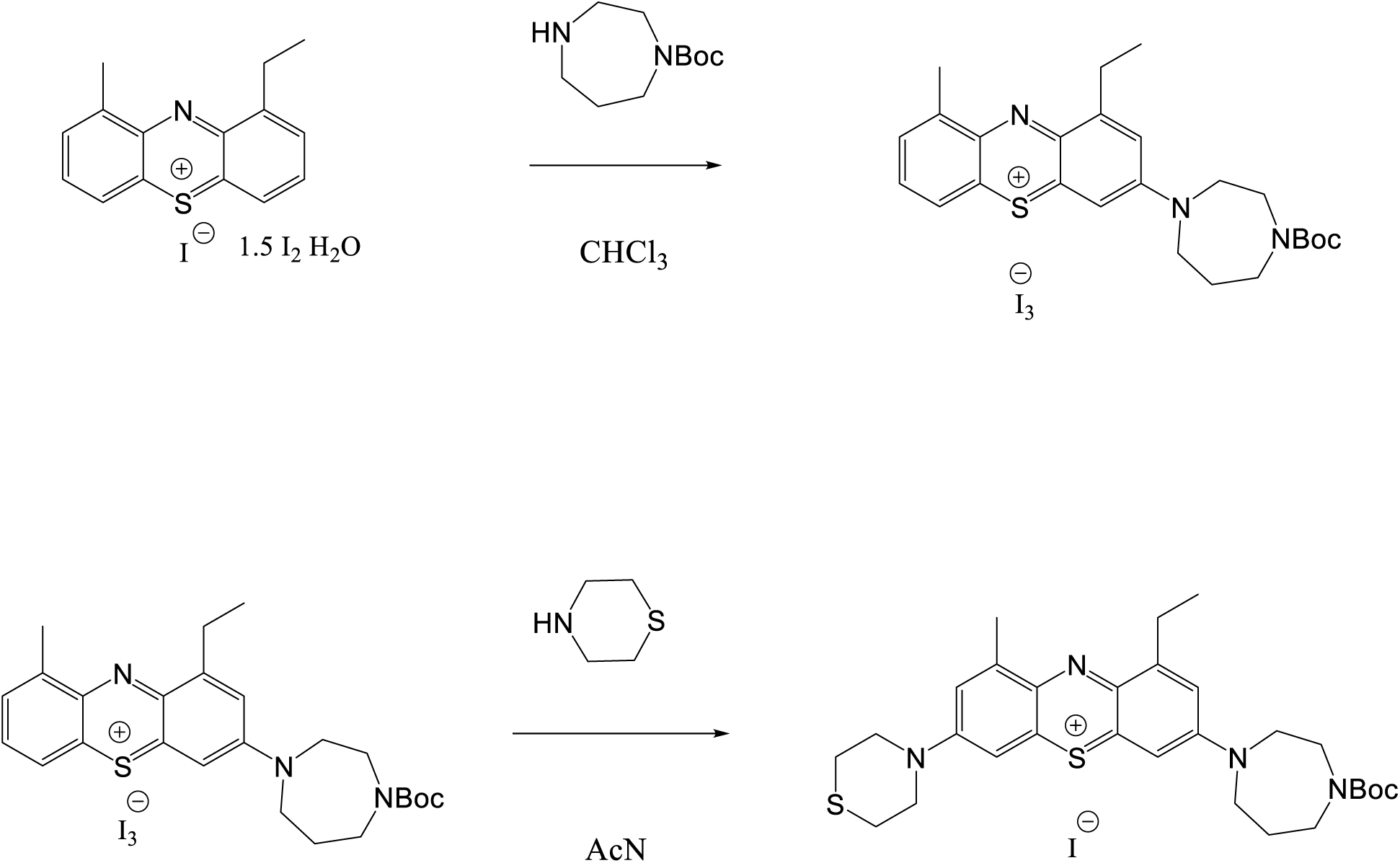

To the stirred mixture of 1-ethyl-9-methylphenothiazin-5-ium tetraiodide hydrate (780 mg, 1.0 mmol) in anhydrous CHCl_3_ (15 mL) 1-Boc-1,4-diazepane (400 mg, 2.0 mmol) was added at room temperature. The resulting mixture was stirred at this temperature for 4 h. Solvent was removed under vacuum. A solution of 1-ethyl-9-methyl-3-(4-Boc-1,4-diazepane-1-yl)phenothiazin-5-ium triiodide (165 mg, 0.2 mmol) in acetonitrile (10mL) and tiomorpholine (72 mg, 0.8 mmol) was stirred for 4 h at room temperature. The resulting mixture was concentrated to dryness and purified by flash chromatography using the methanol-chloroform gradient. Structure of compounds PAV-173, PAV-322, PAV-395 and PAV-396 was confirmed by stereospecific synthesis by the method described below for PAV-645.

##### 1,9-Diethylphenothiazin-5-ium tetraiodide hydrate

The same scheme and procedures like for 1-ethyl-9-methylphenothiazinium salt. Compound 3 is 2-ethylbromobenzene.

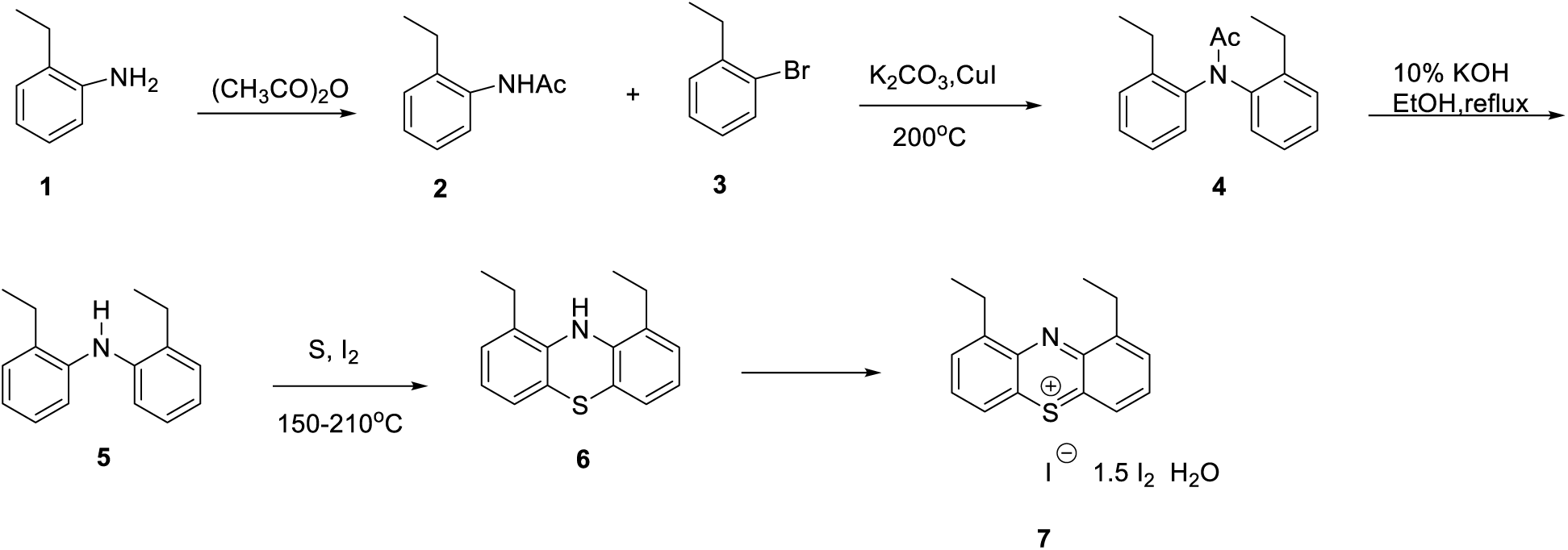

##### 1,9-Diethyl-3-(1,4-diazepane-1-yl)-7-dimethylaminophenothiazin-5-ium iodide (PAV-493)

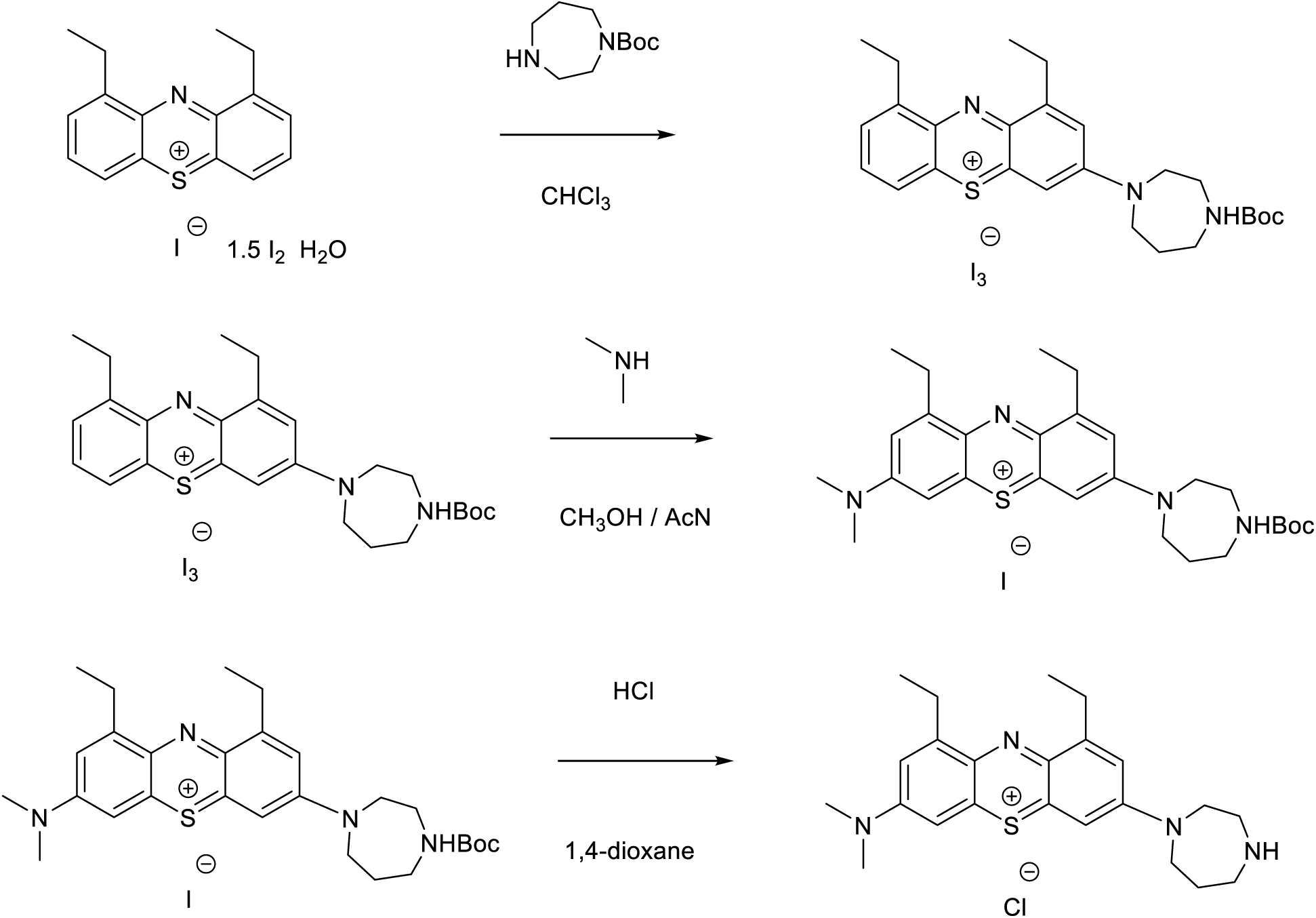

To the stirred mixture of 1,9-diethylphenothiazin-5-ium tetraiodide hydrate (3.9 g, 5.0 mmol) in anhydrous CHCl3 (100 mL) 1-Boc-1,4-diazepane (1.2 g, 6.0 mmol) was added at room temperature. The resulting mixture was stirred at this temperature for 2 h. Solvent was removed under vacuum. A solution of 1,9-diethyl-3-(4-Boc-1,4-diazepane-1-yl)phenothiazin-5-ium triiodide in mixture methanol-acetonitrile (1:1) (150mL) and dimethylamine (10 mL 2 M sol. in THF) was stirred for 1 h at room temperature. The resulting mixture was concentrated to dryness and purified by flash chromatography using the methanol-chloroform gradient. Product (300 mg) was concentrated to dryness. HCl (5 mL 4 M solution in 1,4-dioxane) was added with stirring. After 30 min. mixture was concentrated and purified by prep-HPLC to provide the title compound.

##### 1-Ethyl-7-(piperazin-1-yl)-3-diethylaminophenothiazin-5-ium chloride (PAV-645)

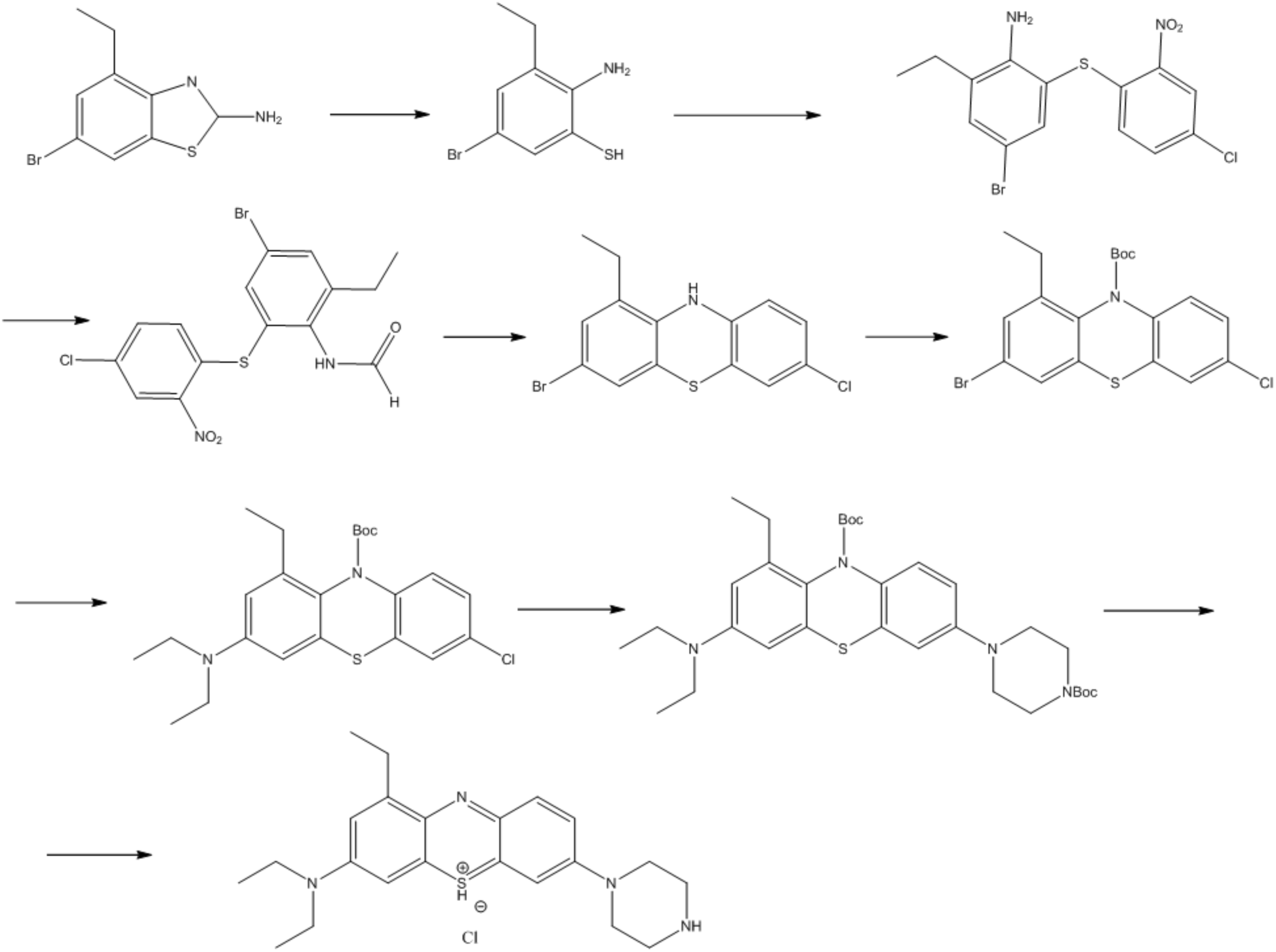

2-Amino-5-bromo-3-ethyl-benzenethiol: 6-Bromo-4-ethyl-1,3-benzothiazol-2-amine (2.0 g, 7.7 mmol) was added to a solution of KOH (13.5 g, 240 mmol) and H2O (25 mL) and the reaction mixture was heated to 150oC overnight. The reaction was monitored by LCMS for consumption of starting material. At the completion of the reaction, the reaction was allowed to cool to RT and ice bath was added as the mixture was slowly neutralized with conc. HCl to pH= 6. The solid was filtered off and dried under vacuum overnight. Both the solid and the filtrate were washed with Et2O and the organic layers were combined. The resulting organic layer was washed with brine, dried over MgSO4, filtered and evaporated to give the product, which was used without further purification. LCMS, M+H=232.0.

4-Bromo-2-(4-chloro-2-nitro-phenyl)sulfanyl-6-ethyl-aniline: 2-Amino-5-bromo-3-ethylbenzenethiol (2.3 g, 10 mmol) was combined with 1,4-dichloro-2-nitrobenzene (2.03 g, 10.6 mmol), Cs2CO3 (10.29 g, 31.6 mmol) and acetonitrile (50mL) and stirred at room temperature overnight. The starting material was monitored by TLC. At the completion of the reaction, the reaction was filtered, and the solvent evaporated to give a residue. The residue was purified by flash silica gel chromatography to give the desired product. LCMS: M+H=388 N-[4-Bromo-2-(4-chloro-2-nitro-phenyl)sulfanyl-6-ethyl-phenyl]formamide: 4-Bromo-2-ethyl-6-(4-chloro-2-nitro-phenyl)sulfanyl-aniline (14.6 g, 39.22 mmol) was dissolved in formic acid (50mL) and heated at 100oC overnight. The reaction was allowed to cool to RT and ice water was added, keeping the temperature near 0oC. The resulting solid was filtered off, washed with cold water and dried overnight under vacuum. LCMS, M+H=416

3-Bromo-7-chloro-1-ethyl-10H-phenothiazine: N-[4-Bromo-2-ethyl-6-(4-chloro-2-nitrophenyl) sulfanyl-phenyl]formamide was combined with acetone (20 mL) and heated to reflux and an alcoholic solution of KOH (0.7 g,12.4 mmol) in EtOH (20ml) was added. The resulting solution was refluxed for 30 minutes to 1 hour. Another portion of alcoholic KOH (0.7 g,12.4 mmol) was added and the resulting reaction was refluxed for 4 hours. The mixture was allowed to cool to room temperature. The solvent was evaporated, and the residue was extracted with CHCl3 and brine. The combined organic layers were dried over MgSO4, filtered, and evaporated to give a residue. The crude material was purified by flash silica gel chromatography to give the desired compound. LCMS, M+H=341 tert-Butyl 3-bromo-7-chloro-1-ethyl-phenothiazine-10-carboxylate: 3-Bromo-1-ethyl-7-chloro-10H-phenothiazine (1.5 g, 4.40 mmol), was combined with Boc2O (1.92 g, 8.81 mmol), DMAP (0.54 g, 4.40 mmol), and acetonitrile (30 mL). The resulting reaction mixture was stirred and, heated to reflux overnight. After cooling the reaction mixture was concentrated to dryness and purified on the ISCO using EtOAc / Hexanes gradient to afford the desired compound as a waxy solid. M+H =441tert-Butyl 7-chloro-3-diethylamino-1-ethylphenothiazine-10-carboxylate: To a mixture of Na t-OBu (23 mg, 0.237mmol), Pd(dba)2 (2.3 mg, 0.004mmol), BINAP (2.5 mg, 0.004mmol), diethylamine (17 mg, 0.237mmol) and the tert-butyl 3-bromo-7-chloro-phenothiazine-10-carboxylate (85 mg, 0.206mmol) was added dioxane (2.5mL) (all in a flame dried screw top vial). The mixture was stirred, under argon, at 100oC for 4h (take an aliquot after 2h and check for completeness). Upon completion the reaction was cooled diluted with dioxane (5mL) and filtered through a small celite plug. The filtrate was rotary evaporated to dryness and the residue purified by the flash-chromatography. Yield 75mg (91%) LCMS, M+H=433.

tert-butyl 7-(N-Bocpiperazin-1-yl)-3-diethylamino-1-ethylphenothiazine-10-carboxylate: To a mixture of Na t-OBu (23 mg, 0.237mmol), PdRuPhos (3 mg, 0.004 mmol), RuPhos (2 mg, 0.004 mmol), N-Bocpiperazine (47 mg, 0.237mmol) and tert-Butyl 7-chloro-3-diethylamino-1-ethylphenothiazine-10-carboxylate (83mg, 0.2mmol) was added THF (2.5mL) (all in a flame dried screw top vial). The mixture was stirred under argon at 85oC for 4h (take an aliquot after 2h and check for completeness). Upon completion the reaction was cooled diluted with 10ml of THF and filtered through a small celite plug. The filtrate was rotary evaporated to dryness and the residue purified on the ISCO. Yield 72mg (80%) LCMS, M+H=583.

3-Diethylamino-7-(piperazin-1-yl)-1-ethylphenothiazin-5-ium: A 10mg sample of tert-butyl 3-diethylamino-7-(piperazin-1-yl)-1-ethylphenothiazine-10-carboxylate was treated with 200ul of 4N HCl in dioxane for 1h with stirring. The mixture was then rotary evaporated to dryness. Yield=7.5 mg as HCl salt. LCMS, M+H=418

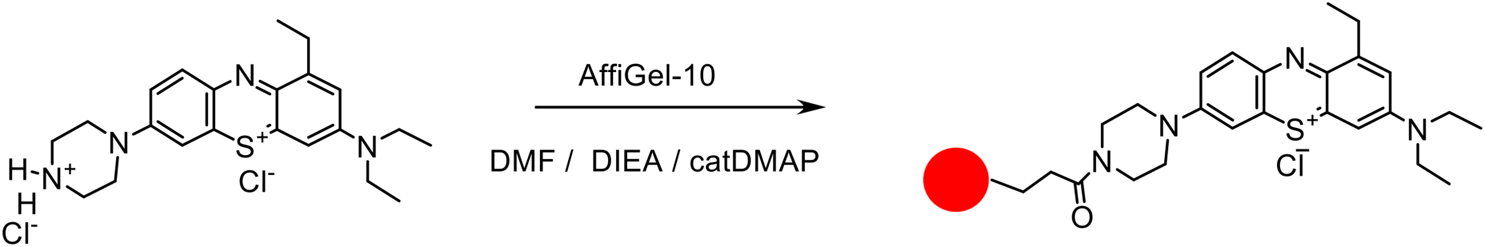

A solution of 3-Diethylamino-7-(piperazin-1-yl)-1-ethylphenothiazin-5-ium chloride (13,6 mg, 0.03 mmol), DIEA (60 µL, 034 mmol) and DMAP (2 mg) in 2 mL of DMF was added to 3 mL of AffiGel-10 resin (Bio-Rad) in an Econo-Pac Chromatography Column. After agitating overnight at room temperature the resin was washed 3x with 4 mL of DMF and then stored under 3 mL of i-propanol.

PAV-802 is methylene blue, a commercially available compounds which was purchased from Aldrich fine chemicals, Inc.

### Synthesis of 3,7-Di(pyrrolidine-1-yl)phenothiazine-5-ium iodide (PAV-617)

#### Phenothiazin-5-ium tetraiodide hydrate

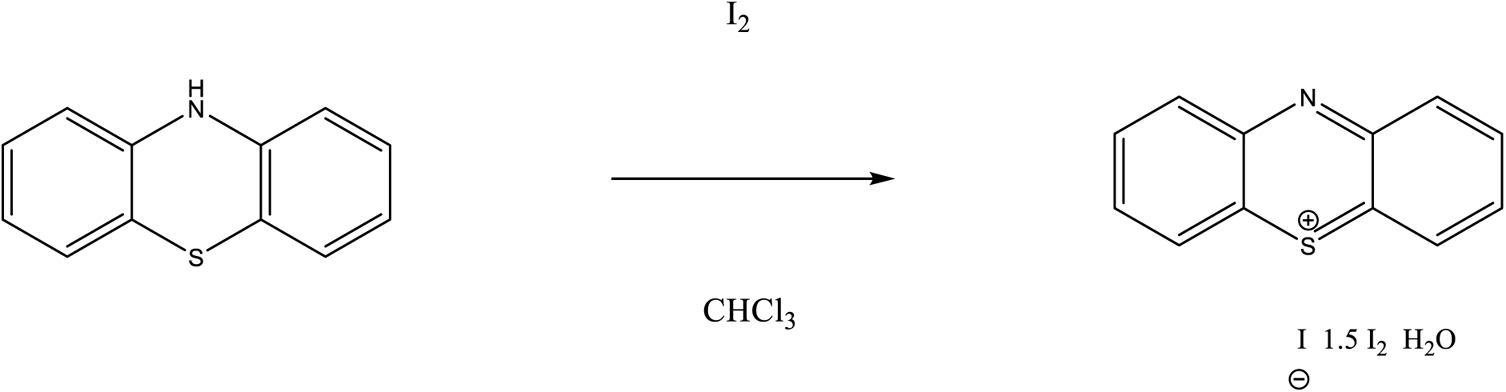

Phenothiazin-5-ium tetraiodide hydrate: a solution of phenothiazine (4.98 g, 25 mmol) in anhydrous chloroform (50 ml) was stirred at 5^0^C and the solution of iodine (12.7 g, 50 mmol) in CHCl_3_ (250 ml) was added dropwise over 4h. The resulting dark solution was stirred for an additional 3h at 5^0^C, monitored by TLC. After the disappearance of the starting material, the resulting precipitate was filtered, washed with a copious amount of chloroform, dried overnight in vacuo to afford a dark solid (13.9 g, 74%).

#### 3,7-Di(pyrrolidine-1-yl)phenothiazine-5-ium iodide

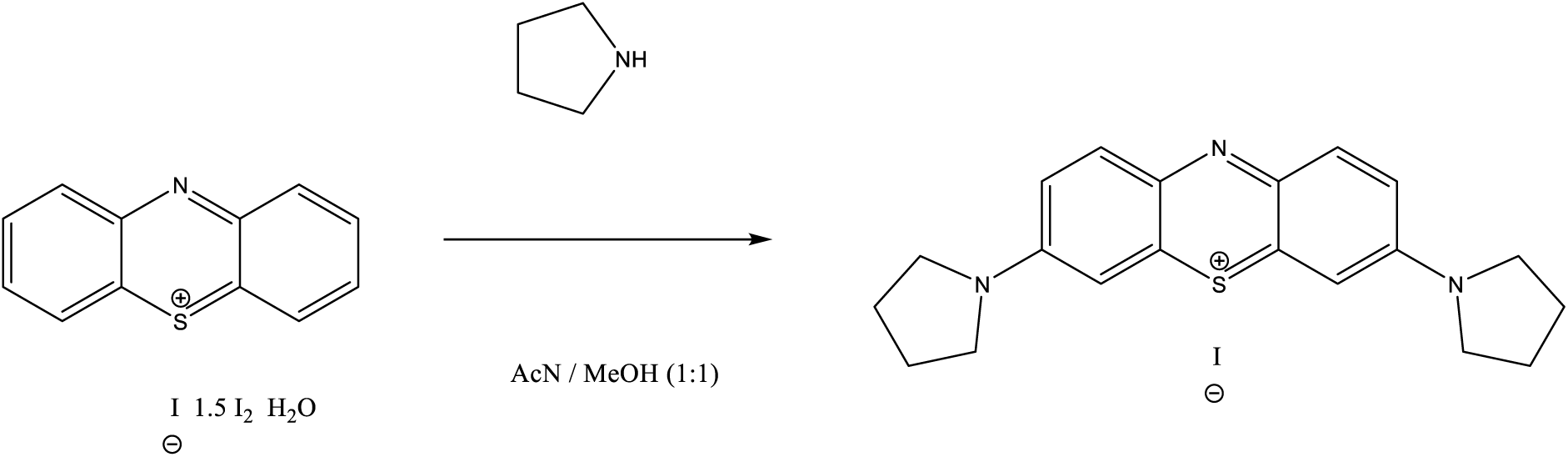

A solution of phenothiazin-5-ium tetraiodide hydrate (2.8 g, 3.6 mmol) in mixture acetonitrile/methanol (50 ml) and pyrrolidine (710 mg, 10 mmol) was stirred for 4 h at room temperature. The resulting mixture was concentrated to dryness and purified by flash chromatography using the methanol-chloroform gradient to provide the title compound.

### Generation of recombinant vectors

For the generation of a tau-P301S expression plasmid the multiple cloning site (MCS) of the pLHCX vector (TaKaRa) was modified by inserting a synthetic MCS-sequence (ACCGGTCTCGAGGCGGCCGCGGCCAAAAAGGCCGGATCCGTTAACACCAAAAA ATGGCACGTGGCCGGCACGCGTGGGCCCGTCGAC). For this the modified_MCS_fw and modified_MCS_rev Oligonucleotides were annealed and cut with FastDigest HindIII and ClaI (both Thermo Fisher Scientific) and then inserted into pLHCX that was opened with HindIII and ClaI and dephophorylized with FastAP (Thermo Fisher Scientific). The ligation product (pLHCX-mod-MCS) was transformed and amplified in DH5α (Thermo Fisher Scientific). The ORF of human Tau 2N4R including the complete 3’-UTR was ligated into pLHCX-mod-MCS via XhoI and BamHI. Then the P301S mutation was introduced by site-directed mutagenesis (CCG>TCG) using the QuikChange Site-directed Mutagenesis Kit (Stratagene) and the oligonucleotides Tau5 P301S FOR and Tau6 P301S REV.

### Generation of retroviruses

Retroviruses were produced in GP2-293 cells (Clontech). A confluent 10 cm dish with GP2-293 cells were split 1:4 and seeded into T75 cell culture flasks that were coated with Poly-Ornithin (Sigma). At the next day, when the cells reached about 70% confluency, the cells were transfected with 9.75 µg pLHCX-tau-P301S, 9.75 µg pVSV-G (Addgene) and 58.5 µL Gene Juice diluted in Optimem (Gibco) and incubated for 20 min. After changing the medium of the GP2-293 cells the Gene Juice:DNA complex was added to the cells in a drop-wise manner. The cells were incubated for 48h. Then the medium was carefully removed and replaced with 6.5 mL of fresh medium. The retroviral particles were collected after 24h, passed through a 0.45 µm filter (VWR) and aliquots were stored at −20°C until use.

### Generation of stable cell lines

SH-SY5Y cells constitutively expressing human tau 2N4R-P301S were generated by retroviral infection with pLHCX-mod-MCS-tau-P301S. 1×10^6^ SH-SY5Y cells were seeded on 6-well plates. 1 mL of the retroviral supernatant was quickly thawed in a 37°C water bath and mixed with 1mL of fresh DMEM/F12 medium and 10 µg polybrene. After incubation of 24h cells were washed with PBS and fresh medium was added. At the next day cells were splitted 1:10 into a T75 flask containing DMEM/F12 and 400 µg/mL hygromycin B (Thermo Fisher Scientific). The selection medium was changed every 2-3 days until stable cell clones appeared.

### Generation of SH-SY5Y-tau-P301S MIF knockout line by CRISPR-Cas9

For generation of the SH-SY5Y-tau-P301S MIF knockout line, we identified an sgRNA sequence specific for human MIF (exon 1) using the MIT-CRISPR server (http://crispr.mit.edu/): AGCTCGGAGAGGAACCCGTC. As negative control we used a scrambled sgRNA sequence: GCACTCACATCGCTACATCA (Applied Biological Materials, Richmond, BC, Canada). MIF-ko-5:CACCGAGCTCGGAGAGGAACCCGTC and MIF-ko-3: AAACGACGGGTTCCTCTCCGAGCTC as well as scrambled-5: CACCGGCACTCACATCGCTACATCA and scrambled-3: AAACTGATGTAGCGATGTGAGTGCC were phosphorylated with T4 PNK (Thermo Fisher Scientific) in T4 ligation buffer (Thermo Fisher Scientific). After annealing each set of oligonucleotides they were ligated into the lentiCRISPRv2 shuttle vector (Addgene) that was linearized with BsmBI (NEB) and dephosphorylated with FastAP (Thermo Fisher Scientific). Finally, the product was transformed into Stbl3 bacteria (Invitrogen). The cloning products were validated by sequencing.

Lentiviral particles were produced in 90-100% confluent HEK293-FT cells (Clontech) in a T75 flask. The cells were washed once with PBS and 5 mL Optimem (Invitrogen) were added 1h before transfection. The transfection mixture was prepared in 667 µL Optimem and contained 7 µg of the lentiviral shuttle vector as well as 4 µg pMDLG, 4 µg pRSV_Rev, and 4 µg pMD2.G for packaging. In another tube, 640 µL Optimem were supplemented with 27 µL Lipofectamine 2000 (Thermo Fisher Scientific), and both mixtures were incubated for 5 min at RT. The DNA was then combined with the Lipofectamine-containing medium, followed by 20 min incubation at RT before it was added to the cells in a drop wise manner. After 14h the medium was changed to 6 mL of normal growth medium (DMEM). Following another 24h of incubation the cell culture supernatant containing the viral particles was passed through a sterile 0.45 µm filter (VWR) to remove cell debris. Aliquots of 1 mL thereof were stored at −20°C.

For infection with lentiviral particles SH-SY5Y-tau-P301S cells were seeded into T25 flasks. At the next day the medium was replaced with 1 mL fresh complete growth medium, 1 mL of the viral particles and 10 µg polybrene (Sigma). One flask was left uninfected for selection control and was provided with 2 mL polybrene-containing growth medium instead. After incubation for 24h the medium was removed and cells were washed twice with PBS. Then 6 mL of fresh growth medium were added and cells incubated for another 24h. For selection the cells were trypsinized and resuspended in 10 mL growth medium containing 1 µg/mL of the selection marker puromycin and transferred to a new T75 flask. The medium was changed every 3-4 days and selection marker was added until stable cell clones appeared and non-infected control cells completely died. The knockout efficiency of MIF was analyzed by Western Blot.

### HSV-1 KOS viral stock generation

The HSV-1 KOS strain (ATCC-VR-1493) with 2×10^7^ plaque forming units (pfu)/mL was used to prepare viral stocks after infection of Vero cells (ATCC-CCL-81). A confluent 75-cm^2^ flask with Vero cells was infected with HSV-1 KOS at a multiplicity of infection (MOI) of 0.01 pfu/cell. For this the virus was diluted in 3 mL of Medium 199 (Gibco). After incubation for 2 h at 37°C the medium was aspirated and 3 mL of fresh DMEM (Gibco) supplemented with 5% New Born Calf Serum (NBCS; Gibco) was added to the flask. The virus was allowed to replicate for 3 days. Then the virus was released by freeze thawing following 3x sonication for 30 sec using a Misonix ultrasonic water bath at 100% amplitude. Aliquots of the virus were stored in liquid N_2_. The virus titer was determined by virus plaque assay.

### Virus plaque assay

Virus titer determination was carried out in 25-cm^2^ flasks or 6-well plates with confluent Vero cells. The virus stock was serially diluted in Medium 199. 1 mL of each of the dilutions were added to the Vero cells in duplicates and incubated for 2 h. Then the inoculum was aspirated and DMEM supplemented with 5% NBCS and 7.5 µg/mL human immunoglobulin (Sigma) was added. After two days the cells were washed twice with PBS and fixed with MeOH for 5 min at RT. Then 1:10 diluted KaryoMax Giemsa stain solution (Gibco) was added to the cells and incubated for 20 min at RT. Following washing with water the plaques were counted under the microscope and the titer calculated.

### Infection of cell lines with HSV-1

4×10^5^ Vero cells, 8×10^5^ SH-SY5Y or 2×10^6^ differentiated LUHMES (differentiation day 2) cells were seeded into 12-well plates. After 24h the cells were treated with compounds (as indicated in the figures) and after 1h they were infected with HSV-1 applying MOIs as described in the figures. After 20h (or at indicated time points) cells were either washed twice with PBS, then lysed in cold PBS/1% NP40 including the cOmplete EDTA-free protease inhibitor cocktail (Roche) and phosphatase inhibitor cocktail (Sigma) or virus was released by freeze-thawing and ultrasonic treatment for analysis in viral plaque assay (see above).

### In-cell ELISA

Antiviral activity of the compounds was measured by in-cell ELISA that was adapted from Fabiani et al. (Fabiani et al., 2017). 4×10^4^ Vero cells were seeded in 96-well plates and grown to confluence for 24h. Then, the medium was removed and cells were treated with serially diluted compounds or DMSO (Sigma) added to 50 µL of Medium 199. Thereafter, the cells were infected with 50 µL of HSV-1 (MOI of 1) diluted in Medium 199. In parallel cells were infected with serial dilutions of HSV-1 to generate a standard curve. After incubation for 2 h the wells were washed twice with PBS and then 100 µL DMEM plus compound or DMSO were added and the cells were incubated for 24h at 37°C. At the next day, the cells were washed twice with PBS, fixed with 50 µL of PBS/4% paraformaldehyde (PFA) for 15 min and then permeabilized with 50 µL of PBS/0.1% TX-100 (Sigma) for 5 min at RT. Following blocking with 5% BSA in PBS/0,05% Tween-20 (Sigma, PBS-T) for 60 min the cells were incubated with 50 µL of anti-HSV-1-gC antibody (abcam) diluted 1 to 10,000 in PBS-T over night at 4°C. The following day, cells were washed four times with PBS-T and then incubated with Hrp-conjugated goat anti-mouse antibody (Thermo Fisher Scientific) at a dilution of 1:10,000 in PBS-T for 60 min. Then, cells were washed four times with PBS-T before adding 50 µL of the substrate cocktail (OptEIA, BD). The reaction was stopped after 30 min by adding 50 µL of 2N H_2_SO_4_, and signals were spectrometrically analyzed using a Tecan Safire at 450 nm.

### Infection of human brain organoids

60d old organoids with an estimated cell number of 2.5×10^6^ were preincubated with DMSO or 250 nM of PAV-174 for 1h (final DMSO concentration was 0.5%). Then organoids were infected with HSV-1 at a MOI of 0.1 or 1.0. At the next day the medium was renewed and new compound was added. After another 24h, the organoids were washed two times with PBS and then either fixed with 4% PFA at 4°C overnight (for immunocytochemistry and TUNEL assay) or lysed with cold RIPA buffer (50 mM Tris-HCL, pH 7.4, 1% NP-40, 0.5% Na-deoxycholate, 0.1% SDS, 150 mM NaCl, 5 mM EDTA, 50 mM NaF) plus 1X Protease inhibitor cocktail (Roche) and phosphatase inhibitor cocktail (Sigma) followed by 4x sonication for 30s using a Misonix ultrasonic water bath at 100% amplitude. Protein concentration was determined with the DC Protein Assay (Bio-Rad). For immunocytochemistry and TUNEL assay fixed organoids were embedded in paraffin and section of 5 µm were prepared.

### TUNEL assay

Apoptotic cells in sections of brain organoids were detected by TdT-mediated dUTP-biotin nick end labeling (TUNEL) using the DeadEnd Fluorometric TUNEL System (Promega, G3250) according to the manufacturers recommendations. At least three 5-µm sections of the brain organoids were analyzed by taking at least 10 images of each organoid. Dead cells were counted using Fiji/ ImageJ (NIH).

### Tau assay

For the analysis of compound effects on tau phosphorylation 4×10^5^ SH-SY5Y tau-P301S cells were seeded on 24-well plates. At the next day the cells were treated with compounds diluted in DMSO. Final concentration of DMSO was 0.2%. After an incubation at 37°C for 48h the cells were washed twice with PBS and then lysed in PBS/1% NP40 (Applichem) including the cOmplete protease inhibitor cocktail 2 (Roche) and phosphatase inhibitor cocktail (Sigma). The protein concentration of the samples was determined with the DC Protein Assay (Bio-Rad).

### Immunoblot

After the protein concentration of cell lysates and homogenates had been determined using the DC Protein Assay Kit (Bio-Rad), 20 µg of each sample were dissolved in NuPAGE LDS Sample buffer (including 2% β-mercaptoethanol), boiled at 100°C for 5 min and then separated on NuPAGE Novex 4-12% Bis-Tris midi gels (Invitrogen) using NuPAGE MES SDS Running buffer (Invitrogen). Proteins were then transferred to a 0.2 µm nitrocellulose membrane (Amersham) in 24 mM Tris/192 mM glycine/ 20% MeOH at 400 mA for 2h using a tank blot system (Bio-Rad). In order to detect any blotting artifacts, the proteins on the membranes were visualized with Ponceau S (Sigma). The membranes were destained with PBS and then blocked with PBS/ 5% nonfat milk (Oxoid) for 1h at RT. Incubations with primary antibodies (diluted 1:1,000) in PBS-T were carried out overnight at 4°C. Then the membranes were washed three times with PBS-T for 10 min and afterwards incubated with fluorescent secondary antibodies (IR Dye 680 or 800: Li-COR Biosciences.) diluted 1:20,000 in PBS-T for 1h at RT. Following three washing steps with PBS-T for 10 min each, the membranes were scanned using the LI-COR Odyssey CLX. Signal intensities were calculated using the Image Studio Version 2.1 software (LI-COR Biosciences). Blots were reused for staining with additional primary antibodies following the same procedure.

### Sandwich ELISA

For quantitative determination of oxidized MIF (oxMIF), 96-well-Immuno Maxisorp plates (Nunc) were coated with 0.5 µg/ well of 10C3 in 50 mM carbonate buffer pH 9.4 to capture total MIF species. The wells were blocked with PBS/5% BSA/ 0.1% Tween-20 for 2h at RT. After washing with PBS-T, samples, either 25 µg brain homogenate or 20 µl of cell culture supernatant were diluted in 100 µL PBS/1% BSA/ 0.1% Tween-20, were added and incubated overnight at 4°C. The wells were washed 3x with PBS-T and incubated with the oxMIF specific antibody Imalumab at a 1:2,000 dilution for 4h at RT. Then, the wells were again washed 3x with PBST before a goat anti-human IgG-HrP conjugate (Southern Biotech; 1:5,000 dilution) was added for 1h at RT. After 4x washing with PBS-T the ELISA was developed with 1-step Ultra TMB ELISA (Thermo Scientific). For generation of an oxMIF standard curve, recombinant wt-MIF was treated with 0.2% Proclin300 (Sigma) that transforms recombinant MIF into an oxMIF surrogate as described in (Thiele et al., 2015).

### MTT assay

For viability analysis 4×10^4^ Vero or 8×10^4^ SH-SY5Y-tau-P301S cells were plated in a 96-well plate and incubated at 37°C overnight. Then the compounds diluted in DMSO were added to the cells. The final concentration of DMSO was 1%. The cells were then incubated for 24h (Vero) or 48h (SH-SY5Y). Afterwards 20 µL of a Thiazolyl Blue Tetrazolium Bromide (MTT, Sigma) solution (5 mg/mL in PBS) was added to the cells and incubated for 4h allowing viable cells to reduce the yellow MTT into blue formazan metabolites. Following aspiration of the medium, the formazan was resuspended in 200 µL isopropanol/ 40 mM HCl and incubated for 30 min at RT. The diluted formazan was spectrometrically analyzed using a Tecan Safire Spectrometer at 560 nm and background at 670 nm was subtracted.

### Immunocytochemistry

For immunocytochemistry analysis 5×10^4^ LUHMES cells at differentiation day 2 were seeded on 13 mm coverslips that were coated with L-ornithine and fibronectine (see above) and placed into 24-well plates. After four additional days of differentiation the cells were infected with HSV-1 (MOI = 1). After 24h the cells were carefully washed with PBS and then fixed with PBS/4% PFA for 15 min at RT. The fixed cells were then permeabilized and blocked with 1%BSA (Sigma) /0.5% saponin (Sigma) and 5% (w/v) nonfat milk (Oxoid) for 1h at RT. Anti-HSV-1-gC diluted 1:500 in 1%BSA/0.5% saponin was applied overnight at 4°C. At the next day the cells were washed three times with PBS and then incubated with anti-mouse Alexa-fluor 488 (Invitrogen) diluted 1:1,000 in PBS for 1h at RT. Following washing three times with PBS and two times with water, cells were mounted with ProLong Gold with DAPI (Invitrogen), and imaged with a Zeiss Axiovision Apotome.2 microscope.

For immunocytochemistry of organoid sections, frozen slides were thawed and kept on RT until they were dried. Slides were washed with PBS and incubated for 5 min. Then the sections were incubated with 30 mM glycine in PBS for 5 min and permeabilized with PBS/ 0,5% TX100/0,1% Tween-20 for 10 min. To equally distribute the solutions a parafilm layer was placed on top. Following permeabilization the sections were blocked with PBS/ 0,5% fish gelatin/0,1% TX-100 for 2h at RT. Anti-HSV-1-gC was diluted in blocking solution (1:500) and incubated at 4°C overnight. At the next day the sections were washed three times with blocking solution before adding the secondary antibody (Anti-Mouse IgG-Alexa Fluor 594) diluted 1:500 in blocking solution for 4h at RT. Again, sections were washed three times with blocking solution and the anti-tubulin antibody (TUJ1, 1:500) was added and incubated over night at 4°C. The following day sections were washed three times with blocking solution before they were incubated with anti-Rabbit IgG-Alexa Fluor 647 (1:500) for 4h at RT. Finally, sections were washed two times with PBS and two times with water before mounting with Mowiol (Sigma). Raw data were collected using a Leica SP8 confocal system.

### Drug affinity chromatography (DRAC) and proteomics

Pigs were raised in Struve Labs, Manning, Iowa with proper food and free movement as per their IACUC protocol. They were euthanized and brains were removed immediately and homogenized in PBB containing 10 mM HEPES, 10 mM NaCl, 1 mM Mg, and 0.35% TX-100. 10% human brain homogenates were prepared in cold PBB and aliquots were flash frozen in liquid N_2_. Homogenates were centrifuged at 10,000g for 10 min and supernatants were collected. Cells were harvested by scraping into cold PBS and pelleted at 3,000g for 10 min at 4°C. The pellet was then resuspended in PBB yielding a concentration of approx. 5 mg/mL. The lysates were cleared by centrifugation and the supernatants were flash frozen.

For drug affinity chromatography (DRAC), supernatant of lysates or brain extracts were supplemented with 1 mM of ribonucleoside triphosphates (ATP, GTP, CTP, UTP; NEB), 10 mM creatine phosphate (Sigma) and 10 µL/mL of a 5 mg/mL stock solution of creatine kinase (Sigma). The samples were then loaded onto 20 µL of PAV-645 resin that were equilibrated with PBB-buffer (plus supplements). After one hour of incubation at 26°C, the beads were washed with PBB (100x bed volume). Bound proteins were then eluted either with 200 µM PAV-645 (pig brain) or 100 µM PAV-174 (human brain samples) in the same buffer used as competitor solution or urea (cell lysates) for 1h. All elutions were kept frozen until further analysis.

For Mass spectrometric identification, pig brain eluates were run on a freshly prepared 12% SDS gel made up with all filtered solutions and chemicals to avoid Keratin contamination. Proteins bands were separated by gel electrophoresis at 100 V, constant for 1.5h. The marker and protein bands were fixed and stained with freshly prepared Coomassie blue (G250). All stains and destains were carried out with filtered solutions and all operation was done in a laminar flow hood again to avoid keratin contamination. Each stained band (as visualized under a white light box) was sliced (1 mm^3^ per well (in 96 well MS/MS reaction plate (Intavis AG) in duplicates and sent out to USDA lab, Richmond, CA for further analysis. They were digested and analyzed by mass spectrometry (With Scaffold-2 viewer).

### Generation of MIF expression constructs

The open reading frame of human wildtype MIF was amplified by PCR from the vector pCMV6-entry MIF (Origene) and ligated into the pET11a vector (Novagen) via NdeI/BamHI using the pET11a MIF-fw and pET11a MIF-rev oligonucleotides allowing the tag-free expression in *E.coli*. The C80W-mutation was inserted into pET11a-hMIF using the Quik change site directed mutagenesis kit (Stratagene) and the MIF-C80W_fw and MIF-C80W_rev oligonucleotides.

### Expression and purification of recombinant MIF

For expression of recombinant human wildtype-MIF BL21-(DE3)-Rosetta-pLysS bacteria (Novagen) were transformed with pET11a-MIF or pET11a-MIF-C80W and spread on LB-Agar plates containing 50 µg/mL carbenicillin (Duchefa) and 34 µg/mL chloramphenicol (Sigma). One colony was used to inoculate a preculture of either 10 mL of 2-YT consisting of 16g/L tryptone (Serva), 10 g/L yeast extract (Sigma) and 5g/L NaCl or 10 mL M9 medium (plus ^13^C-labeled glucose and ^15^N-labeled NH_4_Cl) for expression of ^13^C-and ^15^N-labeled MIF, overnight at 37°C. At the next day 2x 500 mL 2-YT or 2 x 500 mL M9 medium containing ^15^N-labeled NH_4_Cl and ^13^C labeled glucose were inoculated with the preculture. The bacteria were grown to an OD_600_ of 0.9 at 37°C before expression of MIF was induced by adding 1 mM IPTG (isopropyl b-D-thiogalactopyranoside; Applichem). The protein was expressed overnight at 18°C. At the next day the bacteria were harvested by centrifugation, resuspended in 20 mM NaPi pH7.0 and lysed by sonication. The lysate was cleared by centrifugation at 20,000g for 30 min and the supernatant was filtered through a 0.45 µm filter (VWR). In order to purify MIF, the lysate was first passed over a 5-mL CM-sepharose column (GE Healthcare), and after washing with 10 column volumes (CV) of 20 mM NaPi pH7.0 MIF was eluted with 20 mM NaPi pH8.0. The eluate was then adjusted to pH8.0 and passed over a 5 mL Q-sepharose column (GE Healthcare). The Flow through was then concentrated using Amicon Ultra-4 centrifugal filters with a cut-off of 10kDa. The purity of MIF was validated by SDS-PAGE and aliquots were stored at −80°C.

### NMR structure

The NMR F_1_-[^13^C,^15^N]-filtered-[^1^H,^1^H]-NOESY spectra for structure calculation were recorded on a 700 MHz Bruker Avance Neo spectrometer equipped with cryoprobe, and the ^15^N-HSQC were recorded on a 600 MHz Bruker Avance III spectrometer equipped with cryoprobe. The filtered NOESY were recorded for MIF 1.0 mM and saturation of PAV-174, i.e., 1.0 mM of PAV-174. at 310K to reduce the rotational correlation time of the system, with two different mixing times 60 and 100ms; the free induction decay was measured for 106ms (2048 points) and the indirect dimension was set to 20ms (400 points); the signal acquisition was performed with 160 scans and the interscan delay was set to 1.5s. The ^15^N-HSQC were measured at 298K with 121ms (2048 points) in the direct dimension and 53 ms (256 points) in the indirect dimension; the signal acquisition was performed through the accumulation of 16 scans and an interscan delay of 1s was applied. The cross-peaks corresponding to ligand-proteins were used to derive ligand-protein distant restraints using the initial rates of the normalized NOE intensity build-up curves (Strotz D et al., 2017). The intermolecular distance restraints involved four different protein aromatic signals and only one protein methyl, consistently with the binding site suggested by the ^15^N-HSQC spectra, which is highly populated with aromatic residues. The intermolecular distance restraints were kept semiambiguous, i.e., only the ligand signals were assigned, and the MIF-PAV-174 complex was calculated with the NMR^2^ software (Orts J et al., 2016) using the CYANA software for tortion angle dynamics structure calculation.

A series of eight ^15^N-T_1_-HSQC with varying T_1_ delays of 0, 40, 70, 110, 150, 200, 250, and 300 ms were acquired. The ^15^N-T_1_-HSQC relaxation experiments were measured at 298K with 121ms (2048 points) in the direct dimension and 53ms (256 points) in the indirect dimension; the signal acquisition was performed through the accumulation of 32 scans and an interscan delay of 1.2 s was applied.

A series of eight ^15^N-T_1_ρ-HSQC with varying T_1_ρ delays of 5, 10, 20, 40, 60, 80, 100, and 120 ms. The ^15^N-T_1_-HSQC relaxation experiments were measured at 298K with 121ms (2048 points) in the direct dimension and 53ms (256 points) in the indirect dimension; the signal acquisition was performed through the accumulation of 64 scans and an interscan delay of 1.5 s was applied.

The relaxation rates were obtained by fitting the signal decays with the software Relax.

### SEC-HPLC-MALS

The samples for SEC-HPLC-MALS were 100 µM of MIF (WT or mutant C80W) in 20 mM phosphate buffer, with or without compound PAV-174 (100 µM).

The HPLC-MALS was performed at 0.5 mL/min and 25°C with an Agilent 1200 series HPLC system and a size exclusion column TSKgel G2000SWXL from Tosoh bioscience. The detection was performed recording the UV absorbance at 280 nm with the HPLC detector and the light scattering with a miniDawn Treos spectrometer from Wyatt. The molecular weight of the detected species is calculated using the extinction coefficients at 280 nm (12950 M^−1^ cm^−1^ for WT and 18450 M^−1^ cm^−1^ for C80W), absorbance at 280 nm and the Rayleigh ratio using the built-in function in Astra software.

### Animal experiments

TgTau58/2 mice (van Eersel et al., 2015) were a kind gift from Novartis (Boston, USA). Two months old males, homozygous for the tau-P301S transgene (n = 12 per group), were either treated with 10% DMSO or 5 mg / kg body weight compound PAV-617 for four weeks administering 100 µL volumes intraperitoneally three times a week, after which animals were perfused with PBS and terminated. The brains were extracted cut into two halves and flash frozen in liquid N_2_. One brain halve was homogenized in cold RAB Hi-Salt buffer (0.1 M MES pH7.0, 1mM EGTA, 0.5 mM MgSO4, 0.75M NaCl, 0.1 mM EDTA, 1x Protease inhibitor cocktail, 1x phosphatase inhibitor cocktail 2) to 10%. For screening 1% NP40, 0.1mg//mL Dnase1 and 5 mM MgSO_4_ were added to the homogenates and after 30 min incubation on ice the protein concentration was determined using the DC Protein Assay Kit (Bio-Rad). Equal amounts of protein were then either directly used for detection of tau in the whole homogenate by immunoblot or sarkosyl insoluble tau was precipitated following a protocol of (Schaeffer V et al., 2012). Succrose was added to 200 µL homogenates, containing 750 µg protein, yielding a final concentration of 10%. After centrifugation at 6,000g for 20 min at 4°C the supernatant was carefully taken and sarkosyl was added to 1% end concentration. The samples were spun down at 100,000g for 1h at 4°C. The supernatants were removed and the pellets carefully washed with 200 µL RAB buffer without sucrose. The pellets were then incubated in 60 µL PBS with 8 M urea at room temperature overnight. 20 µL were used for immunoblot analysis.

## QUANTIFICATION AND STATISTICAL ANALYSIS

All statistical analyses were performed as indicated using GraphPad Prism (Versions 6 & 9.4); GraphPad Software Inc., San Diego, CA, USA). Data are presented as mean +/− SEM and appropriate statistical tests and p-values are stated in the respective figure legends. Outlier were checked and removed by ROUT analysis (Q = 1%). p-values of *p ⩽ 0.05, **p ⩽ 0.01, ***p ⩽ 0.001, ****p ⩽ 0.0001 were used as significance levels.

