## Supplemental Figures for "Oxidized MIF is an Alzheimer’s Disease drug target relaying external risk factors to tau pathology"

### Supplementary Figures

**Supplementary Figure 1 (referring to table 1)**

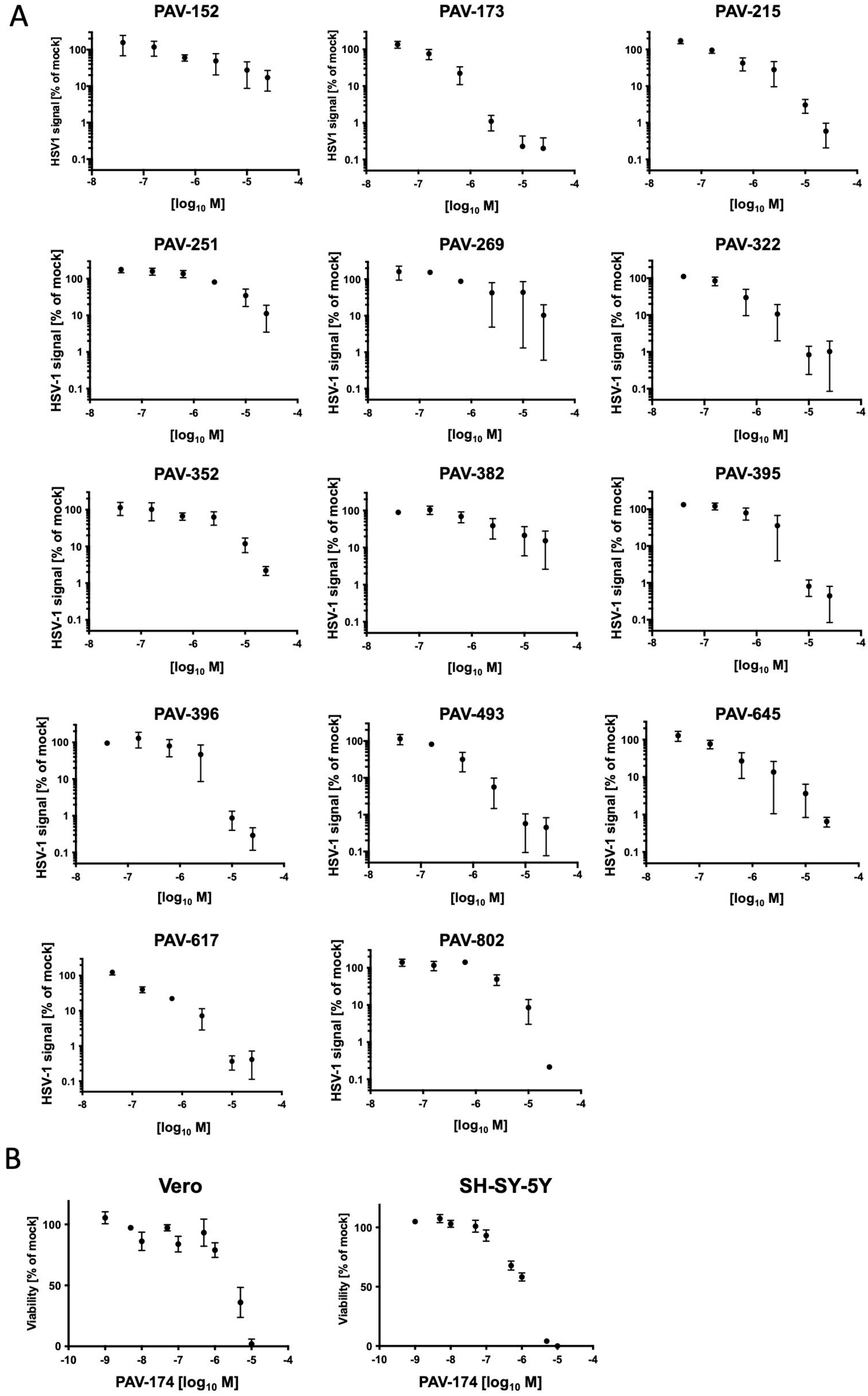

**Supplementary Figure 1 (related to Table 1)**

(A) The $IC_{50}$ of analogs of PAV-174 were determined by in-Cell ELISA after Vero cells were treated with increasing concentrations of the compounds and infected with HSV-1 (MOI=1) for 20h. Each data point displays the mean +/- SEM of three independent experiments (n=3). $IC_{50}$ values were calculated using the logarithm of inhibitor concentration on GraphPad Prism 6.0.

(B) MTT assay of PAV-174 in Vero (left) or SH-SY5Y-tau-P301S (right) cells treated with increasing concentrations of compound for 24h (Vero) or 48h (SH-SY5Y) resulted in a $CC_{50}$ of 1.12 μM (Vero) or 1.48 μM (SH-SY5Y). Each data point displays the mean +/- SEM of three (Vero) or four (SH-SY5Y) experiments (n=3 or 4). $CC_{50}$ values were calculated using the logarithm of inhibitor concentration on GraphPad Prism 6.0.

**Supplementary Figure 2 (referring to Figure 2)**

A

| Protein name | MW (kDa) | Accession number | Peptides identified | Mascot ion score | % coverage |
|---|---|---|---|---|---|
| MIF | 12 | MIF_PIG | (M)PMFVVNTNVPR(A) | 51.5 | 17 |
| | | | (K)LLCGLLAER(L) | 60,9 | |
| cutA | 19 | F1RZR6_PIG | (K)GKIEEDSEVLMMIK(T) | 37,3 | 16 |
| | | | (K)IEEDSEVLMMIK(T) | 40,8 | |
| | | | (K)TQSSLVPALTDFVR(S) | 58,7 | |
| GSTP1 | 23 | GSTP1_PIG | (-)PPYTITYFPVR(G) | 56,5 | 27 |
| | | | (K)FQDGDLTLYQSNAILR(H) | 84,8 | |
| | | | (K)EAALVDMVNDGVEDLR(C) | 130,1 | |
| | | | (K)YATLIYTNYEAGK(E) | 41,4 | |
| PRDX2 | 22 | F1SDX9_PIG | (R)NLSLDYGVLKEDEGIAYR(G) | 55,5 | 15 |
| | | | (R)QITVNDLPVGR(S) | 57,2 | |
| PKM2 | 57 | F1SHL7_PIG | (R)NTGIICTIGPASR(S) | 56,5 | 7 |
| | | | (K)VYVDDGLISLLVK(Q) | 61,2 | |
| | | | (R)GDLGIEIPAEK(V) | 52,5 | |
| CALML3 | 12 | F1SQH2_PIG | (K)EAFSLFDK(D) | 59,6 | 42 |
| | | | (K)EAFSLFDKDGDGTITTK(E) | 73,4 | |
| | | | (K)DTDSEEEIREAFR(V) | 46,0 | |
| | | | (R)VFDKDGNGYISAAELR(H) | 77,3 | |
| | | | (R)VFDKDGNGYISAAELR(H) | 69,4 | |
| | | | (K)DGNGYISAAELR(H) | 63,7 | |
| HSPCB | 83 | F1RQU2_PIG | (R)TLTLVDTGIGMTK(A) | 70,3 | 5,4 |
| | | | (K)ADLINNLGTIAK(S) | 80,6 | |
| | | | (R)GVVDSEDLPLNISR(E) | 69,8 | |

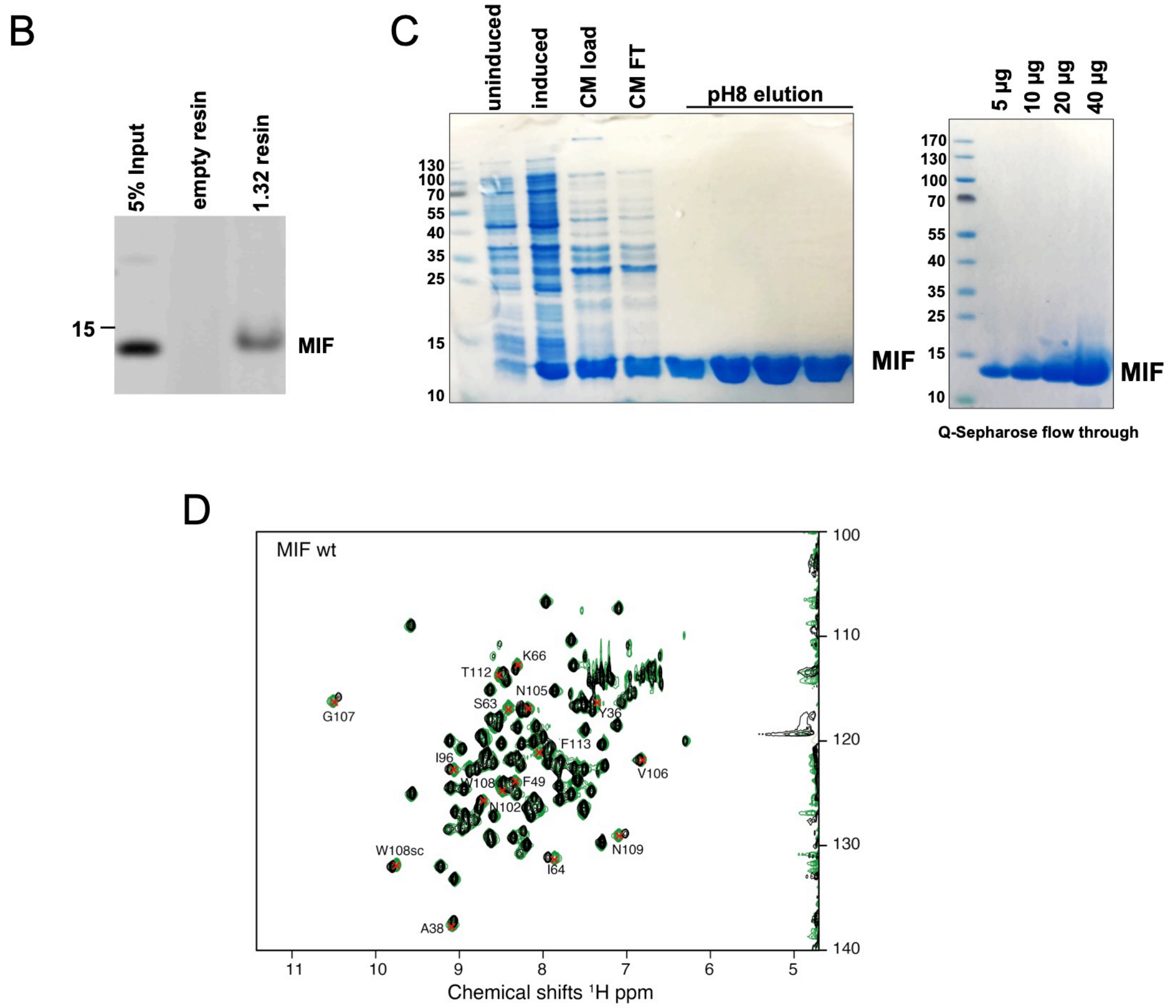

**Supplementary Figure 2 (related to Figure 2)**

(A) List of peptides identified by DRAC analysis including Mascot ion score.

(B) DRAC assay with cell lysates (500 µg) derived from SH-SY5Y-tau-P301S cells that were applied either on empty control resin or resin presenting PAV-645. MIF was eluted from drug resins by urea. On the left 5% of input material is shown.

(C) Recombinant expression and purification of human wildtype MIF in *E.coli*. Expression of tag-free MIF was induced in BL21 bacteria and MIF was then purified by ion exchange chromatography (CM Sepharose). The purification steps are shown in the SDS-PAGE on the left. The right image shows the purity (>95%) of concentrated MIF after polishing via Q-sepharose.

(D) $^{15}$N-HSQC spectra of the apo-MIF (green) and of the MIF in the presence of PAV-174 (black). The resonances showing the highest perturbations are zoomed in within the squares in Figure 2A. The residues showing the most important chemical shift perturbations are shown in the MIF structure in the Figure 2A.

**Supplementary Figure 3 (referring to Figure 3)**

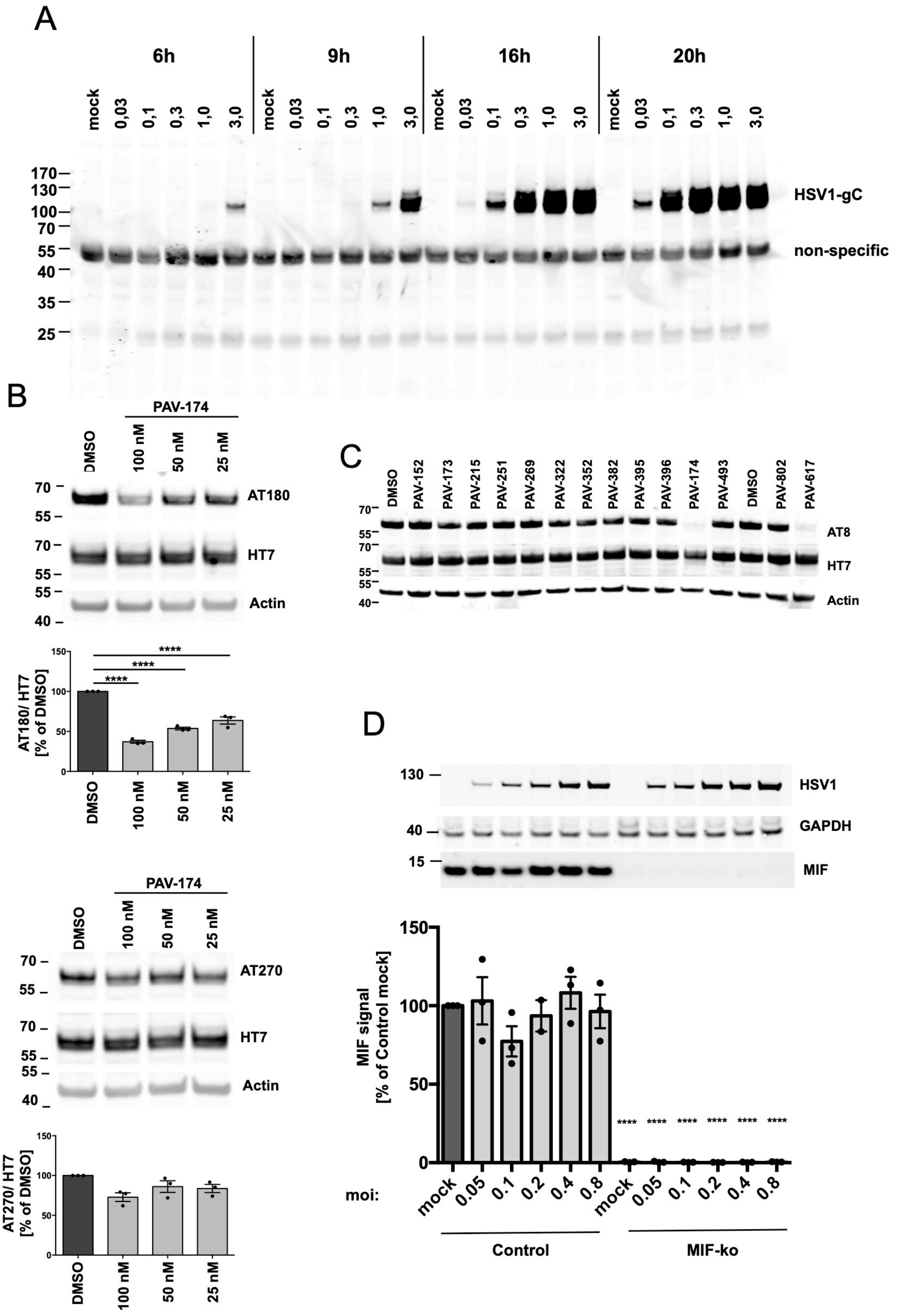

**Supplementary Figure 3 (related to Figure 3)**

(A) SH-SY5Y-tau-P301S cells could efficiently be infected with HSV-1. The cells were infected with increasing MOIs (0.03 to 3.0) and then lysed after 6h, 9h, 16h or 24h. The lysates were analyzed by Western Blot using an antibody against the glycoprotein C (gC) of HSV-1.

(B) PAV-174 selectively reduced tau phosphorylation. SH-SY5Y-tau-P301S cells were treated with PAV-174 for 48h. Lysates were analyzed with phospho-tau specific antibodies by Western Blot. Actin served as loading control. PAV-174 efficiently reduced tau phosphorylation at T231 recognized by AT180 (top) but only at high concentrations at T181 recognized by AT270 (below). The diagrams show the signals of the phospho-tau specific antibodies normalized to total tau (HT7) from three (n=3) independent experiments. Data were analyzed by One-way ANOVA (Dunnet´s post-hoc).

(C) Representative Western Blot of results shown in Fig. 3C.

(D) Expression of MIF was not modulated upon infection with HSV-1. SH-SY5Y-tau-P301S-CRISPR-control and SH-SY5Y-tau P301S-MIF-ko cells were infected with increasing MOIs of HSV-1 for 16h and lysates were analyzed for MIF expression. The expression of HSV-1 antigen is shown on top and GAPDH served as loading control. The diagram displays the quantification of MIF normalized to GAPDH derived from three independent experiments (n=3). Data were analyzed by Two-way ANOVA (Sidak´s post-hoc).

### Supplementary Figure 4 (referring to Figure 3)

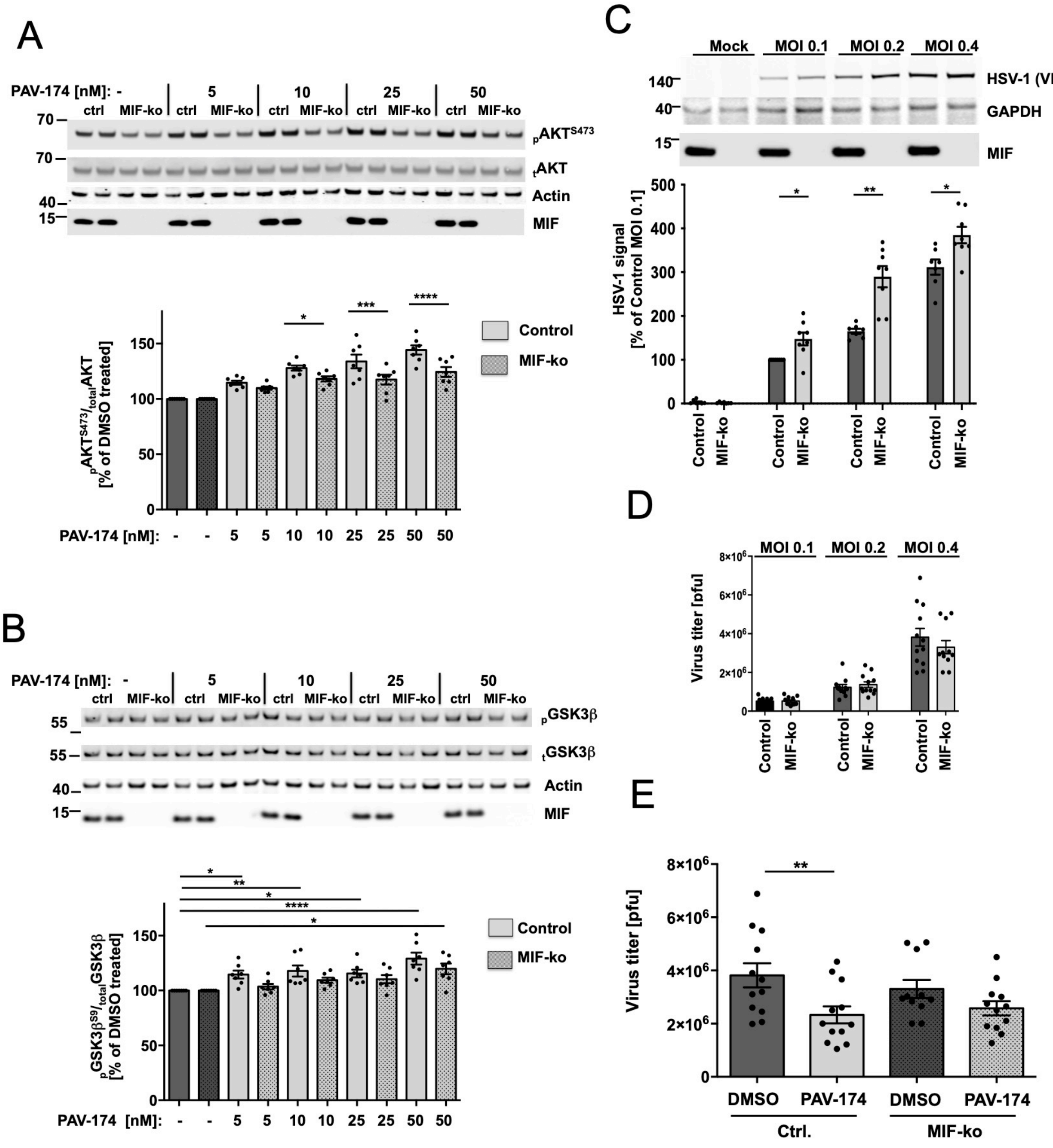

**Supplementary Figure 4 (related to Figure 3)**

(A/B) PAV-174 induces phosphorylation of Akt at $S^{473}$ (A) and GSK3β at $S^{9}$ (B) in a MIF dependent manner. SH-tauP301S-CRISPR-control or -MIF-ko cells were treated with increasing concentrations of PAV-174 for 6h. PAV-174 led to a dose dependent and significantly higher phosphorylation of $Akt^{473}$ in SH-SY5Y-tau-P301S-control than in -MIF-ko cells. Similar PAV-174 significantly increased phosphorylation of GSK3β at $Ser^{9}$ already at low concentrations only in SH-tauP301-control but not in -MIF-ko cells. The diagrams show the average values of $pAkt^{S473}$ normalized to total Akt or $pGSK3\beta^{S9}$ normalized to total GSK3β as % of DMSO treated cells of the respective cell line derived from seven independent experiments (n=7). Data were analyzed by Two-way ANOVA (Sidak´s post-hoc).

(C) Increased production of HSV-1 capsid proteins in MIF-ko cells compared to control cells. SH-SY5Y-tau-P301S-CRISPR-control and -MIF-ko cells were infected with the indicated MOIs of HSV-1. The amount of viral capsid proteins within the lysate 16h p.i. were determined by Western Blot using an antibody against the VP5 capsid protein of HSV-1. GAPDH was used as internal control. The absence of MIF in SH-tau-MIF-ko was verified using a polyclonal antibody against MIF. The diagram shows the result from eight infections (n=8). Data were normalized to Control, MOI = 0.1 and analyzed by Two-way ANOVA (Sidak´s post-hoc).

(D) Infection of SH-SY5Ytau-P301S-MIF-ko cells led not to an increased production of HSV-1 infectious particles. SH-SY5Y-tau-P301S control and -MIF-ko cells were infected with the indicated MOIs of HSV-1. The viral titer was determined by plaque assay 16h p.i.. The diagram shows the result from twelve infections (n=12). Data were analyzed by Two-way ANOVA (Sidak´s post-hoc). No significant differences were found between both cell lines.

(E) MIF is a functional target of PAV-174. SH-SY5Y-tau-P301S-CRISPR control and -MIF-ko cells were infected with HSV-1 (MOI 0.4) and either treated with DMSO or PAV-174 (5 nM). Plaque assays were performed 16h p.i. The diagram shows the viral titers from twelve (n=12) infections. Data were analyzed by Two-way ANOVA (Sidak´s post-hoc).

**Supplementary Figure 5**

A

| | PAV-174 | PAV-617 |
|---|---|---|
| MTD [mg/kg] | 0,5 | 20 |
| Bioavailability [ng.h/ mL] | 122 (0.5 mg/ kg) | 3093 (5 mg/kg) |
| Protein binding [%] | 99,9 | 91,2 |

B

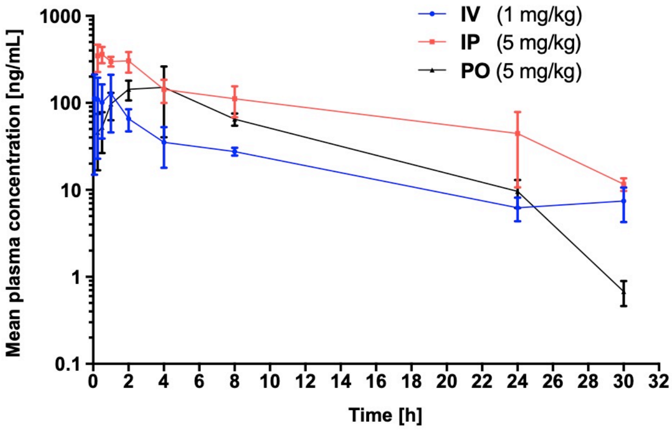


| Route | Dose (mg/kg) | AUC $_{(last)}$ (ng.h/mL) | AUC $_{(0\ to\ inf)}$ (ng.h/mL) | Cmax (ng/ml) | Tmax (h) | Kel (1/h) | $t_{1/2}$ (h) | MRT (h) | CL (mL/min/ kg) | Vz (L/Kg) | % F |
|---|---|---|---|---|---|---|---|---|---|---|---|
| IV | 1 | 747.2 | 831.5 | 128.0 | 1.0 | 0.1 | 7.8 | 7.1 | 20.0 | 12.7 | |
| IP | 5 | 2974.7 | 3093.0 | 362.6 | 0.5 | 0.1 | 7.0 | 8.3 | | | 80 |
| PO | 5 | 1531.8 | 1535.1 | 151.1 | 4.0 | 0.2 | 3.3 | 6.6 | | | 41 |

C

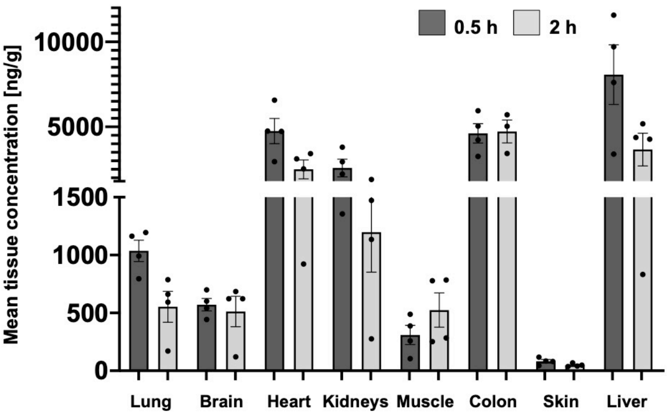


| Tissue | Mean tissue Conc (ng/g) | |
|---|---|---|
| | 0.5 h | 2 h |
| Plasma (ng/mL) | 311.4 ± 8.22 | 287.7 ± 134.0 |
| Lung | 1036.5 ± 184.0 | 553.1 ± 267.3 |
| Brain | 570.8 ± 107.6 | 643.3 ± 36.4 |
| Heart | 4752.1 ± 1476.7 | 2498.1 ± 1114.0 |
| Kidneys | 2580.8 ± 1040.2 | 1197.2 ± 690.0 |
| Muscle | 309.4 ± 166.2 | 524.4 ± 296.7 |
| Colon | 4618.3 ± 1144.1 | 4727.4 ± 233.8 |
| Skin | 82.0 ± 30.1 | 48.5 ± 14.3 |
| Liver | 8070.6 ± 3511.7 | 3664.0 ± 1930.9 |

| Tissue | Tissue/plasma Ratio (ng/g)/(ng/mL) | |
|---|---|---|
| | 0.5 h | 2 h |
| Lung | 3.4 ± 0.7 | 1.9 ± 0.2 |
| Brain | 1.7 ± 0.2 | 1.8 ± 0.1 |
| Heart | 13.3 ± 3.3 | 9.0 ± 1.1 |
| Kidneys | 7.9 ± 4.0 | 4.0 ± 1.1 |
| Muscle | 0.8 ± 0.4 | 2.1 ± 1.0 |
| Colon | 14.3 ± 4.0 | 13.2 ± 2.4 |
| Skin | 0.2 ± 0.1 | 0.2 ± 0.1 |
| Liver | 22.1 ± 10.2 | 12.2 ± 2.8 |

D

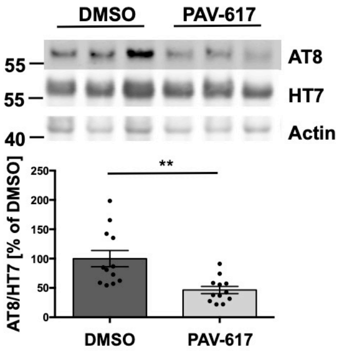


E

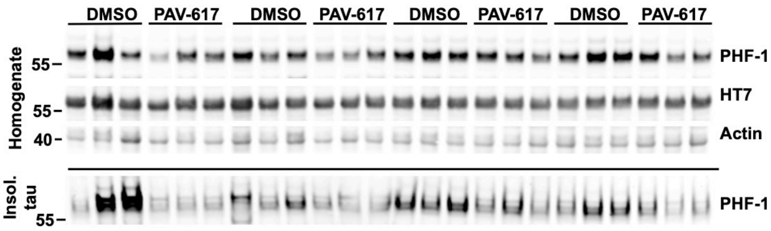

**Supplementary Figure 5 (related to Figure 4)**

(A) MTD (Maximum Tolerated Dose) assessment of PAV compounds in BALB/c mice. BALB/c mice were dosed through Intraperitoneal (IP) route at different doses as per body weight of animals and mice were observed at different time interval for toxic signs till 48 h, PD (Post dose). MTD at was declared at NOAEL (No-observed-adverse-effect level) conc. of PAV compound. PAV-617 showed a markedly reduced protein binding (91.2%) compared to PAV-174 (99.9%).

(B) Pharmacokinetic assessment of PAV-617 in Sprague Dawley rats. Male SD rats (*N=4*) were dosed via three routes namely Intravenous (IV)- 1 mg/kg, Intraperitoneal (IP)- 5 mg/kg and Per Oral (PO)- 5 mg/kg. Plasma concentration was analyzed at different time intervals. Pharmacokinetic parameters were determined using WinNonlin.

Cmax (peak plasma concentration) achieved through IV was 128 ng/mL, IP was 362.6 ng/mL and PO was 151.1 ng/mL at Tmax (time of peak concentration observed) 1 h, 0.5 h and 4 h respectively. Compound PAV-617 has moderate t1/2 (terminal half-life) through IV- 7.8 h, IP-7 h and PO- 3.3 h. CL (steady-state clearance) was found to be moderate- 20 mL/min/Kg which was 2.75 times lower than rat liver blood flow. Vz (volume of distribution) was found to be high- 12.7 L/kg and F (fraction bioavailability) was found to be very good through IP- 80 % and moderate through PO- 41%.

(C) Tissue partitioning of PAV-617 in Sprague Dawley rats. Male SD rats (*N=4*) were dosed via Intraperitoneal (IP)-5 mg/kg. Plasma and major organs such as lung, brain, heart, kidney, muscle, colon, skin, and liver were collected at 0.5 h and 2 h post dosing. Partitioning of PAV-617 by each organ was analyzed.

(D) Reduction of tau phosphorylation was observed *in vivo* after treating tau58/2 mice with 5 mg/kg of PAV-617, a structural analog of PAV-174. Phosphorylated tau (AT8) was reduced in the homogenates of the treated mice. The diagrams show the average signals of 12 mice per treatment group (n = 12) derived from two independent Western Blots. Data were analyzed by unpaired two-tailed t-test.

(E) Representative blots of results described in Figure 5B/C.

### Supplementary Figure 6

A

| sample | sex | age | braak | amyloid | pmd [h] | ph | weight | apoe |
|---|---|---|---|---|---|---|---|---|
| AD 1 | f | 72 | 6 | C | 5,92 | 6,54 | 960 | 32 |
| AD 2 | f | 68 | 6 | C | 3,83 | 6,50 | 985 | 32 |
| AD 3 | f | 84 | 6 | C | 4,83 | 6,06 | 960 | 33 |
| AD 4 | f | 86 | 6 | C | 5,25 | 6,20 | 1090 | 33 |
| AD 5 | m | 87 | 5 | C | 6,17 | 6,14 | 1047 | 33 |
| AD 6 | f | 90 | 6 | C | 4,75 | 6,39 | 1045 | 33 |
| AD 7 | m | 71 | 6 | C | 6,58 | 6,20 | 1092 | 33 |
| AD 8 | m | 70 | 6 | C | 6,33 | 6,40 | 1261 | 43 |
| AD 9 | f | 89 | 6 | C | 4 | 6,40 | 977 | 43 |
| AD 10 | m | 81 | 6 | C | 7,83 | 6,15 | 1120 | 43 |
| N 1 | f | 60 | 0 | O | 8,17 | 6,58 | 1310 | 32 |
| N 2 | f | 75 | 1 | A | 9,17 | 6,57 | 1305 | 32 |
| N 3 | f | 72 | 1 | A | 6,83 | 7,22 | 1165 | 33 |
| N 4 | f | 78 | 1 | A | 7,17 | 6,32 | 1120 | 33 |
| N 5 | m | 90 | 2 | O | 5,75 | 6,35 | 1215 | 33 |
| N 6 | m | 73 | 2 | A | 8 | 5,37 | 1553 | 33 |
| N 7 | f | 88 | 3 | A | 5,42 | 6,33 | 1010 | 33 |
| N 8 | f | 91 | 3 | A | 4,17 | 6,58 | 1052 | 43 |
| N 9 | m | 83 | 1 | B | 5,75 | 6,35 | 1590 | 43 |
| N 10 | m | 102 | 3 | A | 5 | 6,64 | 1124 | 43 |

B

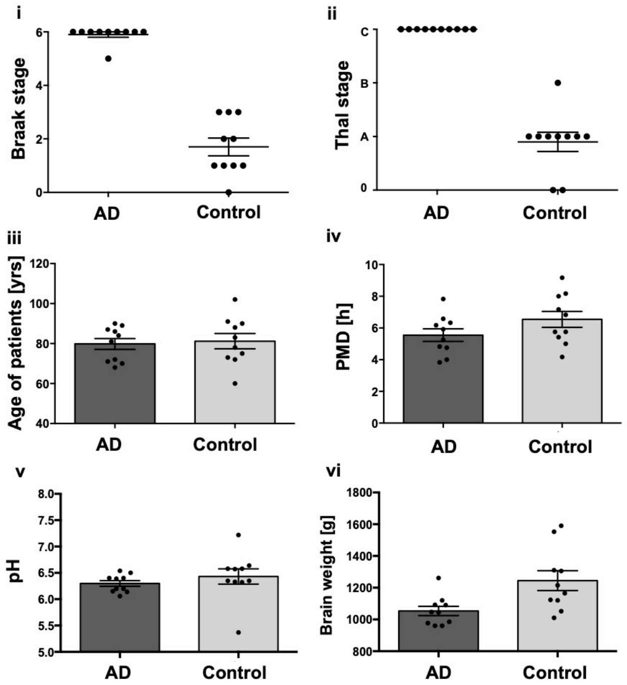


C

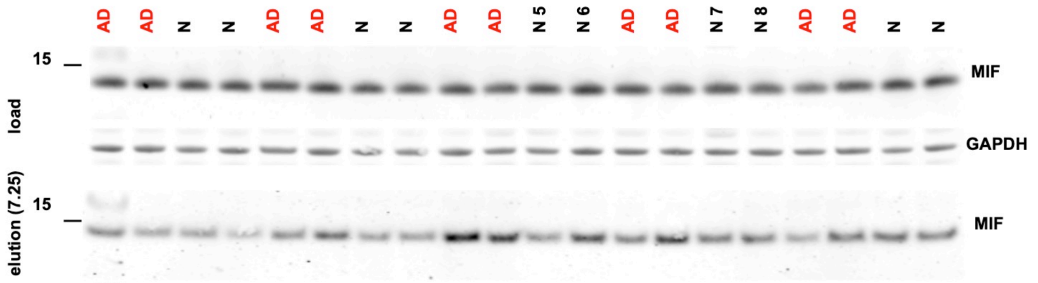


D

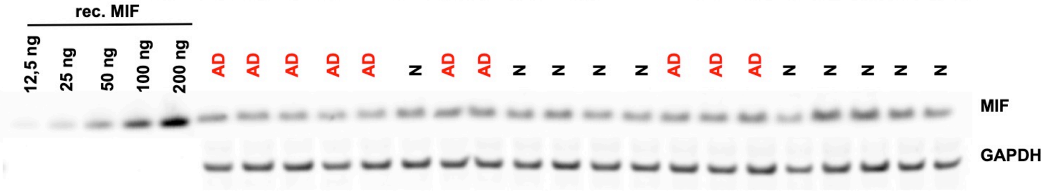


E

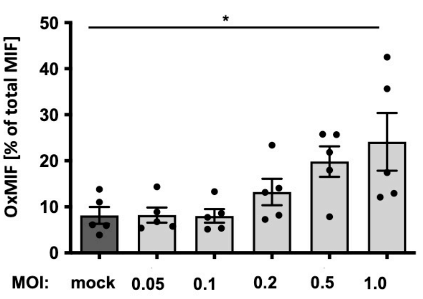


F

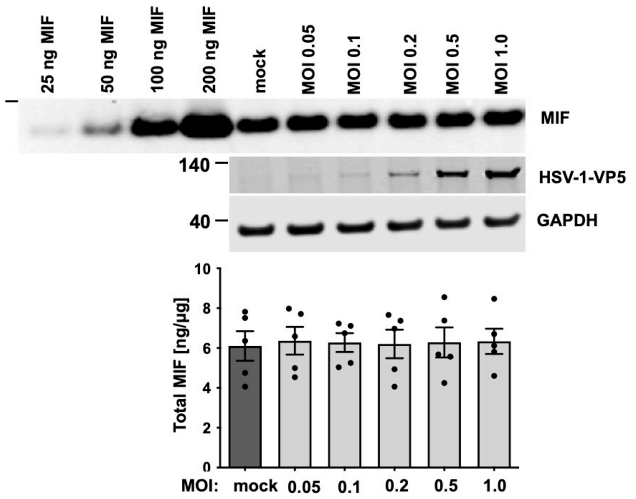

**Supplementary Figure 6 (related to Figure 5)**

(A) Table describing the brain samples retrieved from the The Netherlands Brain Bank regarding, sex, age, tau pathology (Braak stages), amyloid pathology (according to Thal staging), *post mortem* time (pmd), pH of sample, brain weight and ApoE gene status.

(B) All AD samples displayed severe tau (i) as well as amyloid pathology (ii) and show an equal distribution of age (iii) and no differences in *post mor*tem delay (pmd) (iv) or pH (v). The weight of the brains from the AD patients was significantly reduced compared to the controls (vi). Data of 10 samples per group were analyzed by unpaired two-tailed t-test.

(C) Representative complete Western Blot of DRAC analysis of brain samples shown in Fig. 5A). The upper panel displays the loading controls (MIF and GAPDH) of brain homogenates and the lower panel the precipitated and PAV-174-eluted MIF.

(D) Representative quantitative Western Blot of brain samples used for normalizing the oxMIF concentrations determined by the sandwich ELISA shown in Fig. 5C. Increasing concentrations (12,5 ng to 200 ng) of recombinant expressed human wt-MIF were used to generate a standard curve allowing the quantification of MIF within the brain samples. MIF signals were normalized to GAPDH.

(E). Induction of oxMIF in differentiated LUHMES cells upon infection with HSV-1. Differentiated LUHMES cells were infected with increasing MOIs of HSV-1. Cells were lysed 16h p.i and the amount of oxMIF was measured by sandwich ELISA. The results were normalized to total MIF as determined by Western Blot (F). The diagram shows the results from five independent infections (n=5). Data were analyzed by One-way ANOVA (Dunnet´s post-hoc).

(F) Quantification of total MIF in lysates of differentiated LUHMES cells. Different amounts of recombinant MIF were used to generate a standard curve. MIF signals were normalized to GAPDH. The infection of LUHMES cells with HSV-1 was demonstrated with an antibody against a capsid protein of HSV-1 (VP5). The diagram shows the results from five independent infections (n=5). Data were analyzed by One-way ANOVA (Dunnet´s post-hoc).

### Supplementary Figure 7

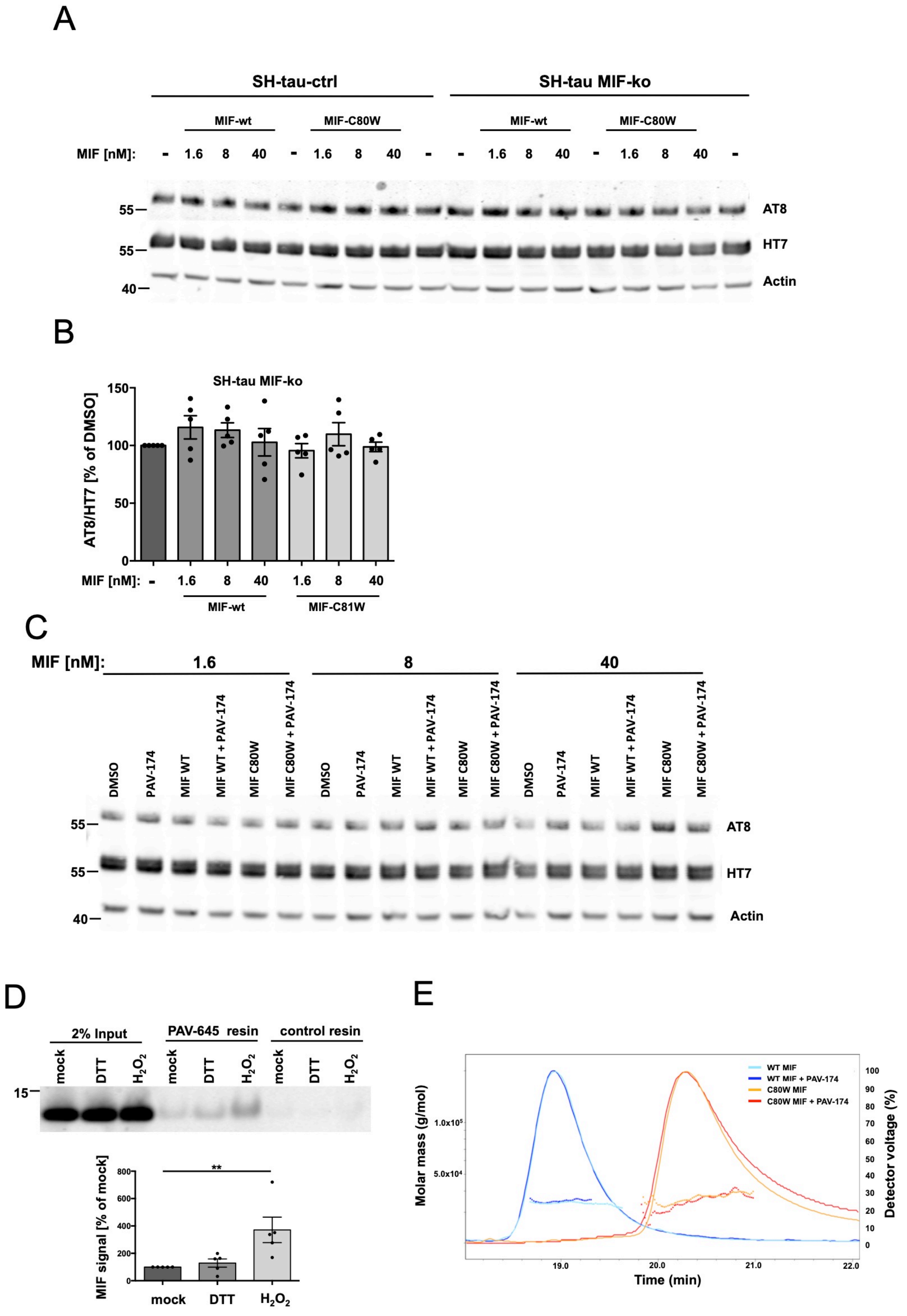

**Supplementary Figure 7 (related to Figure 6)**

(A) Representative complete Western Blot of the analysis presented in Figure 6A. The upper panel displays AT8 and below total tau as detected by HT7 is shown. Actin served as loading control.

(B) Exogenous applied oxMIF did not induce tau phosphorylation in SH-SY5Y-tauP301S–MIF-ko cells. The cells were treated with recombinant wt-MIF or MIF-C80W for 6h. The diagram shows the average values of five independent experiments (n=5). Data were analyzed by one-way ANOVA (Tukey´s post-hoc).

(C) Representative complete Western Blot of the analysis presented in Figure 6B. The upper panel displays AT8 and below total tau as detected by HT7 is shown. Actin served as loading control.

(D) DRAC analysis with reduced (6 mM DTT) or oxidized (6 mM $H_2O_2$) wt-MIF. Significantly more MIF bound to the PAV-645 resin when MIF was oxidized with $H_2O_2$. The diagram shows the signal of eluted MIF normalized to non-treated wt-MIF derived from five (n=5) pull-downs. Data were analyzed by One-way ANOVA (Dunnett´s post-hoc).

(E) Chromatograms of the different sample injections detected with MALS (solid lines, detector voltage). The molar mass (squares) is calculated from the Rayleigh ratio (derived by Astra from the detector voltage) corrected by the UV absorption at 280 nm. The different samples are all concentrated at 100 µM of MIF (WT or C80W) +/- 100 µM of PAV-174 compound. The sample C80W MIF was injected twice to facilitate the calculation molar mass.
